# LIQ HD (Lick Instance Quantifier Home cage Device): An open-source tool for recording undisturbed two-bottle drinking behavior in a home cage environment

**DOI:** 10.1101/2022.12.16.520661

**Authors:** Nicholas Petersen, Danielle N. Adank, Ritika Raghavan, Danny G. Winder, Marie A. Doyle

## Abstract

Investigation of rodent drinking behavior has provided insight into drivers of thirst, circadian rhythms, anhedonia, and drug and ethanol consumption. Traditional methods of recording fluid intake involve weighing bottles, which is cumbersome and lacks temporal resolution. Several open-source devices have been designed to improve drink monitoring, particularly for two-bottle choice tasks. However, recent designs are limited by the use of infrared photobeam sensors and incompatibility with prolonged undisturbed use in ventilated home cages. Beam-break sensors lack accuracy for bout microstructure analysis and are prone to damage from rodents. Thus, we designed LIQ HD (Lick Instance Quantifier Home cage Device) with the goal of utilizing capacitive sensors to increase accuracy and analyze lick microstructure, building a device compatible with ventilated home cages, increasing scale with prolonged undisturbed recordings, and creating a design that is easy to build and use with an intuitive touchscreen graphical user interface. The system tracks two-bottle choice licking behavior in up to 18 rodent cages, or 36 single bottles, on a minute-to-minute timescale controlled by a single Arduino microcontroller. The data are logged to a single SD card, allowing for efficient downstream analysis. With sucrose, quinine, and ethanol two-bottle choice tasks, we validated that LIQ HD has superior accuracy compared to photobeam sensors. The system measures preference over time and changes in bout microstructure, with undisturbed recordings lasting up to 7 days. All designs and software are open-source to allow other researchers to build upon the system and adapt LIQ HD to their animal home cages.

**Significance Statement:** Two-bottle choice drinking tasks are traditionally performed by periodically weighing bottles, which is cumbersome and lacks temporal resolution. Several open-source tools have been developed to improve drink monitoring in various settings. However, no open-source devices have been designed specifically to investigate temporally precise two-bottle choice drinking behavior and bout microstructure during prolonged undisturbed tasks in mouse ventilated home cages at a large scale. Our design, LIQ HD (Lick Instance Quantifier Home cage Device), is a home cage compatible system that utilizes capacitive sensors for highly accurate lick detection during two-bottle choice tasks in up to 18 cages driven by a single Arduino microcontroller. The system is low-cost, easy to build, and controlled via touchscreen with an intuitive graphical user interface.

## Introduction

Monitoring of fluid intake and drinking behaviors is a powerful toolset in neuroscience research. These data provide insight into maladaptive behaviors observed in a wide range of disorders, such as obesity, substance use, depression, and others. A rich literature describes key brain regions in which populations of neurons are drivers of thirst, anhedonia, circadian rhythms of fluid consumption, and drug and ethanol consumption. Typical examples of such studies utilize standard voluntary two-bottle choice tasks in which animals are provided bottles in the home cage, one containing water and the other an experimental solution. Measures of fluid intake and preference are then calculated from bottle weight measurements manually taken by experimenters.

While two-bottle choice tasks remain the most common method for studying voluntary intake in rodents, performing the task manually is cumbersome, as data are traditionally collected by taking weight measurements throughout a specific time period, usually 1-3 days. Although this technique provides valuable information to researchers, it lacks temporal resolution. Increasing the frequency of bottle weighing increases variability and adds additional stress, as the animals must be disturbed to collect the data. Commercially available systems can track drinking behavior in a more automatized fashion in a home cage environment (Mingrone et al., 2020). These automated home cage monitoring systems are valuable because they generate data from a substantial number of different behavioral and metabolic measurements and drinking behavior. However, they are costly, require trained personnel, limit the number of cages that can be used, and are only available at a limited number of research institutions. Similarly, operant conditioning chambers can be used for assaying motivated behaviors related to fluid intake and tracking fine details associated with these behaviors, including lick microstructures; however, these tasks are not performed in a home cage environment and similarly require additional equipment and specialized training.

Caveats like those described above have inspired groups to develop open-source tools to study rodent drinking behaviors. The technology to detect licks (“lickometer”) or drink events for these devices has generally fallen into three categories: electrical lick sensors, optical lick sensors, and force lick sensors (Ulman et al., 2008; Weijnen, 1998). Early lickometer designs, although accurate, were hindered by a lack of easy-to-use commercially available components (Dole et al., 1983; Mundl and Malmo, 1979) or sub-optimal compatibility with the home cage environment (Schoenbaum et al., 2001). Newer systems typically involve the use of inexpensive, commercially available sensors and components controlled by either a standard computer or microcontrollers, such as an Arduino or Raspberry Pi. Recently, two versions of infrared (IR) photobeam-break sensor based systems have been a common choice for detecting rodent drink events in a home cage environment (Frie and Khokhar, 2019; Godynyuk et al., 2019). The devices generated by these groups have filled many user needs, chiefly generating easy-to-use open-source designs that greatly improved the temporal resolution of drinking data. Godynyuk et al. created a mouse system optimized for use with *in vivo* recordings, such as fiber photometry; however, this system is not compatible with ventilated home cage systems and only has a capacity of 15 mL per bottle, which would require significant investigator work to refill bottles for chronic fluid measurements. Frie et al. built off this first system by adapting it for rat cages and adding a capacitance-sensing eTape to monitor changes in fluid levels within each water bottle. Again, while this design includes an innovative eTape-based measure of fluid levels, it is not compatible with ventilated mouse home cages due to its size. Finally, as the IR sensors are triggered by the animal’s snout in addition to its tongue, they detect drinking events rather than individual licks, thus making all IR beam break-based systems incompatible for the analysis of bout microstructure.

The use of electrical- or capacitive-sensing has shown to have superior accuracy in detecting licks (Longley et al., 2017; Melo et al., 2022; Parkison et al., 2012). However, previously developed electrical-sensing devices are not designed to be compatible with a home cage because they require the use of a metal floor plate in a custom-built enclosure (Melo et al., 2022; Raymond et al., 2018). Devices that utilize a capacitive sensor on a chip do not require a metal place, providing greater design flexibility. Capacative sensors have allowed groups to design systems optimized for detecting licks in combination with recording movement (Parkison et al., 2012) and rat home cage operant devices (Longley et al., 2017). However, current designs using this technology are limited by their incompatibility with ventilated mouse home cages within a typical animal vivarium (Parkison et al., 2012) or have limited scalability (one device per microcontroller) (Longley et al., 2017), which is a common issue across most devices (Godynyuk et al., 2019; Raymond et al., 2018). Thus, our goal was to design a device that utilizes capacitive sensing on a chip for accuracy and consistency, is compatible with a proper undisturbed mouse home cage environment and is scalable to record from multiple cages from a single microcontroller. In addition, we sought for the device to be used undisturbed for more extended periods of time (hold more fluid volume), have in-cage sensing components that are resistant to rodent destruction, and be intuitive and easy to build and use.

Here we present LIQ HD (Lick Instance Quantifier Home cage Device): an affordable, intuitive, and easy-to-build device that utilizes capacitive sensor technology to track two-bottle choice drinking behavior in up to 18 rodent home cages, or 36 single bottles, on a minute-to-minute timescale running off a single Arduino microcontroller. The system is built with 3D-printed parts and affordable commercially available electronics. Unlike most currently available open-source systems, our device is designed to be implemented directly in the animal’s home cage while on ventilated animal facility racks without any cage modification or requiring special housing conditions. The data for all cages are logged to a single SD card, allowing for efficient downstream analysis. Additionally, the system features a touchscreen controller with an intuitive graphical user interface to prevent the need for any code modification between experiments. Licks captured with LIQ HD strongly correlate with the volume consumed and has been tested in our hands with continued use over several months, with undisturbed runs for up to 7 days. Within the minute-by-minute data, in addition to lick number and lick duration, researchers can track drink preference as well as changes to the animals’ bout microstructure (bout duration, bout size, lick frequency, inter-lick interval) over time. It is our goal that LIQ HD will provide researchers with the tools necessary to gather fine-tuned drinking behavior data and streamline two-bottle choice paradigms, particularly those involving long-term home cage monitoring, such as ethanol two-bottle choice.

## Materials and Methods

### LIQ HD Build Instructions

LIQ HD is built from commercially available electronic components combined with 3D-printed parts. The system controls 3 separate 12-channel MPR121 capacitive sensor breakout boards (Adafruit) for a total of 18 individual devices (with 2 sippers each) and runs off a single Arduino Mega microcontroller (**Figure 1A,B**). This capacitance-based sensor determines touch events by calculating a change in capacitance, or the ability of an object or substance to store an electric charge. This method does not require for a circuit to be completed for touch detection, thus there is no need for a conductive plate in the animal cage, and normal bedding can be utilized. Each metal sipper is in contact with conductive copper foil tape that is wired to an MPR121 capacitance sensor breakout board. With only the very tip of the sipper exposed, the device can sense individual licks without interference from other parts of the animal. While we have not done so, the Arduino code can be modified to support up to 4 separate MPR121 boards for a total of 24 devices (48 sippers). Date and time are kept with the Adafruit Data Logger Shield, which keeps time even if the device is unplugged or reset. The Data Logger Shield also writes the data to a single file on an SD card for efficient downstream analysis. Finally, a 2.8” Adafruit Touchscreen Shield is used to display an easy-to-use graphical user interface. This allows for the user to change device settings (time, light/dark schedule, sensor sensitivity, metrics to record), to mount/eject the SD card, and to start/stop/pause recordings. The screen also displays the total number of licks for each sipper during recordings (which can be refreshed to the user’s convenience) and will prompt the user if there are any errors, such as disconnected sensors or SD card.

**Figure 1.**
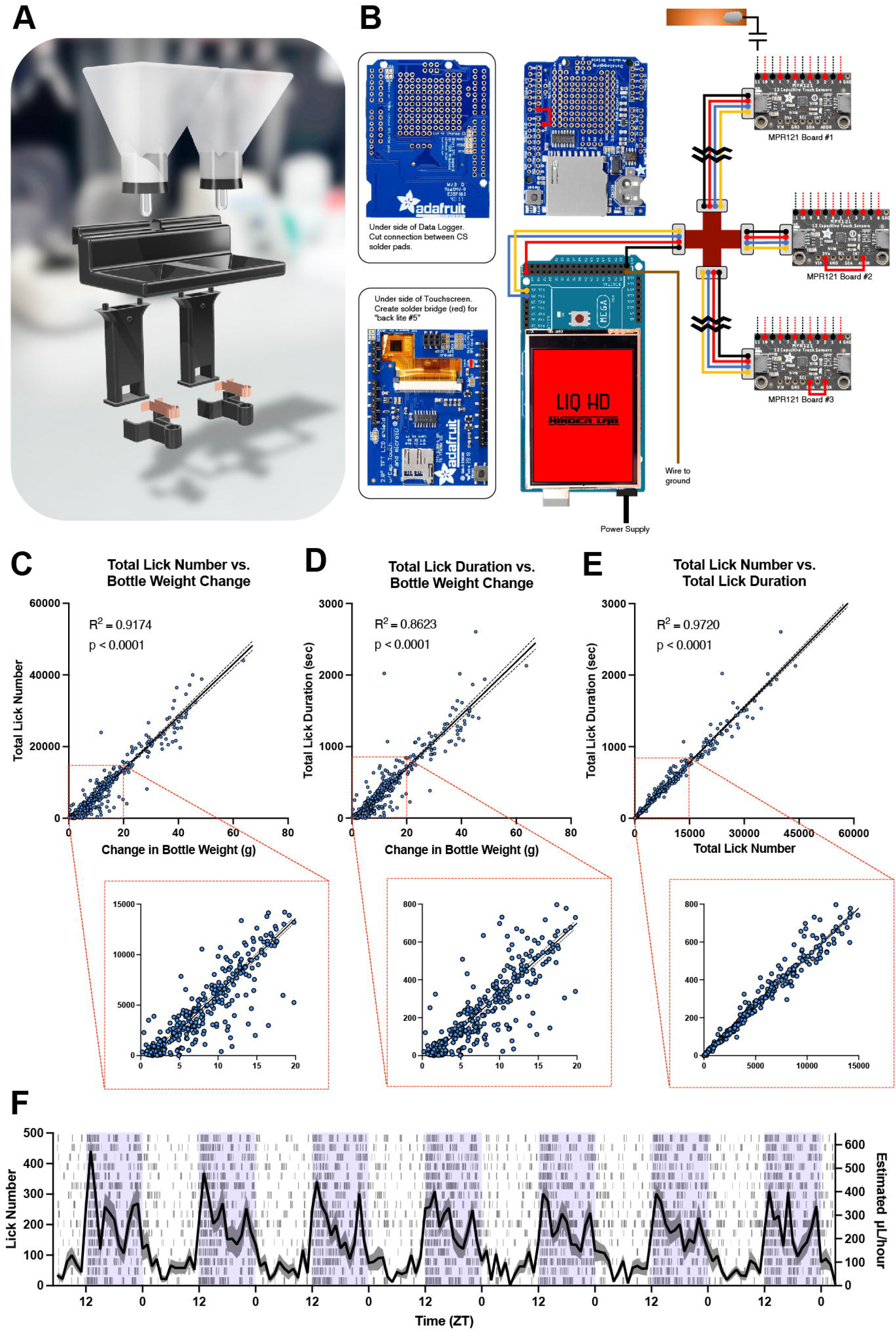
LIQ HD design and validation. ***A***, 3D rendering of LIQ HD disassembled components, including 3D-printed parts, rubber stoppers with sippers, and conductive copper foil tape. ***B***, LIQ HD electronic parts and wiring diagram. ***C***, Correlation between total lick number and change in bottle weight for each recording period. ***D***, Correlation between total lick duration and change in bottle weight for each recording period. ***E***, Correlation between total lick number and lick duration for each recording period. In correlation graphs, solid lines represent a fitted simple linear regression model, and dashed lines denote 95% confidence intervals. ***F***, Lick number and estimated water consumption throughout an undisturbed 7-day recording period with access to two water bottles. Data from both water bottles are combined to show the total intake per cage. The shaded purple area signifies the dark/active phase. The solid line represents the mean lick number and estimated intake in 1-hour bins, and the shaded area represents ±SEM (n = 16 cages). The raster plot displays licks detected in 1-minute bins for each cage.

In building the electronics, the Data Logger Shield and Touchscreen Shield must be slightly modified to be made compatible with the Arduino Mega. First, install the Shield Stacking Headers on the Data Logger Shield. Then, solder a wire connecting the CS pin to pin 7. On the underside of the Data Logger Shield, cut the thin connection on the CS solder pad by carefully etching with a sharp blade. On the Touchscreen Shield, create a solder bridge across the pads labeled “back lite #5” to allow for dimming of the screen during the animals’ dark cycle. Mount the Data Logger Shield and the Touchscreen Shield onto the Arduino Mega by aligning and inserting the headers. The MPR121 breakout boards communicate with the Arduino Mega via I^2^C, which allows the microcontroller to communicate with multiple devices connected to the same pins if they have different I^2^C addresses. The MPR121 Board #1 will remain unmodified, but to modify the I^2^C address of the other MPR121 boards, solder a wire connection from “ADDR” to “3V” on one board (Board #2) and from “ADDR” to “SDA” on another (Board #3). Next, solder each pair of wires of the 2-pin connectors to the sensor pins (0-11). For consistency, red wires are soldered to even numbers, and black to odd numbers. We provide additional reinforcement from accidental wire detachment by applying a layer of hot glue over the solder points. Inputs 0-11 on Board #1 correspond to sensors 1-12 (cages 1-6), inputs 0-11 on Board #2 correspond to sensors 13-24 (cages 7-12), and Inputs 0-11 on Board #3 correspond to sensors 25-36 (cages 13-18). To connect the boards, attach the Qwiic cable with breadboard jumpers to the Arduino Mega (blue – pin 20, yellow – pin 21, red – 5V, black – GND) and secure with hot glue. Connect the 4-way Qwiic Multiport Connector and plug in each MPR121 board with the Qwiic cables. Lastly, secure the device in the 3D-printed housing.

All 3D models were generated with Shapr3D and 3D-printed components were printed with PETG filament on an Ultimaker S5 printer. PETG was chosen for its high strength, durability, chemical resistance, ease of use, and food-safe properties. Bottles were printed with translucent filament and then coated internally with food-safe epoxy resin to prevent potential leaks and fill the space between printed layers. The in-cage device body was printed in pieces with black PETG. The legs were assembled to the upper portion with hot glue. For each device, two wire ends were soldered to the ends of two 3” × ¼” pieces of conductive copper tape. Copper tape is adhered to the inner part of each sipper clip, and wires are threaded up through the device body and out of the top. For consistency, red wire was used for the left side and black wire for the right side. Secure the sipper clips to the device legs with hot glue. Finally, solder the other 2-pin connectors to the device wires for easy connection to the MPR121 inputs.

The LIQ HD Arduino code was uploaded using the open-source Arduino IDE software (version 1.8.14 on MacOS). Users must first install the necessary libraries through the Arduino IDE before uploading the code. A detailed step-by-step guide, along with the Arduino code and 3D models, can be found at (https://github.com/nickpetersen93/LIQ_HD).

### Operation Instructions

The device starts up with a splash screen followed by the device home page. The home page displays the date, time, and various buttons. Press the cogwheel icon to access the settings page, where the user can modify the date, time, light/dark cycle times, sensor sensitivity and auto calibration settings, parameters to record, bin size, and SD sync interval. Default settings are preloaded, but users should determine which sensor threshold works best for them. On the home page, the user can designate which side the “experimental” solution is on in the cages (i.e. sucrose, quinine, ethanol, etc.) before pressing “Start” to initiate recording. The SD card can also be mounted and ejected to allow for users to transfer data.

To begin recording, first ensure all devices are secured in the animal cages with the sippers properly placed in the clips. Connect each 2-pin wire connector prior to initiating the recording with the “Start” button. After “Start” is pressed, the screen will display the data file name for 2 seconds and the sensors will calibrate. It is vital that the animals are not actively drinking and that the user steps away from the device during calibration for accurate measurements. Data is logged in 1-minute bins and saved to the SD card every 10 minutes by default. On the recording page, the screen will display the cumulative lick number for the sippers in each cage. While these values are updated internally every minute, the user must press “Refresh” to display the updated values. The user also has the option to pause the recording with the “Pause” button. Pausing the recording prevents any new licks from being recorded, safely ejects the SD card for data transfer, and writes a line in the data spreadsheet indicating that the recording was paused. Upon pressing “Resume”, the SD card is mounted, the data file name will be displayed for 2 seconds, and the sensors will recalibrate. If the SD card fails or is removed at any point during the recording, the screen will display a warning along with the date and time that the recording failed. Rarely the I^2^C communication on the Arduino can lock up due to glitches or electrical interference. We have included a timeout detection in the code to identify lock-ups and resume recording without losing significant time. In this case, sensors will be restarted and recalibrated automatically. A line in the data will be logged if a timeout occurs. When the user has finished recording, pressing “Save & Quit” will sync any unsaved data to the SD card and return the device to the home page.

### Infrared photobeam-break device build

The IR photobeam-based drink monitoring device was built from 3D-printed parts and commercially available sensors and components. The design of the device was based on designs from Frie and Khokhar (2019) and Godynyuk et al. (2019) with modifications to allow for 16 cages to be recorded from a single Arduino Mega (**Extended Figure 1–1A**). Briefly, 3D-printed parts were printed and assembled as described above. Beam-break sensor boards were assembled as previously described (Frie and Khokhar, 2019; Godynyuk et al., 2019) and secured into the 3D-printed device. A total of 32 photobeam sensors were wired to the Arduino input/output pins. A 1.3” OLED screen and two buttons were included to display total drink bout numbers for each cage and to operate the device. For this device, *bout number* is defined as the number of times the photobeam was interrupted and *bout duration* is the amount of time the beam was broken. Data were collected in one-minute increments.

### Determination of drinking bouts with LIQ HD

*Lick number* is defined as the number of times the animal licked the sipper, while *lick duration* is defined as the actual contact time on the sipper. As previously described (Siciliano et al., 2019), the start of a drinking *bout* is defined as three licks in less than 1 second and the end of a drinking bout is defined as no licks within 3 seconds. *Bout duration* is defined as the bout time minus the 3-second deadtime at the end of each bout. *Bout size* is defined as the number of licks that occurred during each bout. The *bout lick number* is defined as the number of licks that occur only during bouts, and the *bout lick duration* is the sipper contact time only during bouts. *Lick frequency*, defined as licks per second during bouts, for each bin is calculated by dividing the total bout length by the total bout duration in seconds:

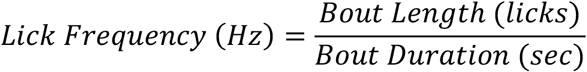

The *estimated inter-lick interval* is defined as the time between the offset of a lick and the onset of the subsequent lick. The average inter-lick interval for each bin is calculated by subtracting the total bout duration in milliseconds by the total bout lick duration and dividing by the total bout length:

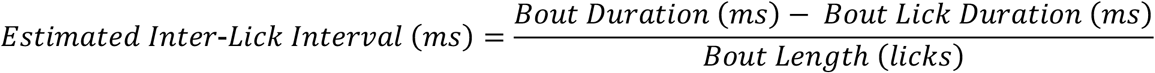

Bout microstructure did not significantly differ between the light and dark phase; thus, bout analysis is binned in 24-hour bins. Occasionally, the touch sensors do not release until touched again, which can erroneously inflate lick duration counts and be displayed in the data as bins with a lick duration value but without any recorded licks. For one-minute bins that have a lick duration value without a lick number value, or bins where the average lick duration (lick duration/lick number) was over 300ms, lick duration was changed to 0.

### Animals

Female C57BL/6J mice (8 weeks of age) were purchased from Jackson Laboratory (#000664). Mice were allowed to habituate to the animal facility for at least 7 days before the start of experimentation. All mice were singly housed on a standard 12hr light-dark cycle at 22-25°C with food and water available *ad libitum*. All fluid measurements conducted by experimenters took place during the light phase. All experiments were approved by the Vanderbilt University Institutional Animal Care and Use Committee (IACUC) and were carried out in accordance with the guidelines set in the Guide for the Care and Use of Laboratory Animals of the National Institutes of Health.

### Two-bottle voluntary choice tasks

For each experiment, mice were singly housed in cages containing LIQ devices. Each device held two bottles. Every experiment began with a 7-day habituation period, during which both bottles contained only water, to allow the mice to acclimate to the housing conditions and the presence of two bottles. During this time no measurements were taken. Next, weights were taken every 48-72 hours (on Monday, Wednesday, and Fridays) for experiment 1 or every 7 days (every Friday) for experiment 2 to gain baseline fluid intake levels from both bottles containing only water. Bottle placement was swapped each time bottles were weighed to account for potential side biases. Following 7 days of water-only measurements, the fluid in one of the bottles was changed to either sucrose, quinine, or ethanol (as described below) while the second bottle remained filled with water. Again, for experiment 1 weights taken by experimenters every 48-72 hours (on Monday, Wednesday, and Fridays), and for experiment 2 weights were taken every 7 days (every Friday). As before, bottle placement was swapped every time bottles were weighed. For each experimental solution, a doseresponse curve was performed in order from lowest to highest dose. For experiment 1, sucrose and quinine doses were each provided for 48-72 hours (72 hours for 0.5% sucrose and 48 hours each for 1% and 10% sucrose) before being switched to the next concentration. For experiment 2, 3% and 7% ethanol solutions were each provided for one week, followed by 4 weeks of 10% ethanol. The following doses were selected based on previous studies indicating altered intake and/or preference across a dose-response curve:

Sucrose: 0.5%, 1.0%, and 10% sucrose in tap water (Doyle et al., 2021; Glendinning et al., 2010; Zukerman et al., 2009)
Quinine: 0.01g/L, 0.03g/L, and 0.1g/L in tap water (Hodge et al., 1999; Winters et al., 2021)
Ethanol: 3%, 7%, and 10% ethanol in tap water (Centanni et al., 2019; Hodge et al., 1999; Winters et al., 2021)

### Statistical Analysis

Data was first extracted and processed with a custom MATLAB script. Unless otherwise stated, data were binned into 1-hour bins. For the calculation of bottle preference over time, the binned data were smoothed with a moving average with a sliding window length of 6. Statistical analyses were performed with Prism 9 (GraphPad). Pearson correlation coefficients and simple linear regressions were computed for all correlation pairs. Repeated measures ANOVA with corrections for multiple comparisons were performed as indicated in the figure legends.

### Code Accessibility

The code/software described in the paper is freely available online at (https://github.com/nickpetersen93/LIQ_HD). The code is available as Extended Data.

## Results

### Assessment of modified beam-break two-bottle choice system

The initial two-bottle choice pilot experiment conducted in the lab used a modified infrared (IR) beam break system (Frie and Khokhar, 2019; Godynyuk et al., 2019) (**Extended Figure 1–1A**) and singly housed female C57BL/6J mice. Specifically, the design of our device was adapted to make it compatible with mouse cages on our ventilated rack system (Lab Products Inc.) as well as to expand the volume the bottles could hold to 90 mL each. Following a week of habituating to two bottles containing water, four additional days of water-only availability were measured by the experimenter in daily sessions. This period was followed by a solution series, during which the fluid in one of the bottles was replaced with the following solutions: two consecutive days of 1% sucrose, one day of 10% sucrose, and one day of 0.1 g/L quinine. During this series, fluid intake was measured daily by an experimenter. To assess accuracy of the beam break-based measurements, the daily weight change values determined by experimenter intervention were correlated with the IR beam-break number and duration of bream-breaks recorded during the same recording period. When the data across the water and solution series were complied, we were able to closely replicate previous findings (Godynyuk et al., 2019) (**Extended Figure 1–1B,C**). The correlation between total beambreak bout number and change in bottle weight (R^2^ = 0.3810, F_(1, 85)_ = 52.31, p < 0.0001) as well as the correlation between total beam-break bout duration (seconds) and change in bottle weight (R^2^ = 0.3844, F_(1, 85)_ = 53.07, p < 0.0001), were statistically significant. However, we encountered several issues with the photobeam sensors throughout the recording period. First, multiple devices required repair or replacement due to mice chewing and damaging the photobeam sensors. We also found that drips hanging from the bottom of the sippers would trigger the sensor and overcount bout duration. Mice would frequently attempt to bury the sensors with bedding, especially in the initial recording days, which also erroneously triggered the sensors. In the presented data, data points were removed in bins where the bout duration lasted an entire bin or where there was a recorded bout duration without a corresponding bout number.

### Assessment of LIQ HD two-bottle choice system

While the data collected with the IR beam-break device significantly correlated with change in bottle weight and were nearly identical to published results, we sought to further improve on the accuracy and reliability of the behavioral recordings as well as increase the precision of the recorded data by designing a device, LIQ HD, that utilizes capacitive sensing technology to detect single licks. We took the modified design described above and replaced the IR beam-break sensors with a capacitance-sensing system. To validate the ability of LIQ HD to measure intake behaviors accurately, we performed a new series of two-bottle choice experiments with singly housed female C57BL/6J mice. Bottle weights were measured every 2-3 days (experiment 1) or every 7 days (experiment 2). Experiment 1 consisted of two groups of mice (8 mice each), where one group received a sucrose dose-response (0.5%, 1%, and 10% sucrose vs. water) and the other group received a quinine dose response (0.01g/L, 0.03g/L and 0.1g/L quinine vs. water). In experiment 2, 16 mice went through a water-only two-bottle choice session, followed by 8 of those mice advancing through an additional ethanol dose-response paradigm (3%, 7% and 10% EtOH vs. water). In both experiments, mice were first habituated to the LIQ devices with water in both bottles for 1 week, where no measurements were taken, and then given an additional week of water only. For experiment 1, the experimental solutions (sucrose and quinine) were changed every 2-3 days, which coincided with weight measurements and swapping the sides of the bottles. For experiment 2, the ethanol and water bottles were weighed, changed, and swapped sides every 7 days. The LIQ HD and bottle measurement data from experiments 1 and 2 were combined to correlate the total lick numbers and total lick durations from each recording period with the corresponding bottle weight measurements (**Figure 1C,D**). We observed a strong, significant correlation between both total lick number and change in bottle weight (R^2^ = 0.9174, F_(1, 363)_ = 4034, p < 0.0001) as well as total lick duration and change in bottle weight (R^2^ = 0.8623, F_(1, 363)_ = 2273, p < 0.0001), substantially higher R^2^ values than were reported in IR beam-break based systems. The fitted regression model for the correlation between lick number and change in bottle weight is Y = 736.4*X - 1172 (slope 95% CI, 713.6 to 759.2). Thus, on average, the mice took 736 licks to drink 1mL of fluid, or 1.4μL per lick. As expected, total lick number and total lick duration also strongly correlate (R^2^ = 0.9720, F_(1, 363)_ = 12601, p < 0.0001) with a fitted regression model of Y = 0.05132*X + 8.027 (slope 95% CI, 0.050 to 0.052) (**Figure 1E**). On average, the individual lick duration was 51ms. These metrics are consistent with the literature (Bollu et al., 2021; Mundl and Malmo, 1979; Parkison et al., 2012; Rossi and Yin, 2015), providing further evidence that LIQ HD is detecting individual lick events with high fidelity. In our tests, the LIQ HD system ran reliably undisturbed between periods of bottle weight measurements for at least seven days (**Figure 1F**).

#### **Experiment 1** - *LIQ HD validation in sucrose and quinine dose-response two-bottle choice tasks*

To test the LIQ HD system in common two-bottle choice paradigms, female C57BL/6J mice were split into two groups. One group underwent a two-bottle choice paradigm with a sucrose dose-response curve, and the other a quinine dose-response curve. Bottles were weighed and swapped sides every 2-3 days. When the recorded data were aggregated, we observed a strong, significant correlation between preference score calculated by total lick number and preference score calculated by bottle weight change (R^2^ = 0.8883, F_(1, 110)_ = 874.5, p < 0.0001) as well as between preference score calculated by total lick duration and preference score calculated bottle weight change (R^2^ = 0.8740, F_(1, 110)_ = 763, p < 0.0001) (**Figure 2A,B**). Additionally, with the added temporal resolution of LIQ HD over standard experimenter-determined measures, we also investigated changes in drink preference over time (**Figure 2C**). As expected, preference for the sucrose bottle over water increased with increasing concentrations of sucrose, and preference for the quinine bottle over water decreased with increasing concentrations of quinine (**Figure 2C**). Moreover, we examined changes in drinking behaviors over the light and dark cycle to detect potential deviations from the typical circadian drinking patterns, with examples displayed in **Figure 2D–I**. Overall, for the percent of total licks occurring per cycle in the sucrose group there was a significant main effect of light cycle (F_(1, 21)_ = 209.7, p < 0.0001) and interaction of light cycle x sucrose concentration (F_(2, 21)_ = 17.64, p < 0.0001). While the mice increased their total licks at the sucrose bottle as the sucrose concentration was increased from 0.5% to 1% sucrose, the percentage of the overall licks that occurred during the light and dark phases did not change (**Figure 2J**). In addition to further increasing total licks at the sucrose bottle during access to 10% sucrose (**Figure 2H**), the mice also displayed a unique increase in percent of sucrose consumption occurring during the light phase (0.5% vs 10% sucrose, p < 0.0001, 1% vs 10% sucrose, p < 0.0001), and corresponding decrease in the dark phase (0.5% vs 10% sucrose, p < 0.0001, 1% vs 10% sucrose, p < 0.0001) (**Figure 2J**). Mice exposed to increasing concentrations of quinine with water did not display any changes in the light cycle drinking pattern (**Figure 2K**) (significant main effect of light cycle only, F_(1, 21)_ = 485.5, p < 0.0001). Taken together, these data indicate that LIQ HD can accurately and consistently record overall preference scores, preference over time, and light cycle-dependent drinking patterns in classic two-bottle choice paradigms.

**Figure 2.**
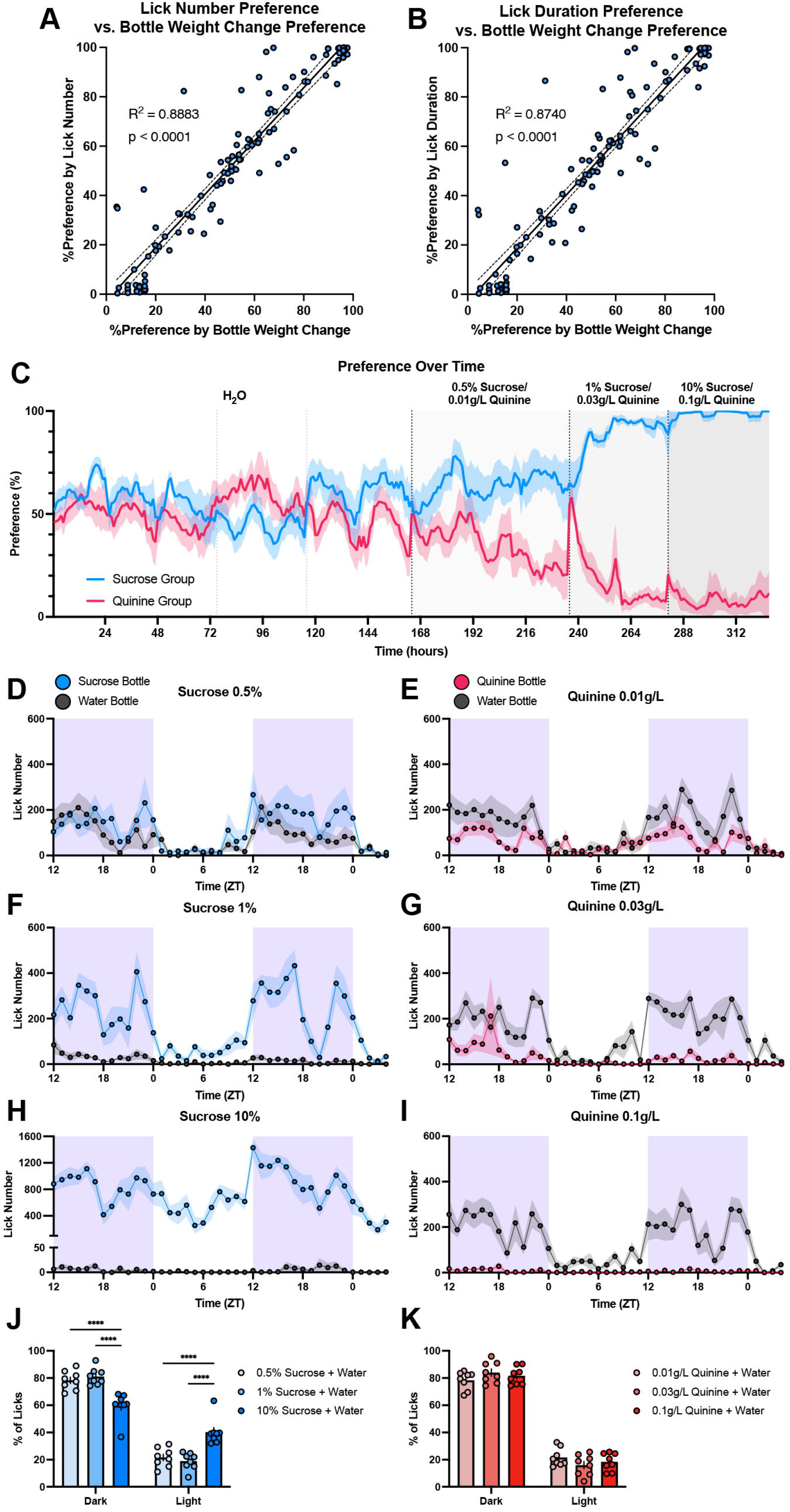
Utilizing LIQ HD to investigate changes in drink preference and light/dark cycle drinking patterns in sucrose and quinine two-bottle choice paradigms. ***A***, Correlation between percent preference calculated with lick number and percent preference calculated with change in bottle weight for each recording period. ***B***, Correlation between percent preference calculated with lick duration and percent preference calculated with change in bottle weight for each recording period. Solid lines represent fitted simple linear regression models, and dashed lines denote 95% confidence intervals. ***C***, Preference over time in 1-hour bins for the sucrose and quinine two-bottle choice dose-response paradigm. The solid line represents smoothed mean (sliding window length of 6), and the shaded area signifies ±SEM (n = 8 cages for each group). Vertical dashed lines indicate when bottles swapped sides and when indicated, a change in experimental solution in one bottle. ***D-I***, Lick number over time in 1-hour bins for recording periods for the two-bottle choice sucrose and quinine dose response paradigms. The shaded purple area signifies the dark/active phase. ***J***, Percentage of total licks that occur during the dark and light phases for recording periods of the sucrose dose-response paradigm (repeated measures two-way ANOVA, with the Geisser-Greenhouse correction and Tukey multiple comparisons test). During access to 10% sucrose, the percentage of licks that occurred during the dark phase was significantly decreased, and the percentage of licks that occurred during the light phase was significantly increased when compared to 0.5% and 1% sucrose availability. ***K***, Percentage of total licks during the dark and light phases for the quinine dose-response paradigm (repeated measures two-way ANOVA, with the Geisser-Greenhouse correction and Tukey multiple comparisons test). Shaded areas and error bars represent ±SEM. \*\*\*\**p* < 0.0001.

#### **Experiment 2** - *LIQ HD bout detection and microstructure analysis in a prolonged ethanol two-bottle choice task*

Because LIQ HD was able to accurately quantify behavioral data from sucrose and quinine two-bottle choice tasks, we next sought to validate its use in longer duration tasks. To do this, we turned to a 6-week continuous access ethanol two-bottle choice task. Further, given that LIQ HD can detect individual lick events, we sought to determine if the system is able to detect drinking bouts and record bout microstructure. To do this, we coded bout detection directly to the main LIQ HD Arduino code, where a “bout” begins when an animal licks at least 3 times within 1 second and ends when no licks have occurred over 3 seconds (Siciliano et al., 2019). With this we can also determine the lick number and lick duration during bouts, which allows us to calculate lick frequency and an estimated inter-lick interval.

First, to test the LIQ HD bout detection, 16 female C57BL/6J mice underwent a two-bottle choice task with access to two water bottles. Mice were first habituated for 1 week with a LIQ device with water in both bottles, during which no measurements were taken. Water-related measurements were then taken during a subsequent week of water-only access. We also sought to determine if LIQ HD is capable to run for prolonged undisturbed recording periods, thus experimenter measurements of bottle weights were taken only every 7 days. The LIQ data from each sipper (16 cages, 32 sippers) was binned into 1-hour bins and the average individual bout duration, bout size, bout lick frequency, and bout estimated inter-lick interval were calculated. Estimated inter-lick interval values >300ms were excluded from the analysis. The mean and median individual bout duration (seconds) during the water drinking period (n = 2919) were 5.35 ± 0.06 (SEM) and 4.77 (IQR 3.34 to 6.67) (**Figure 3A**). The mean and median of individual bout size (licks per bout) during the water drinking period (n = 2914) were 33.9 ± 0.34 (SEM) and 31.0 (IQR 22.0 to 42.5) (**Figure 3B**). The mean and median of lick frequency (Hz) (n = 2915) were 6.62 ± 0.03 (SEM) and 6.77 (IQR 5.81 to 7.63) (**Figure 3C**). The mean and median of estimated inter-lick interval (milliseconds) during the water drinking period (n = 2858) were 106 ± 0.79 (SEM) and 94.7 (IQR 79.1 to 118) (**Figure 3D**). These findings are consistent with results from previous studies (Bollu et al., 2021; Boughter et al., 2007; Johnson et al., 2010; Parkison et al., 2012; Raymond et al., 2018).

**Figure 3.**
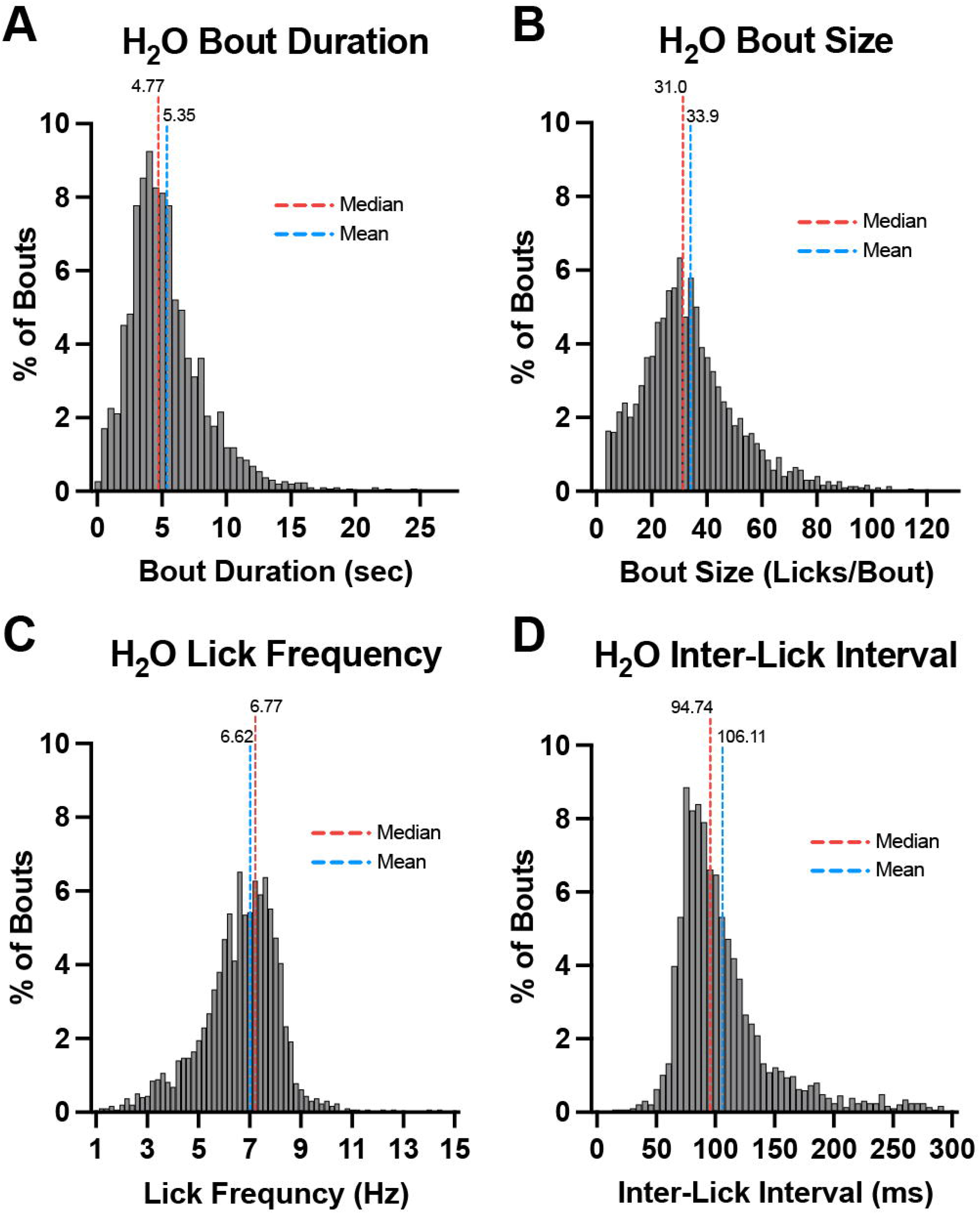
Histograms of average baseline bout microstructure measurements from 1-hour bins during 1-week access to two water bottles (N = 16 mice, n = 32 bottles). Histogram displaying bout duration (***A***), bout size (***B***), lick frequency (***C***), and estimated inter-lick interval (***D***). Red and blue dashed lines represent the median and mean, respectively.

Following the LIQ bout detection validation, 8 of the female C57BL/6J mice underwent a two-bottle choice ethanol drinking paradigm. The two-bottle choice paradigm consisted of an ethanol ramp where one of the water bottles was replaced with an ethanol solution. During the ramp, mice received one week of 3% ethanol, one week of 7% ethanol, and four weeks of 10% ethanol. As stated above, the system can accurately and consistently record drinking behavior over a 7-day period (**Figure 1F**). Moreover, LIQ HD withstood 8 weeks (including water only access and habituation) of continuous usage without any devices needing repair or replacement during experimentation.

To determine the correlation of bout number and bout duration with change in bottle weight, as well as the correlation of preference score calculated by bout number and bout duration with the preference score calculated by change in bottle weight, the water two-bottle choice and ethanol two-bottle choice data were aggregated. We observed a strong, significant correlation between total bout number and change in bottle weight (R^2^ = 0.8062, F_(1, 156)_ = 648.8, p < 0.0001) as well as between total bout duration and change in bottle weight (R^2^ = 0.8787, F_(1, 156)_ = 1130, p < 0.0001) (**Figure 4A,B**). The fitted regression models for bout number versus change in bottle weight and bout duration versus change in bottle weight are Y = 21.51*X + 0.8541 (slope 95% CI, 19.84 to 23.17) and Y = 157.533*X - 37270 (slope 95% CI, 148.276 to 166.790), respectively. These data indicate that on average mice drink 1mL over 21.5 bouts, or 46.5μL per bout, and on average take 157.5 seconds to drink 1mL during bouts, or 6.35μL per second. We also found a strong, significant correlation between preference score calculated by total bout number and preference score calculated by bottle weight change (R^2^ = 0.8343, F_(1, 69)_ = 347.4, p < 0.0001) as well as between preference score calculated by total bout duration and preference score calculated by bottle weight change (R^2^ = 0.8791, F_(1, 69)_ = 501.8, p < 0.0001) (**Figure 4C,D**). While bout number and bout duration may not be as reliable of a predictor of total change in bottle weight compared to total lick number (R^2^ = 0.9174), these data suggest that the LIQ HD bout detection software can accurately detect individual clustered drinking events throughout long recording periods.

**Figure 4.**
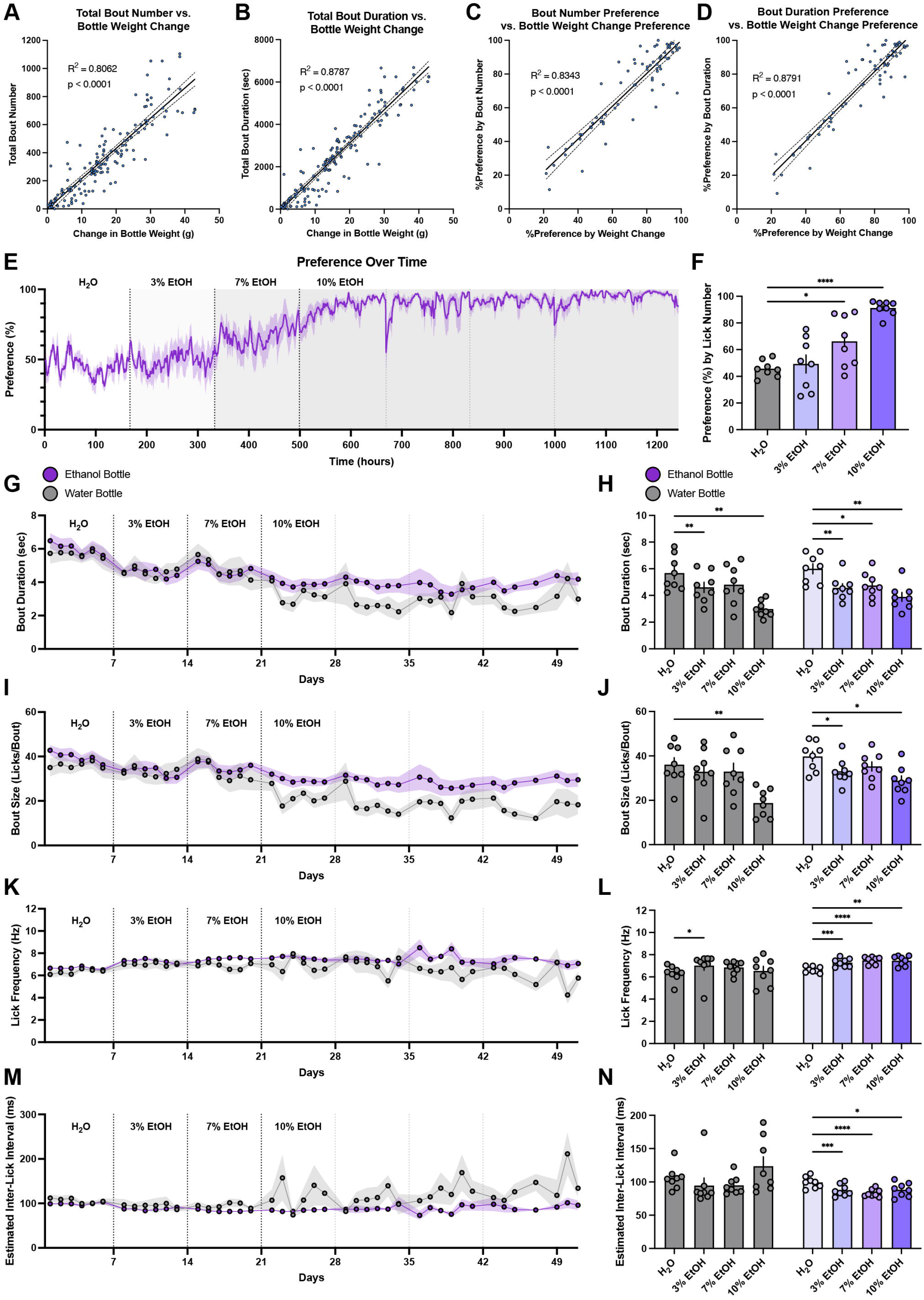
Validation of LIQ HD bout detection and bout microstructure in an ethanol two-bottle choice paradigm. ***A***, Correlation between total bout number and change in bottle weight for each recording period. ***B***, Correlation between total bout duration and change in bottle weight for each recording period. ***C***, Correlation between percent preference calculated with bout number and percent preference calculated with change in bottle weight for each recording period. ***D***, Correlation between percent preference calculated with bout duration and percent preference calculated with change in bottle weight for each recording period. Solid lines represent a fitted simple linear regression model, and dashed lines denote 95% confidence intervals. ***E***, Preference over time in 1-hour bins for the ethanol two-bottle choice paradigm. The solid line represents smoothed mean (sliding window length of 6), and shaded area signifies ±SEM (n = 8 cages). Vertical dashed lines indicate when bottles swapped sides and when indicated, a change of experimental solution in one bottle. ***F***, Average preference for experimental bottle during ethanol two-bottle choice paradigm (repeated measures one-way ANOVA, with the Geisser-Greenhouse correction and Dunnett’s multiple comparisons test). Mice show significantly increased preference compared to baseline for 7% and 10% ethanol. Bout duration (***G***), bout size (***I***), lick frequency (***K***), and estimated inter-lick interval (***M***) over time in 24-hour bins for water and ethanol bottles. Average bout duration (***H***), bout size (***J***), lick frequency (***L***), and estimated inter-lick interval (***N***) during access to two water bottles and access to water with increasing concentrations of ethanol (repeated measures two-way ANOVA, with the Geisser-Greenhouse correction and Dunnett’s multiple comparisons test). At the ethanol bottle, mice display a significant decrease in bout duration (***G***), bout size (***I***), and inter-lick interval (***N***) with a significant increase in lick frequency (***L***) during access to increasing concentrations of ethanol. At the water bottle, mice display a significant increase in lick frequency only during access to 3% ethanol (***L***), a significant decrease in bout duration during access to 3% and 10% ethanol (***H***), and a significant decrease in bout size only during access to 10% ethanol (***J***). Shaded areas and error bars represent ±SEM. \**p* < 0.05, \*\**p* < 0.01, \*\*\**p* < 0.001, \*\*\*\**p* < 0.0001.

In addition to preference over time in the prolonged ethanol two-bottle choice paradigm, where mice showed expected significantly elevated preference for both 7% ethanol (p = 0.0481) and 10% ethanol (p < 0.0001) compared to water (**Figure 4E–F**), LIQ HD can also be used to analyze drinking bout microstructure over time (**Figure 4G–N**). Bout microstructure data were grouped into 24-hour bins, as we did not observe differences between the light and dark cycle (data not shown). Throughout this increasing preference for ethanol, the average bout duration, bout size, lick frequency and estimated inter-lick interval were altered. Although bouts at the ethanol bottle became more frequent compared to those at the water bottle, access to ethanol significantly decreased the average bout duration (significant main effect of ethanol concentration, F_(2.129, 29.80)_ = 23.68, p < 0.0001) at both the water bottle (H_2_O vs. 3% EtOH, p = 0.0096, H2O vs. 10% EtOH, p = 0.0022) and the ethanol bottle (H2O vs. 3% EtOH p = 0.0055, H2O vs. 7% EtOH, p = 0.0310, H2O vs. 10% EtOH, p = 0.0054) (**Figure 4H**). Bout size also significantly decreased (significant main effect of ethanol concentration, F_(1.882, 26.34)_ = 17.01, p < 0.0001) at the ethanol bottle during ethanol exposure (H2O vs. 3% EtOH, p = 0.0288, H2O vs. 10% EtOH, p = 0.0189), but only decreased at the water bottle during exposure to the highest ethanol dose (H2O vs. 10% EtOH, p = 0.0047) (**Figure 4J**). Interestingly, lick frequency significantly increased (significant main effect of ethanol concentration, F_(2.300, 32.20)_ = 8.236, p = 0.0008) at the ethanol bottle at all doses (H2O vs. 3% EtOH, p = 0.0005, H2O vs. 7% EtOH, p < 0.0001, H2O vs. 10% EtOH, p = 0.0017) (**Figure 4L**) with a concurrent decrease in the estimated inter-lick interval (significant main effect of ethanol concentration, F_(2.300, 32.20)_ = 8.236, p = 0.0008, significant interaction ethanol concentration x bottle F_(3, 42)_ = 3.801, p = 0.0169, H2O vs. 3% EtOH, p = 0.0002, H2O vs. 7% EtOH, p < 0.0001, H2O vs. 10% EtOH, p = 0.0236) (**Figure 4N**). We only observed an increase in lick frequency at the water bottle during access to 3% ethanol (H2O vs. 3% EtOH, p = 0.0434) without any significant differences in estimated inter-lick interval (**Figure 4L,N**). Overall, these data replicated known effects of prolonged, continuous access to increasing concentrations of ethanol in an undisturbed home cage environment on preference for ethanol over water and revealed significant changes to the bout lick microstructure in female C57BL/6J mice.

## Discussion

Here we present LIQ HD, Lick Instance Quantifier Home cage Device, a capacitance sensor-based two-bottle choice open-source system capable of detecting undisturbed licking behavior in a true rodent home cage environment. A single LIQ HD Arduino system records drinking behavior in up to 18 cages. The system includes a touchscreen with a graphical user interface for an intuitive user experience. LIQ HD detects single lick events, and lick number and duration strongly correlate with change in liquid volume as measured by manually weighing the bottles. The use of capacitive sensors significantly outperformed our modified beam-break sensor based device (R^2^ of 0.9174 versus 0.3844), which closely matched the accuracy of other beam-break devices (Godynyuk et al., 2019). Additionally, each 3D-printed bottle holds roughly 90mL of liquid, allowing for prolonged, undisturbed recording sessions. In this study we utilized LIQ HD continuously for several months with undisturbed recordings lasting up to 7 days. We observed that female C57BL/6J mice drank about 7mL of water per day, suggesting that the system could potentially run for multiple weeks undisturbed. It is important to note that the maximum length of the recording period will depend on preference values, mouse strain (Bachmanov et al., 2002), and the animal housing regulations of the research institution.

In a series of two-bottle choice paradigms, we show that LIQ HD accurately measures drink preference, and the minute-by-minute data also allow for the analysis of circadian drinking patterns. For example, here we show that access to 10% sucrose shifts the typical dark/light drinking pattern, with a significantly greater percentage of licks occurring in the light phase and less in the dark phase when compared to 0.5% or 1% sucrose availability (**Figure 2J**). These features may be helpful in studying individual differences in drinking behaviors, such as investigating differences in the acquisition of drink preference or susceptibility to circadian dysregulation. Moreover, the ability of the LIQ HD to detect individual licks allows for analysis of the mouse bout microstructure, further simplified by our software which detects bouts directly from the Arduino source code. While it is challenging to compare lick microstructure across various experimental modalities, such as unlimited continuous access, intermittent access, operant conditioning tasks, etc., our baseline bout microstructure measurements are consistent with the data in the literature (Bollu et al., 2021; Boughter et al., 2007; Johnson et al., 2010; Parkison et al., 2012; Raymond et al., 2018). Utilizing this bout detection and microstructure system, we observed significant changes to drinking structure over time in a classic ethanol two-bottle choice task. Taken together, LIQ HD can be utilized to investigate drinking behavior and bout microstructure with high temporal resolution and accuracy for two-bottle choice tasks with prolonged undisturbed recording periods.

The capacitance-sensing LIQ HD system has several advantages compared to currently available IR photobeam-break devices. In addition to significantly improved accuracy and ventilated rack compatibility, the LIQ HD devices are more resilient. In our experience, photobeam sensors were frequently subject to destruction by rodent chewing. In the LIQ HD system, if the sipper remains in contact with the conductive copper foil tape, LIQ HD will continue to detect licks, creating a low likelihood that a mouse could destroy the device to the point that it malfunctions. During these experiments and initial pilot studies, the 16 LIQ HD devices ran for >100 days without any device failures. It is possible that if used with rats or animals with increased chewing behavior, such stressor opioid-exposed mice, the 3D-printed clips that secure the copper tape and sipper will require more frequent repair. However, we expect some damage and have accordingly designed the device so that the clip is easily removable and replaceable. Moreover, the capacitive sensor boards communicate with the Arduino controller though I^2^C communication rather than through the direct input/output pins on the microcontroller. This allows for many devices to run off a single Arduino, as the system is no longer limited to the number of available pins but rather the ability of the sensor boards to have unique I^2^C addresses. In its current state LIQ HD utilizes 3 12-channel MPR121 capacitive sensor boards, for a total of 36 sippers, but it can be readily expanded by the end user to use 4 boards for 48 sippers. With the addition of an I^2^C address multiplexer, users can connect multiple sensor boards that have the same address, further scaling LIQ HD.

Like any open-source system, LIQ HD has several limitations when considering its use. Unlike battery-powered systems, our system must be continuously plugged into an outlet and will stop recording if power is lost. The capacitive sensors may also be subject to electrical interference if not properly grounded. To avoid this interference, the listed power supply contains a grounding prong, and we recommend an additional grounding wire from the Arduino to another grounding source. In addition, the capacitive sensors likely render this system incompatible with electrophysiology, though we have not tested this. It is also important to note that because electrical wires increase capacitance and thus affect sensor sensitivity, it is essential that all wire lengths connecting cages are kept consistent and as short as possible. Finally, LIQ HD is limited by the same factors that limit most two-bottle choice systems, such as requiring animals to be singly housed and requiring periodic switching of bottle sides to avoid side bias.

To conclude, LIQ HD is an affordable, easy-to-build, multi-home cage lickometer system that utilizes capacitive sensors for accurate lick detection. The devices hold sufficient liquid bottle weight change in two-bottles for prolonged undisturbed recordings of drinking behavior and bout microstructure and are highly resistant to functional damage from rodent manipulation. The current system is designed to record up to 18 cages simultaneously, and its precision can eliminate the need for cumbersome bottle weighing, thus rendering it suitable for high-throughput experiments. We encourage users to utilize the open-source code and designs to expand upon our current system to make LIQ HD compatible with other home cages, increase the scale of recordings, and improve overall efficiency.

**Table 1.** A complete list of LIQ HD building components with manufacturer and distributor product information, quantity per system, price per component unit, and total cost in $USD. Prices are accurate as of November 2022. LIQ HD list of components

| Item | Manufacturer (Product #) | Distributor (Product #) | Qty. | Price/Unit (\$USD) | Total (\$USD) |
| --- | --- | --- | --- | --- | --- |
| Arduino Mega 2560 Rev3 | Arduino (A000067) | Digi-Key (1050-1018-ND) | 1 | 48.40 | 48.40 |
| 2.8" Capacitive Touch Shield | Adafruit (1651) | Digi-Key (1528-1027-ND) | 1 | 34.95 | 34.95 |
| Adafruit Data Logging Shield | Adafruit (1141) | Digi-Key (1528-1044-ND) | 1 | 13.95 | 13.95 |
| Shield Stacking Headers for Arduino | Adafruit (85) | Digi-Key (1528-1074-ND) | 1 | 1.95 | 1.95 |
| 12.5mm 3V CR1220 Lithium Battery | Jauch Quartz (CR 1220 JAUCH SB) | Digi-Key (1908-CR1220JAUCHSB-ND) | 1 | 1.09 | 1.09 |
| 8GB microSD Card with Adapter | Adafruit (1294) | Adafruit (1294) | 1 | 9.95 | 9.95 |
| MPR121 12-Key Capacitive Touch Sensor Breakout | Adafruit (1982) | Digi-Key (1528-1038-ND) | 3 | 7.95 | 23.85 |
| 12V 5A Switching Power Supply | Adafruit (352) | Digi-Key (1528-1664-ND) | 1 | 24.95 | 24.95 |
| Qwiic Multiport Connector | SparkFun (BOB-18012) | Digi-Key (1568-BOB-18012-ND) | 1 | 2.10 | 2.10 |
| Qwiic Cable Breadboard Jumper | SparkFun (PRT-14425) | Digi-Key (1568-1709-ND) | 1 | 1.50 | 1.50 |
| Flexible Qwiic Cable – 500mm | SparkFun (PRT-17257) | Digi-Key (1568-PRT-17257-ND) | 2 | 2.38 | 4.76 |
| Flexible Qwiic Cable – 50mm | SparkFun (PRT-17260) | Digi-Key (1568-PRT-17260-ND) | 1 | 1.06 | 1.06 |
| 2.5mm Pitch 2-pin Cable Matching Pair | Adafruit (4872) | Digi-Key (1528-4872-ND) | 18 | 0.95 | 17.10 |
| Hook-up Wire Spool Set - 22AWG Stranded-Core - 10x25ft | Adafruit (3175) | Digi-Key (1528-1746-ND) | 1 | 29.95 | 29.95 |
| Solid-Core Wire Spool - 25ft - 22AWG - Red | Adafruit (288) | Digi-Key (1528-1750-ND) | 1 | 2.95 | 2.95 |
| Kable Kontrol™ Heat Shrink Tubing - 2:1 Polyolefin – 3/16" - 10' Various Colors | Cables Ties And More (HS357-S10) | Cables Ties And More (HS357-S10) | 6 | 5.97 | 35.82 |
| Kable Kontrol™ Heat Shrink Tubing - 2:1 Polyolefin – 1/8" - 10' Black | Cables Ties And More (HS355-S10-BLACK) | Cables Ties And More (HS355-S10-BLACK) | 1 | 5.50 | 5.50 |
| Translucent Clear PRO Series PETG Filament - 2.85mm (1kg) (or equivalent) | MatterHackers (M-A9A-GJF6) | MatterHackers (M-A9A-GJF6) | 1 | 55.00 | 55.00 |
| Black PRO Series PETG Filament - 2.85mm (1kg) (or equivalent) | MatterHackers (M-M82-QGPE) | MatterHackers (M-M82-QGPE) | 2 | 55.00 | 110 |
| Purple MH Build Series PLA Filament - 2.85mm (1kg) (or equivalent) | MatterHackers(M-ZR7-0EQU) | MatterHackers(M-ZR7-0EQU) | 1 | 20.87 | 20.87 |
| Food Safe Clear Epoxy Resin 1 Gallon | ZDSpoxy (B07VZBC6CZ) | Amazon (727040490164) | 1 | 89.98 | 89.98 |
| One-Hole Rubber Stoppers Size 5.5 – Pack of 28 | Grainger (16ZD52) | Grainger (16ZD52) | 2 | 22.71 | 45.42 |
| 1.5" Straight Sipper Tube (custom order request) | Ancare (OT-99) | Ancare (OT-99) | 36 | 1.77 | 63.72 |
| ¼" Conductive Copper Tape | Bertech (CFT-1/4) | Digi-Key (4393-CFT-1/4-ND) | 1 | 11.84 | 11.84 |

**Total 656.66****Average Price per Cage 36.48**

## Supporting information

Extended Date - LIQ HD Arduino Source Code

## Acknowledgements

We thank Lisa Kim for assistance in assembling the LIQ HD devices.

## Figure & Table Legends

**Extended Figure 1-1.**
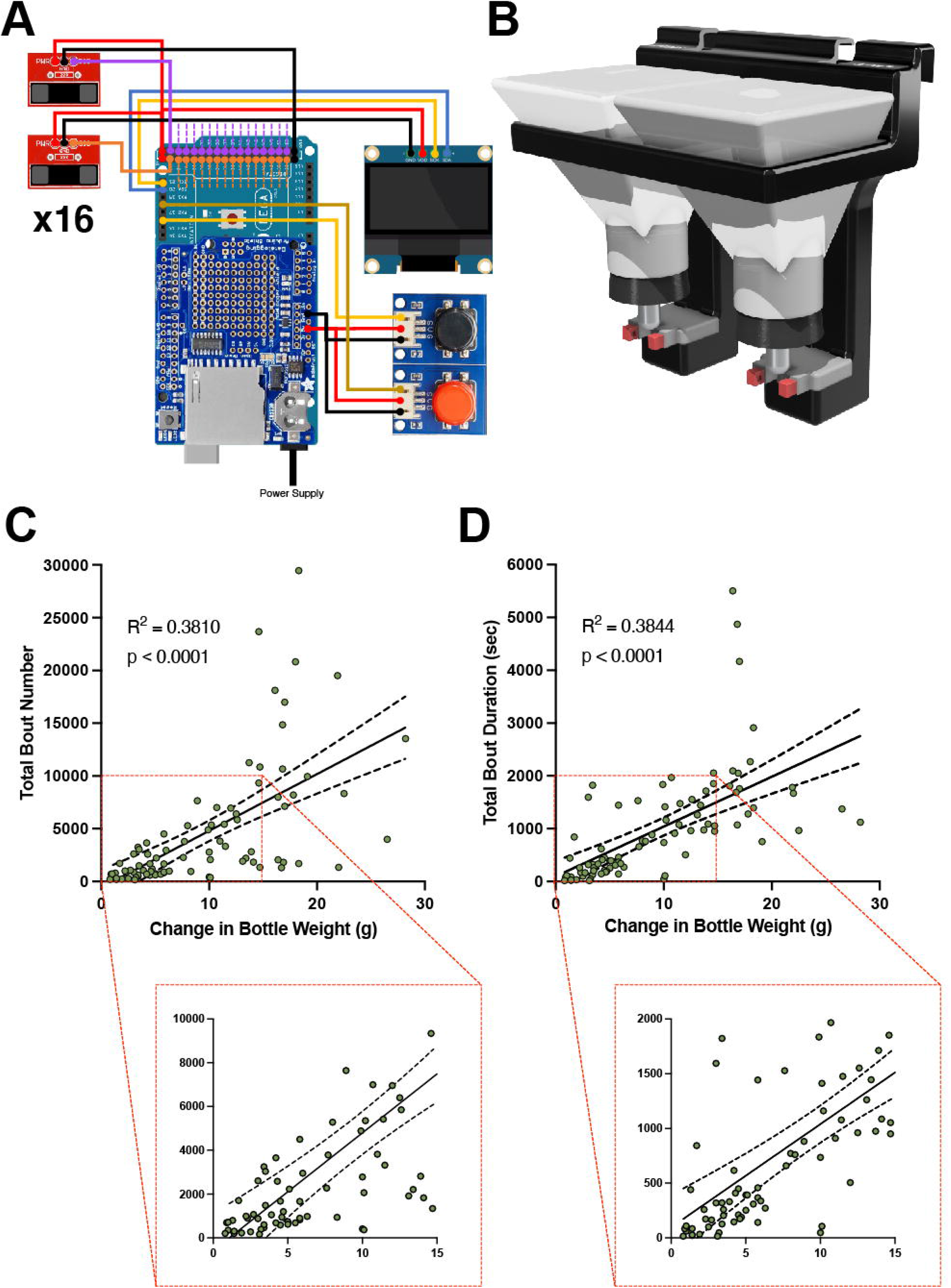
Infrared beam-break-based two-bottle choice device design and validation. ***A***, Electronic parts and wiring diagram for the beam-break system. ***B***, 3D rendering of beam-break device, including 3D-printed components, rubber stoppers and sippers, and photobeam sensors (red). ***C***, Correlation between total bout number and change in bottle weight for each recording period. ***D***, Correlation between total bout duration and change in bottle weight for each recording period. Solid lines represent fitted simple linear regression models, and dashed lines denote 95% confidence intervals.

