## Extended Date - LIQ HD Arduino Source Code for "LIQ HD (Lick Instance Quantifier Home cage Device): An open-source tool for recording undisturbed two-bottle drinking behavior in a home cage environment": LIQ_functions.ino

```
//functions used throughout code

//======================================================================================

void reset_total_LN()
{
  for (int k = 0; k < 36; k++) {
    total_LN[k] = 0; //reset totals
  }
}

//======================================================================================

void reset_variables()
{
  //Reset lick variables
  for (int k = 0; k < 36; k++) {
    LickNumber[k] = 0;
    LickDuration[k] = 0;
    BoutNumber[k] = 0;
    BoutDuration[k] = 0;
    BoutLickNumber[k] = 0;
    BoutLickDuration[k] = 0;
  }
}

//==========================================================================================

void Record_Licks()
{
  for (int k = 0; k < 12; k++) { //for cap sensor 1
    currtouched1 = cap.touched();
    if ((currtouched1 & _BV(k)) && !licking[k]) {
      DEBUG_PRINT(k); DEBUG_PRINTLN(" touched");
      cal_timer = millis();
      time_now[k] = millis();
      licking[k] = true;
      LickNumber[k] += 1;
      lick_bout_countdown[k] += 1;
      if (lick_bout_countdown[k] == 1) {
        bout_start_timer[k] = millis();
      }
      if (lick_bout_countdown[k] == 3 && millis() - bout_start_timer[k] <= 1000) {
        BoutNumber[k] += 1;
        bout_timer[k] = millis();
        in_bout[k] = true;
        DEBUG_PRINT(k); DEBUG_PRINTLN(" bout started!");
      }
      if (lick_bout_countdown[k] >= 3 && millis() - bout_timer[k] <= 3000) { //change this to just if(in_bout && millis() - bout_timer[k] <= 3000)?
        BoutLickNumber[k] += 1;
        bout_timer[k] = millis();
      }
    }
    if (!(currtouched1 & _BV(k)) && licking[k]) {
      DEBUG_PRINT(k); DEBUG_PRINTLN(" released");
      Elapsedtime[k] = millis() - time_now[k];
      LickDuration[k] = LickDuration[k] + Elapsedtime[k];
      licking[k] = false;
      if (in_bout[k]) {
        BoutLickDuration[k] = BoutLickDuration[k] + Elapsedtime[k];
      }
      DEBUG_PRINTLN(LickNumber[k]);
      DEBUG_PRINTLN(LickDuration[k]);
    }
  }
  lasttouched1 = currtouched1;

  for (int k = 12; k < 24; k++) { //for cap sensor 2
    currtouched2 = cap2.touched();
    int s = k - 12;
    if ((currtouched2 & _BV(s)) && !licking[k]) {
      DEBUG_PRINT(k); DEBUG_PRINTLN(" touched");
      cal_timer = millis();
      time_now[k] = millis();
      licking[k] = true;
      LickNumber[k] += 1;
      lick_bout_countdown[k] += 1;
      if (lick_bout_countdown[k] == 1) {
        bout_start_timer[k] = millis();
      }
      if (lick_bout_countdown[k] == 3 && millis() - bout_start_timer[k] <= 1000) {
        BoutNumber[k] += 1;
        bout_timer[k] = millis();
        in_bout[k] = true;
        DEBUG_PRINT(k); DEBUG_PRINTLN(" bout started!");
      }
      if (lick_bout_countdown[k] >= 3 && millis() - bout_timer[k] <= 3000) { //change this to just if(in_bout && millis() - bout_timer[k] <= 3000)?
        BoutLickNumber[k] += 1;
        bout_timer[k] = millis();
      }
    }
    if (!(currtouched2 & _BV(s)) && licking[k]) {
      DEBUG_PRINT(k); DEBUG_PRINTLN(" released");
      Elapsedtime[k] = millis() - time_now[k];
      LickDuration[k] = LickDuration[k] + Elapsedtime[k];
      licking[k] = false;
      if (in_bout[k]) {
        BoutLickDuration[k] = BoutLickDuration[k] + Elapsedtime[k];
      }
      DEBUG_PRINTLN(LickNumber[k]);
      DEBUG_PRINTLN(LickDuration[k]);
    }
  }
  lasttouched2 = currtouched2;

  for (int k = 24; k < 36; k++) { //for cap sensor 3
    currtouched3 = cap3.touched();
    int s = k - 24;
    if ((currtouched3 & _BV(s)) && !licking[k]) {
      DEBUG_PRINT(k); DEBUG_PRINTLN(" touched");
      cal_timer = millis();
      time_now[k] = millis();
      licking[k] = true;
      LickNumber[k] += 1;
      lick_bout_countdown[k] += 1;
      if (lick_bout_countdown[k] == 1) {
        bout_start_timer[k] = millis();
      }
      if (lick_bout_countdown[k] == 3 && millis() - bout_start_timer[k] <= 1000) {
        BoutNumber[k] += 1;
        bout_timer[k] = millis();
        in_bout[k] = true;
        DEBUG_PRINT(k); DEBUG_PRINTLN(" bout started!");
      }
      if (lick_bout_countdown[k] >= 3 && millis() - bout_timer[k] <= 3000) { //change this to just if(in_bout && millis() - bout_timer[k] <= 3000)?
        BoutLickNumber[k] += 1;
        bout_timer[k] = millis();
      }
    }
    if (!(currtouched3 & _BV(s)) && licking[k]) {
      DEBUG_PRINT(k); DEBUG_PRINTLN(" released");
      Elapsedtime[k] = millis() - time_now[k];
      LickDuration[k] = LickDuration[k] + Elapsedtime[k];
      licking[k] = false;
      if (in_bout[k]) {
        BoutLickDuration[k] = BoutLickDuration[k] + Elapsedtime[k];
      }
      DEBUG_PRINTLN(LickNumber[k]);
      DEBUG_PRINTLN(LickDuration[k]);
    }
  }
  lasttouched3 = currtouched3;

  for (int k = 0; k < 36; k++) {
    if (lick_bout_countdown[k] <= 2 && millis() - bout_start_timer[k] >= 1000) {
      bout_start_timer[k] = 0;
      lick_bout_countdown[k] = 0;
    }
    if (lick_bout_countdown[k] >= 3 && millis() - bout_timer[k] >= 3000) {
      DEBUG_PRINT(k); DEBUG_PRINTLN(" bout ended!");
      in_bout[k] = false;
      BoutLickNumber[k] += 2; //add licks missed before bout started
      BoutDuration[k] = BoutDuration[k] + (millis() - bout_start_timer[k]);
      bout_start_timer[k] = 0;
      lick_bout_countdown[k] = 0;
    }
  }
}


//======================================================================================

void update_sippers()
{
  for (int k = 0; k < 36; k++) {
    if (licking[k]) {
      Elapsedtime[k] = millis() - time_now[k];
      LickDuration[k] = LickDuration[k] + Elapsedtime[k];
      if (in_bout[k]) {
        BoutDuration[k] = BoutDuration[k] + (millis() - bout_start_timer[k]);
        BoutLickDuration[k] = BoutLickDuration[k] + Elapsedtime[k];
      }
    }
  }
}

//======================================================================================

void reset_time_now()
{
  for (int k = 0; k < 36; k++) {
    time_now[k] = millis(); //reset time now
    bout_start_timer[k] = millis();
  }
}

//======================================================================================

void calc_total_LN()
{
  for (int k = 0; k < 36; k++) {
    total_LN[k] = total_LN[k] + LickNumber[k]; //calculate total LN for each sipper
  }
}

//======================================================================================

void wait(unsigned long ms)
{
  unsigned long X = millis();
  while (millis() < X + ms) {} //wait without delay for ms (milliseconds)
}

//======================================================================================

void set_defaults()
{
  default_lights_on = lights_on;
  default_lights_off = lights_off;
  default_touch_threshold = touch_threshold;
  default_release_threshold = release_threshold;
  default_LOG_INTERVAL = LOG_INTERVAL;
  dafault_SYNC_INTERVAL = SYNC_INTERVAL;
  default_auto_cal = auto_cal;
  default_auto_cal_time = auto_cal_time;
  default_auto_cal_sec_since_last_lick = auto_cal_sec_since_last_lick;
}

//======================================================================================

void auto_calibration()
{
  if (auto_cal) {
    now = rtc.now();
    if (now.hour() == auto_cal_time && millis() - cal_timer > 1000 * auto_cal_sec_since_last_lick && auto_cal_flag) {
      auto_cal_flag = false;
      cap.begin(0x5C); cap2.begin(0x5B); cap3.begin(0x5C);
      cap.setThresholds(touch_threshold, release_threshold);
      cap2.setThresholds(touch_threshold, release_threshold);
      cap3.setThresholds(touch_threshold, release_threshold);
    }
    if (now.hour() != auto_cal_time) {
      auto_cal_flag = true;
    }
  }
}
```
