## Extended Date - LIQ HD Arduino Source Code for "LIQ HD (Lick Instance Quantifier Home cage Device): An open-source tool for recording undisturbed two-bottle drinking behavior in a home cage environment": LIQ_HD_Ardunio_Mega.ino

```
#include <Adafruit_GFX.h>
#include <SPI.h>
#include <Wire.h>
#include <SD.h>
#include "RTClib.h"
#include "Adafruit_MPR121.h"
#include <Adafruit_ILI9341.h>
#include <Adafruit_FT6206.h>
#include <math.h>
#include <Fonts/FreeSans9pt7b.h>
#include <Fonts/FreeSans12pt7b.h>
#include <Fonts/FreeSans18pt7b.h>
#include <Fonts/FreeSansBold9pt7b.h>
#include <Fonts/FreeSansBold12pt7b.h>
#include <Fonts/FreeSansBold18pt7b.h>
#include <Fonts/FreeSansBold24pt7b.h>


//#define DEBUG //uncomment to turn on debug mode, prints info to Serial Monitor

#ifdef DEBUG
#define DEBUG_PRINT(x)     Serial.print (x)
#define DEBUG_PRINTDEC(x)     Serial.print (x, DEC)
#define DEBUG_PRINTLN(x)  Serial.println (x)
#else
#define DEBUG_PRINT(x)
#define DEBUG_PRINTDEC(x)
#define DEBUG_PRINTLN(x)
#endif

// The FT6206 uses hardware I2C on the shield
Adafruit_FT6206 ts = Adafruit_FT6206();

// The display also uses hardware SPI, plus #9 & #10
#define TFT_CS 10
#define TFT_DC 9
Adafruit_ILI9341 tft = Adafruit_ILI9341(TFT_CS, TFT_DC);

RTC_PCF8523 rtc; // define the Real Time Clock object
const int chipSelect = 7; // for the data logging shield, changed from digital pin 10
#define redLEDpin 13
DateTime now;

//capacitive touch device
#ifndef _BV
#define _BV(bit) (1 << (bit))
#endif

// You can have up to 4 on one i2c bus but one is enough for testing!
Adafruit_MPR121 cap = Adafruit_MPR121();
Adafruit_MPR121 cap2 = Adafruit_MPR121();
Adafruit_MPR121 cap3 = Adafruit_MPR121();

// Keeps track of the last pins touched so we know when buttons are 'released'
uint16_t lasttouched1 = 0;
uint16_t currtouched1 = 0;
uint16_t lasttouched2 = 0;
uint16_t currtouched2 = 0;
uint16_t lasttouched3 = 0;
uint16_t currtouched3 = 0;

File logfile; // the logging file
char filename[] = "00000000.CSV";

TS_Point p;
extern uint8_t welcome_logo[];
extern uint8_t main_logo[];
extern uint8_t settings_icon[];
extern uint8_t settings_icon2[];
String E_side = "RIGHT";
String display_page = "main";
bool show_time = true;
bool eject_button;
int LOG_COUNTER;
int SYNC_COUNTER;
unsigned long currentMillis;
unsigned long previousMillis;
int syncTime;
int min_now;
int prev_min;
int curr_min;
int hour_now;
bool refresh_page = true;
int timeouts = 0;
bool ok;
int default_lights_on;
int default_lights_off;
int default_touch_threshold;
int default_release_threshold;
int default_LOG_INTERVAL;
int dafault_SYNC_INTERVAL;
bool default_auto_cal;
int default_auto_cal_time;
int default_auto_cal_sec_since_last_lick;
bool auto_cal_flag = true;
unsigned long cal_timer;
String logging[6];
bool log_LN = true;
bool log_LD = true;
bool log_BN = true;
bool log_BD = true;
bool log_BLN = true;
bool log_BLD = true;

//enough for 18 cages, 2 lickometers per cage
unsigned long LickDuration[36];
unsigned long LickNumber[36];
unsigned long BoutNumber[36];
unsigned long BoutDuration[36];
unsigned long BoutLickNumber[36];
unsigned long BoutLickDuration[36];
unsigned long Elapsedtime[36];
unsigned long total_LN[36];
int lick_bout_countdown[36];
unsigned long bout_timer[36];
unsigned long bout_start_timer[36];
bool in_bout[36];
bool licking[36];
unsigned long time_now[36];

// how many minutes between grabbing data and logging it.
int LOG_INTERVAL = 1; // IN MINUTES! between entries (reduce to take more/faster data)
// how many milliseconds before writing the logged data permanently to disk
int SYNC_INTERVAL = 10; // IN MINUTES! between calls to flush() - to write data to the card
unsigned long emergency_counter = 60050; //1 minute +50ms

// Typically touch threshold > release threshold for debouncing
// Decrease value to increase sensitivity
int touch_threshold = 2; //default = 12
int release_threshold = 0; //default = 6
bool auto_cal = false;
int auto_cal_time = 7;
int auto_cal_sec_since_last_lick = 60;

int lights_on = 6; //time of day (24hr time) housing lights turn ON
int lights_off = 18; //time of day (24hr time) houseing lights turn OFF

//==========================================================================================

void setup() {

  Serial.begin(115200);

  pinMode(5, OUTPUT); // set screen brightness pin as output
  analogWrite(5, 255);

  set_defaults(); //save coded settings as the default settings

  start_display(); //initialize display

  start_rtc(); //iniitialize rtc

  WelcomeScreen(); //display welcome screen at startup

  set_brightness(); //set screen brightness depending on light cycle

}

//==========================================================================================

void loop() {

  main_menu_page();

  recording_page();

  pause_page();

  SD_error_page();

  settings_page();

}
```
