## Extended Date - LIQ HD Arduino Source Code for "LIQ HD (Lick Instance Quantifier Home cage Device): An open-source tool for recording undisturbed two-bottle drinking behavior in a home cage environment": Logo.c

```
#include <avr/pgmspace.h>
// 'welcome_logo', 240x173px
const unsigned char  welcome_logo [] PROGMEM = {
  0x00, 0x00, 0x00, 0x00, 0x00, 0x00, 0x00, 0x00, 0x00, 0x00, 0x00, 0x00, 0x00, 0x00, 0x00, 0x00,
  0x00, 0x00, 0x00, 0x00, 0x00, 0x00, 0x00, 0x00, 0x00, 0x00, 0x00, 0x00, 0x00, 0x00, 0x00, 0x00,
  0x00, 0x00, 0x00, 0x00, 0x00, 0x00, 0x00, 0x00, 0x00, 0x00, 0x00, 0x00, 0x00, 0x00, 0x00, 0x00,
  0x00, 0x00, 0x00, 0x00, 0x00, 0x00, 0x00, 0x00, 0x00, 0x00, 0x00, 0x00, 0x00, 0x00, 0x00, 0x00,
  0x00, 0x00, 0x00, 0x00, 0x00, 0x00, 0x00, 0x00, 0x00, 0x00, 0x00, 0x00, 0x00, 0x00, 0x00, 0x00,
  0x00, 0x00, 0x00, 0x00, 0x00, 0x00, 0x00, 0x00, 0x00, 0x00, 0x00, 0x00, 0x00, 0x00, 0x00, 0x00,
  0x00, 0x00, 0x00, 0x00, 0x00, 0x00, 0x00, 0x00, 0x00, 0x00, 0x00, 0x00, 0x00, 0x00, 0x00, 0x00,
  0x00, 0x00, 0x00, 0x00, 0x00, 0x00, 0x00, 0x00, 0x00, 0x00, 0x00, 0x00, 0x00, 0x00, 0x00, 0x00,
  0x00, 0x00, 0x00, 0x00, 0x00, 0x00, 0x00, 0x00, 0x00, 0x00, 0x00, 0x00, 0x00, 0x00, 0x00, 0x00,
  0x00, 0x00, 0x00, 0x00, 0x00, 0x00, 0x00, 0x00, 0x00, 0x00, 0x00, 0x00, 0x00, 0x00, 0x00, 0x00,
  0x00, 0x00, 0x00, 0x00, 0x00, 0x00, 0x00, 0x00, 0x00, 0x00, 0x00, 0x00, 0x00, 0x00, 0x00, 0x00,
  0x00, 0x00, 0x00, 0x00, 0x00, 0x00, 0x00, 0x00, 0x00, 0x00, 0x00, 0x00, 0x00, 0x00, 0x00, 0x00,
  0x00, 0x00, 0x00, 0x00, 0x00, 0x00, 0x00, 0x00, 0x00, 0x00, 0x00, 0x00, 0x00, 0x00, 0x00, 0x00,
  0x00, 0x00, 0x00, 0x00, 0x00, 0x00, 0x00, 0x00, 0x00, 0x00, 0x00, 0x00, 0x00, 0x00, 0x00, 0x00,
  0x00, 0x00, 0x00, 0x00, 0x00, 0x00, 0x00, 0x00, 0x00, 0x00, 0x00, 0x00, 0x00, 0x00, 0x00, 0x00,
  0x00, 0x00, 0x00, 0x00, 0x00, 0x00, 0x00, 0x00, 0x00, 0x00, 0x00, 0x00, 0x00, 0x00, 0x00, 0x00,
  0x00, 0x00, 0x00, 0x00, 0x00, 0x00, 0x00, 0x00, 0x00, 0x00, 0x00, 0x00, 0x00, 0x00, 0x00, 0x00,
  0x00, 0x00, 0x00, 0x00, 0x00, 0x00, 0x00, 0x00, 0x00, 0x00, 0x00, 0x00, 0x00, 0x00, 0x00, 0x00,
  0x00, 0x00, 0x00, 0x00, 0x00, 0x00, 0x00, 0x00, 0x00, 0x00, 0x00, 0x00, 0x00, 0x00, 0x00, 0x00,
  0x00, 0x00, 0x00, 0x00, 0x00, 0x00, 0x00, 0x00, 0x00, 0x00, 0x00, 0x00, 0x00, 0x00, 0x00, 0x00,
  0x00, 0x00, 0x00, 0x00, 0x00, 0x00, 0x00, 0x00, 0x00, 0x00, 0x00, 0x00, 0x00, 0x00, 0x00, 0x00,
  0x00, 0x00, 0x00, 0x00, 0x00, 0x00, 0x00, 0x00, 0x00, 0x00, 0x00, 0x00, 0x00, 0x00, 0x00, 0x00,
  0x00, 0x00, 0x00, 0x00, 0x00, 0x00, 0x00, 0x00, 0x00, 0x00, 0x00, 0x00, 0x00, 0x00, 0x00, 0x00,
  0x00, 0x00, 0x00, 0x00, 0x00, 0x00, 0x00, 0x00, 0x00, 0x00, 0x00, 0x00, 0x00, 0x00, 0x00, 0x00,
  0x00, 0x00, 0x00, 0x00, 0x00, 0x00, 0x00, 0x00, 0x00, 0x00, 0x00, 0x00, 0x00, 0x00, 0x00, 0x00,
  0x00, 0x00, 0x00, 0x00, 0x00, 0x00, 0x00, 0x00, 0x00, 0x00, 0x00, 0x00, 0x00, 0x00, 0x00, 0x00,
  0x00, 0x00, 0x00, 0x00, 0x00, 0x00, 0x00, 0x00, 0x00, 0x00, 0x00, 0x00, 0x00, 0x00, 0x00, 0x00,
  0x00, 0x00, 0x00, 0x00, 0x00, 0x00, 0x00, 0x00, 0x00, 0x00, 0x00, 0x00, 0x00, 0x00, 0x00, 0x00,
  0x00, 0x00, 0x00, 0x00, 0x00, 0x00, 0x00, 0x00, 0x00, 0x00, 0x00, 0x00, 0x00, 0x00, 0x00, 0x00,
  0x00, 0x00, 0x00, 0x00, 0x00, 0x00, 0x00, 0x00, 0x00, 0x00, 0x00, 0x00, 0x00, 0x00, 0x00, 0x00,
  0x00, 0x00, 0x00, 0x00, 0x00, 0x00, 0x00, 0x00, 0x00, 0x00, 0x00, 0x00, 0x00, 0x00, 0x00, 0x00,
  0x00, 0x00, 0x00, 0x00, 0x00, 0x00, 0x00, 0x00, 0x00, 0x00, 0x00, 0x00, 0x00, 0x00, 0x00, 0x00,
  0x00, 0x00, 0x00, 0x00, 0x00, 0x00, 0x00, 0x00, 0x00, 0x00, 0x00, 0x00, 0x00, 0x00, 0x00, 0x00,
  0x00, 0x00, 0x00, 0x00, 0x00, 0x00, 0x00, 0x00, 0x00, 0x00, 0x00, 0x00, 0x00, 0x00, 0x00, 0x00,
  0x00, 0x00, 0x00, 0x00, 0x00, 0x00, 0x00, 0x00, 0x00, 0x00, 0x00, 0x00, 0x00, 0x00, 0x00, 0x00,
  0x00, 0x00, 0x00, 0x00, 0x00, 0x00, 0x00, 0x00, 0x00, 0x00, 0x00, 0x00, 0x00, 0x00, 0x00, 0x00,
  0x00, 0x00, 0x00, 0x00, 0x00, 0x00, 0x00, 0x00, 0x00, 0x00, 0x00, 0x00, 0x00, 0x00, 0x00, 0x00,
  0x00, 0x00, 0x00, 0x00, 0x00, 0x00, 0x00, 0x00, 0x00, 0x00, 0x00, 0x00, 0x00, 0x00, 0x00, 0x00,
  0x00, 0x00, 0x00, 0x00, 0x00, 0x00, 0x00, 0x00, 0x00, 0x00, 0x00, 0x00, 0x00, 0x00, 0x00, 0x00,
  0x00, 0x00, 0x00, 0x00, 0x00, 0x00, 0x00, 0x00, 0x00, 0x00, 0x00, 0x00, 0x00, 0x00, 0x00, 0x00,
  0x00, 0x00, 0x00, 0x00, 0x00, 0x00, 0x00, 0x00, 0x00, 0x00, 0x00, 0x00, 0x00, 0x00, 0x00, 0x00,
  0x00, 0x00, 0x00, 0x00, 0x00, 0x00, 0x00, 0x00, 0x00, 0x00, 0x00, 0x00, 0x00, 0x00, 0x00, 0x00,
  0x00, 0x00, 0x00, 0x00, 0x00, 0x00, 0x00, 0x00, 0x00, 0x00, 0x00, 0x00, 0x00, 0x00, 0x00, 0x00,
  0x00, 0x00, 0x00, 0x00, 0x00, 0x00, 0x00, 0x00, 0x00, 0x00, 0x00, 0x00, 0x00, 0x00, 0x00, 0x00,
  0x00, 0x00, 0x00, 0x00, 0x00, 0x00, 0x00, 0x00, 0x00, 0x00, 0x00, 0x00, 0x00, 0x00, 0x00, 0x00,
  0x00, 0x00, 0x00, 0x00, 0x00, 0x00, 0x00, 0x00, 0x00, 0x00, 0x00, 0x00, 0x00, 0x00, 0x00, 0x00,
  0x00, 0x00, 0x00, 0x00, 0x00, 0x00, 0x00, 0x00, 0x00, 0x00, 0x00, 0x00, 0x00, 0x00, 0x00, 0x00,
  0x00, 0x00, 0x00, 0x00, 0x00, 0x00, 0x00, 0x00, 0x00, 0x00, 0x00, 0x00, 0x00, 0x00, 0x00, 0x00,
  0x00, 0x00, 0x00, 0x00, 0x00, 0x00, 0x00, 0x00, 0x00, 0x00, 0x00, 0x00, 0x00, 0x00, 0x00, 0x00,
  0x00, 0x00, 0x00, 0x00, 0x00, 0x00, 0x00, 0x00, 0x00, 0x00, 0x00, 0x00, 0x00, 0x00, 0x00, 0x00,
  0x00, 0x00, 0x00, 0x00, 0x00, 0x00, 0x00, 0x00, 0x00, 0x00, 0x00, 0x00, 0x00, 0x00, 0x00, 0x00,
  0x00, 0x00, 0x00, 0x00, 0x00, 0x00, 0x00, 0x00, 0x00, 0x00, 0x00, 0x00, 0x00, 0x00, 0x00, 0x00,
  0x00, 0x00, 0x00, 0x00, 0x00, 0x00, 0x00, 0x00, 0x00, 0x00, 0x00, 0x00, 0x00, 0x00, 0x00, 0x00,
  0x00, 0x00, 0x00, 0x00, 0x00, 0x00, 0x00, 0x00, 0x00, 0x00, 0x00, 0x00, 0x00, 0x00, 0x00, 0x00,
  0x00, 0x00, 0x00, 0x00, 0x00, 0x00, 0x00, 0x00, 0x00, 0x00, 0x00, 0x00, 0x00, 0x00, 0x00, 0x00,
  0x00, 0x00, 0x00, 0x00, 0x00, 0x00, 0x00, 0x00, 0x00, 0x00, 0x00, 0x00, 0x00, 0x00, 0x00, 0x00,
  0x00, 0x00, 0x00, 0x00, 0x00, 0x00, 0x00, 0x00, 0x0f, 0xe0, 0x00, 0x01, 0xfc, 0x00, 0x0f, 0xff,
  0xf8, 0x00, 0x00, 0x00, 0x01, 0xfc, 0x00, 0x03, 0xf8, 0x0f, 0xff, 0xff, 0xf0, 0x00, 0x00, 0x00,
  0x00, 0x00, 0x00, 0x00, 0x00, 0x00, 0x0f, 0xe0, 0x00, 0x01, 0xfc, 0x00, 0x3f, 0xff, 0xfe, 0x00,
  0x00, 0x00, 0x01, 0xfc, 0x00, 0x03, 0xf8, 0x0f, 0xff, 0xff, 0xfc, 0x00, 0x00, 0x00, 0x00, 0x00,
  0x00, 0x00, 0x00, 0x00, 0x0f, 0xe0, 0x00, 0x01, 0xfc, 0x00, 0x7f, 0xff, 0xff, 0x00, 0x00, 0x00,
  0x01, 0xfc, 0x00, 0x03, 0xf8, 0x0f, 0xff, 0xff, 0xfe, 0x00, 0x00, 0x00, 0x00, 0x00, 0x00, 0x00,
  0x00, 0x00, 0x0f, 0xe0, 0x00, 0x01, 0xfc, 0x00, 0xff, 0xff, 0xff, 0x80, 0x00, 0x00, 0x01, 0xfc,
  0x00, 0x03, 0xf8, 0x0f, 0xff, 0xff, 0xff, 0x00, 0x00, 0x00, 0x00, 0x00, 0x00, 0x00, 0x00, 0x00,
  0x0f, 0xe0, 0x00, 0x01, 0xfc, 0x01, 0xff, 0xff, 0xff, 0xc0, 0x00, 0x00, 0x01, 0xfc, 0x00, 0x03,
  0xf8, 0x0f, 0xff, 0xff, 0xff, 0x80, 0x00, 0x00, 0x00, 0x00, 0x00, 0x00, 0x00, 0x00, 0x0f, 0xe0,
  0x00, 0x01, 0xfc, 0x03, 0xff, 0xff, 0xff, 0xe0, 0x00, 0x00, 0x01, 0xfc, 0x00, 0x03, 0xf8, 0x0f,
  0xff, 0xff, 0xff, 0xc0, 0x00, 0x00, 0x00, 0x00, 0x00, 0x00, 0x00, 0x00, 0x0f, 0xe0, 0x00, 0x01,
  0xfc, 0x03, 0xff, 0xff, 0xff, 0xe0, 0x00, 0x00, 0x01, 0xfc, 0x00, 0x03, 0xf8, 0x0f, 0xff, 0xff,
  0xff, 0xc0, 0x00, 0x00, 0x00, 0x00, 0x00, 0x00, 0x00, 0x00, 0x0f, 0xe0, 0x00, 0x01, 0xfc, 0x03,
  0xf8, 0x00, 0x0f, 0xe0, 0x00, 0x00, 0x01, 0xfc, 0x00, 0x03, 0xf8, 0x0f, 0xe0, 0x00, 0x1f, 0xc0,
  0x00, 0x00, 0x00, 0x00, 0x00, 0x00, 0x00, 0x00, 0x0f, 0xe0, 0x00, 0x01, 0xfc, 0x07, 0xf0, 0x00,
  0x07, 0xf0, 0x00, 0x00, 0x01, 0xfc, 0x00, 0x03, 0xf8, 0x0f, 0xe0, 0x00, 0x0f, 0xe0, 0x00, 0x00,
  0x00, 0x00, 0x00, 0x00, 0x00, 0x00, 0x0f, 0xe0, 0x00, 0x01, 0xfc, 0x07, 0xf0, 0x00, 0x07, 0xf0,
  0x00, 0x00, 0x01, 0xfc, 0x00, 0x03, 0xf8, 0x0f, 0xe0, 0x00, 0x0f, 0xe0, 0x00, 0x00, 0x00, 0x00,
  0x00, 0x00, 0x00, 0x00, 0x0f, 0xe0, 0x00, 0x01, 0xfc, 0x07, 0xf0, 0x00, 0x07, 0xf0, 0x00, 0x00,
  0x01, 0xfc, 0x00, 0x03, 0xf8, 0x0f, 0xe0, 0x00, 0x0f, 0xe0, 0x00, 0x00, 0x00, 0x00, 0x00, 0x00,
  0x00, 0x00, 0x0f, 0xe0, 0x00, 0x01, 0xfc, 0x07, 0xf0, 0x00, 0x07, 0xf0, 0x00, 0x00, 0x01, 0xfc,
  0x00, 0x03, 0xf8, 0x0f, 0xe0, 0x00, 0x0f, 0xe0, 0x00, 0x00, 0x00, 0x00, 0x00, 0x00, 0x00, 0x00,
  0x0f, 0xe0, 0x00, 0x01, 0xfc, 0x07, 0xf0, 0x00, 0x07, 0xf0, 0x00, 0x00, 0x01, 0xfc, 0x00, 0x03,
  0xf8, 0x0f, 0xe0, 0x00, 0x0f, 0xe0, 0x00, 0x00, 0x00, 0x00, 0x00, 0x00, 0x00, 0x00, 0x0f, 0xe0,
  0x00, 0x01, 0xfc, 0x07, 0xf0, 0x00, 0x07, 0xf0, 0x00, 0x00, 0x01, 0xfc, 0x00, 0x03, 0xf8, 0x0f,
  0xe0, 0x00, 0x0f, 0xe0, 0x00, 0x00, 0x00, 0x00, 0x00, 0x00, 0x00, 0x00, 0x0f, 0xe0, 0x00, 0x01,
  0xfc, 0x07, 0xf0, 0x00, 0x07, 0xf0, 0x00, 0x00, 0x01, 0xfc, 0x00, 0x03, 0xf8, 0x0f, 0xe0, 0x00,
  0x0f, 0xe0, 0x00, 0x00, 0x00, 0x00, 0x00, 0x00, 0x00, 0x00, 0x0f, 0xe0, 0x00, 0x01, 0xfc, 0x07,
  0xf0, 0x00, 0x07, 0xf0, 0x00, 0x00, 0x01, 0xfc, 0x00, 0x03, 0xf8, 0x0f, 0xe0, 0x00, 0x0f, 0xe0,
  0x00, 0x00, 0x00, 0x00, 0x00, 0x00, 0x00, 0x00, 0x0f, 0xe0, 0x00, 0x01, 0xfc, 0x07, 0xf0, 0x00,
  0x07, 0xf0, 0x00, 0x00, 0x01, 0xfc, 0x00, 0x03, 0xf8, 0x0f, 0xe0, 0x00, 0x0f, 0xe0, 0x00, 0x00,
  0x00, 0x00, 0x00, 0x00, 0x00, 0x00, 0x0f, 0xe0, 0x00, 0x01, 0xfc, 0x07, 0xf0, 0x00, 0x07, 0xf0,
  0x00, 0x00, 0x01, 0xfc, 0x00, 0x03, 0xf8, 0x0f, 0xe0, 0x00, 0x0f, 0xe0, 0x00, 0x00, 0x00, 0x00,
  0x00, 0x00, 0x00, 0x00, 0x0f, 0xe0, 0x00, 0x01, 0xfc, 0x07, 0xf0, 0x00, 0x07, 0xf0, 0x00, 0x00,
  0x01, 0xfc, 0x00, 0x03, 0xf8, 0x0f, 0xe0, 0x00, 0x0f, 0xe0, 0x00, 0x00, 0x00, 0x00, 0x00, 0x00,
  0x00, 0x00, 0x0f, 0xe0, 0x00, 0x01, 0xfc, 0x07, 0xf0, 0x00, 0x07, 0xf0, 0x00, 0x00, 0x01, 0xfc,
  0x00, 0x03, 0xf8, 0x0f, 0xe0, 0x00, 0x0f, 0xe0, 0x00, 0x00, 0x00, 0x00, 0x00, 0x00, 0x00, 0x00,
  0x0f, 0xe0, 0x00, 0x01, 0xfc, 0x07, 0xf0, 0x00, 0x07, 0xf0, 0x00, 0x00, 0x01, 0xfc, 0x00, 0x03,
  0xf8, 0x0f, 0xe0, 0x00, 0x0f, 0xe0, 0x00, 0x00, 0x00, 0x00, 0x00, 0x00, 0x00, 0x00, 0x0f, 0xe0,
  0x00, 0x01, 0xfc, 0x07, 0xf0, 0x00, 0x07, 0xf0, 0x00, 0x00, 0x01, 0xfc, 0x00, 0x03, 0xf8, 0x0f,
  0xe0, 0x00, 0x0f, 0xe0, 0x00, 0x00, 0x00, 0x00, 0x00, 0x00, 0x00, 0x00, 0x0f, 0xe0, 0x00, 0x01,
  0xfc, 0x07, 0xf0, 0x00, 0x07, 0xf0, 0x00, 0x00, 0x01, 0xfc, 0x00, 0x03, 0xf8, 0x0f, 0xe0, 0x00,
  0x0f, 0xe0, 0x00, 0x00, 0x00, 0x00, 0x00, 0x00, 0x00, 0x00, 0x0f, 0xe0, 0x00, 0x01, 0xfc, 0x07,
  0xf0, 0x00, 0x07, 0xf0, 0x00, 0x00, 0x01, 0xfc, 0x00, 0x03, 0xf8, 0x0f, 0xe0, 0x00, 0x0f, 0xe0,
  0x00, 0x00, 0x00, 0x00, 0x00, 0x00, 0x00, 0x00, 0x0f, 0xe0, 0x00, 0x01, 0xfc, 0x07, 0xf0, 0x00,
  0x07, 0xf0, 0x00, 0x00, 0x01, 0xfc, 0x00, 0x03, 0xf8, 0x0f, 0xe0, 0x00, 0x0f, 0xe0, 0x00, 0x00,
  0x00, 0x00, 0x00, 0x00, 0x00, 0x00, 0x0f, 0xe0, 0x00, 0x01, 0xfc, 0x07, 0xf0, 0x00, 0x07, 0xf0,
  0x00, 0x00, 0x01, 0xfc, 0x00, 0x03, 0xf8, 0x0f, 0xe0, 0x00, 0x0f, 0xe0, 0x00, 0x00, 0x00, 0x00,
  0x00, 0x00, 0x00, 0x00, 0x0f, 0xe0, 0x00, 0x01, 0xfc, 0x07, 0xf0, 0x00, 0x07, 0xf0, 0x00, 0x00,
  0x01, 0xfc, 0x00, 0x03, 0xf8, 0x0f, 0xe0, 0x00, 0x0f, 0xe0, 0x00, 0x00, 0x00, 0x00, 0x00, 0x00,
  0x00, 0x00, 0x0f, 0xe0, 0x00, 0x01, 0xfc, 0x07, 0xf0, 0x00, 0x07, 0xf0, 0x00, 0x00, 0x01, 0xfc,
  0x00, 0x03, 0xf8, 0x0f, 0xe0, 0x00, 0x0f, 0xe0, 0x00, 0x00, 0x00, 0x00, 0x00, 0x00, 0x00, 0x00,
  0x0f, 0xe0, 0x00, 0x01, 0xfc, 0x07, 0xf0, 0x00, 0x07, 0xf0, 0x00, 0x00, 0x01, 0xfc, 0x00, 0x03,
  0xf8, 0x0f, 0xe0, 0x00, 0x0f, 0xe0, 0x00, 0x00, 0x00, 0x00, 0x00, 0x00, 0x00, 0x00, 0x0f, 0xe0,
  0x00, 0x01, 0xfc, 0x07, 0xf0, 0x00, 0x07, 0xf0, 0x00, 0x00, 0x01, 0xfc, 0x00, 0x03, 0xf8, 0x0f,
  0xe0, 0x00, 0x0f, 0xe0, 0x00, 0x00, 0x00, 0x00, 0x00, 0x00, 0x00, 0x00, 0x0f, 0xe0, 0x00, 0x01,
  0xfc, 0x07, 0xf0, 0x00, 0x07, 0xf0, 0x00, 0x00, 0x01, 0xfc, 0x00, 0x03, 0xf8, 0x0f, 0xe0, 0x00,
  0x0f, 0xe0, 0x00, 0x00, 0x00, 0x00, 0x00, 0x00, 0x00, 0x00, 0x0f, 0xe0, 0x00, 0x01, 0xfc, 0x07,
  0xf0, 0x00, 0x07, 0xf0, 0x00, 0x00, 0x0f, 0xff, 0xff, 0xff, 0xf8, 0x0f, 0xe0, 0x00, 0x0f, 0xe0,
  0x00, 0x00, 0x00, 0x00, 0x00, 0x00, 0x00, 0x00, 0x0f, 0xe0, 0x00, 0x01, 0xfc, 0x07, 0xf0, 0x00,
  0x07, 0xf0, 0x00, 0x00, 0x0f, 0xff, 0xff, 0xff, 0xf8, 0x0f, 0xe0, 0x00, 0x0f, 0xe0, 0x00, 0x00,
  0x00, 0x00, 0x00, 0x00, 0x00, 0x00, 0x0f, 0xe0, 0x00, 0x01, 0xfc, 0x07, 0xf0, 0x00, 0x07, 0xf0,
  0x00, 0x00, 0x0f, 0xff, 0xff, 0xff, 0xf8, 0x0f, 0xe0, 0x00, 0x0f, 0xe0, 0x00, 0x00, 0x00, 0x00,
  0x00, 0x00, 0x00, 0x00, 0x0f, 0xe0, 0x00, 0x01, 0xfc, 0x07, 0xf0, 0x00, 0x07, 0xf0, 0x00, 0x00,
  0x0f, 0xff, 0xff, 0xff, 0xf8, 0x0f, 0xe0, 0x00, 0x0f, 0xe0, 0x00, 0x00, 0x00, 0x00, 0x00, 0x00,
  0x00, 0x00, 0x0f, 0xe0, 0x00, 0x01, 0xfc, 0x07, 0xf0, 0x00, 0x07, 0xf0, 0x00, 0x00, 0x0f, 0xff,
  0xff, 0xff, 0xf8, 0x0f, 0xe0, 0x00, 0x0f, 0xe0, 0x00, 0x00, 0x00, 0x00, 0x00, 0x00, 0x00, 0x00,
  0x0f, 0xe0, 0x00, 0x01, 0xfc, 0x07, 0xf0, 0x00, 0x07, 0xf0, 0x00, 0x00, 0x0f, 0xff, 0xff, 0xff,
  0xf8, 0x0f, 0xe0, 0x00, 0x0f, 0xe0, 0x00, 0x00, 0x00, 0x00, 0x00, 0x00, 0x00, 0x00, 0x0f, 0xe0,
  0x00, 0x01, 0xfc, 0x07, 0xf0, 0x00, 0x07, 0xf0, 0x00, 0x00, 0x0f, 0xff, 0xff, 0xff, 0xf8, 0x0f,
  0xe0, 0x00, 0x0f, 0xe0, 0x00, 0x00, 0x00, 0x00, 0x00, 0x00, 0x00, 0x00, 0x0f, 0xe0, 0x00, 0x01,
  0xfc, 0x07, 0xf0, 0x00, 0x07, 0xf0, 0x00, 0x00, 0x01, 0xfc, 0x00, 0x03, 0xf8, 0x0f, 0xe0, 0x00,
  0x0f, 0xe0, 0x00, 0x00, 0x00, 0x00, 0x00, 0x00, 0x00, 0x00, 0x0f, 0xe0, 0x00, 0x01, 0xfc, 0x07,
  0xf0, 0x00, 0x07, 0xf0, 0x00, 0x00, 0x01, 0xfc, 0x00, 0x03, 0xf8, 0x0f, 0xe0, 0x00, 0x0f, 0xe0,
  0x00, 0x00, 0x00, 0x00, 0x00, 0x00, 0x00, 0x00, 0x0f, 0xe0, 0x00, 0x01, 0xfc, 0x07, 0xf0, 0x00,
  0x07, 0xf0, 0x00, 0x00, 0x01, 0xfc, 0x00, 0x03, 0xf8, 0x0f, 0xe0, 0x00, 0x0f, 0xe0, 0x00, 0x00,
  0x00, 0x00, 0x00, 0x00, 0x00, 0x00, 0x0f, 0xe0, 0x00, 0x01, 0xfc, 0x07, 0xf0, 0x00, 0x07, 0xf0,
  0x00, 0x00, 0x01, 0xfc, 0x00, 0x03, 0xf8, 0x0f, 0xe0, 0x00, 0x0f, 0xe0, 0x00, 0x00, 0x00, 0x00,
  0x00, 0x00, 0x00, 0x00, 0x0f, 0xe0, 0x00, 0x01, 0xfc, 0x07, 0xf0, 0x00, 0x07, 0xf0, 0x00, 0x00,
  0x01, 0xfc, 0x00, 0x03, 0xf8, 0x0f, 0xe0, 0x00, 0x0f, 0xe0, 0x00, 0x00, 0x00, 0x00, 0x00, 0x00,
  0x00, 0x00, 0x0f, 0xe0, 0x00, 0x01, 0xfc, 0x07, 0xf8, 0x00, 0x0f, 0xe0, 0x00, 0x00, 0x01, 0xfc,
  0x00, 0x03, 0xf8, 0x0f, 0xe0, 0x00, 0x1f, 0xc0, 0x00, 0x00, 0x00, 0x00, 0x00, 0x00, 0x00, 0x00,
  0x0f, 0xff, 0xff, 0x01, 0xfc, 0x03, 0xff, 0xff, 0xff, 0xe0, 0x00, 0x00, 0x01, 0xfc, 0x00, 0x03,
  0xf8, 0x0f, 0xff, 0xff, 0xff, 0xc0, 0x00, 0x00, 0x00, 0x00, 0x00, 0x00, 0x00, 0x00, 0x0f, 0xff,
  0xff, 0x01, 0xfc, 0x03, 0xff, 0xff, 0xff, 0xe0, 0x00, 0x00, 0x01, 0xfc, 0x00, 0x03, 0xf8, 0x0f,
  0xff, 0xff, 0xff, 0xc0, 0x00, 0x00, 0x00, 0x00, 0x00, 0x00, 0x00, 0x00, 0x0f, 0xff, 0xff, 0x01,
  0xfc, 0x01, 0xff, 0xff, 0xff, 0xc0, 0x00, 0x00, 0x01, 0xfc, 0x00, 0x03, 0xf8, 0x0f, 0xff, 0xff,
  0xff, 0x80, 0x00, 0x00, 0x00, 0x00, 0x00, 0x00, 0x00, 0x00, 0x0f, 0xff, 0xff, 0x01, 0xfc, 0x00,
  0xff, 0xff, 0xff, 0x80, 0x00, 0x00, 0x01, 0xfc, 0x00, 0x03, 0xf8, 0x0f, 0xff, 0xff, 0xff, 0x00,
  0x00, 0x00, 0x00, 0x00, 0x00, 0x00, 0x00, 0x00, 0x0f, 0xff, 0xff, 0x01, 0xfc, 0x00, 0x7f, 0xff,
  0xff, 0x00, 0x00, 0x00, 0x01, 0xfc, 0x00, 0x03, 0xf8, 0x0f, 0xff, 0xff, 0xfe, 0x00, 0x00, 0x00,
  0x00, 0x00, 0x00, 0x00, 0x00, 0x00, 0x0f, 0xff, 0xff, 0x01, 0xfc, 0x00, 0x3f, 0xff, 0xfe, 0x00,
  0x00, 0x00, 0x01, 0xfc, 0x00, 0x03, 0xf8, 0x0f, 0xff, 0xff, 0xfc, 0x00, 0x00, 0x00, 0x00, 0x00,
  0x00, 0x00, 0x00, 0x00, 0x0f, 0xff, 0xff, 0x01, 0xfc, 0x00, 0x0f, 0xff, 0xf8, 0x00, 0x00, 0x00,
  0x01, 0xfc, 0x00, 0x03, 0xf8, 0x0f, 0xff, 0xff, 0xf0, 0x00, 0x00, 0x00, 0x00, 0x00, 0x00, 0x00,
  0x00, 0x00, 0x00, 0x00, 0x00, 0x00, 0x00, 0x00, 0x00, 0x3f, 0xf0, 0x00, 0x00, 0x00, 0x00, 0x00,
  0x00, 0x00, 0x00, 0x00, 0x00, 0x00, 0x00, 0x00, 0x00, 0x00, 0x00, 0x00, 0x00, 0x00, 0x00, 0x00,
  0x00, 0x00, 0x00, 0x00, 0x00, 0x00, 0x00, 0x1f, 0xf8, 0x00, 0x00, 0x00, 0x00, 0x00, 0x00, 0x00,
  0x00, 0x00, 0x00, 0x00, 0x00, 0x00, 0x00, 0x00, 0x00, 0x00, 0x00, 0x00, 0x00, 0x00, 0x00, 0x00,
  0x00, 0x00, 0x00, 0x00, 0x00, 0x0f, 0xfc, 0x00, 0x00, 0x00, 0x00, 0x00, 0x00, 0x00, 0x00, 0x00,
  0x00, 0x00, 0x00, 0x00, 0x00, 0x00, 0x00, 0x00, 0x00, 0x00, 0x00, 0x00, 0x00, 0x00, 0x00, 0x00,
  0x00, 0x00, 0x00, 0x07, 0xfc, 0x00, 0x00, 0x00, 0x00, 0x00, 0x00, 0x00, 0x00, 0x00, 0x00, 0x00,
  0x00, 0x00, 0x00, 0x00, 0x00, 0x00, 0x00, 0x00, 0x00, 0x00, 0x00, 0x00, 0x00, 0x00, 0x00, 0x00,
  0x00, 0x03, 0xf8, 0x00, 0x00, 0x00, 0x00, 0x00, 0x00, 0x00, 0x00, 0x00, 0x00, 0x00, 0x00, 0x00,
  0x00, 0x00, 0x00, 0x00, 0x00, 0x00, 0x00, 0x00, 0x00, 0x00, 0x00, 0x00, 0x00, 0x00, 0x00, 0x01,
  0xf0, 0x00, 0x00, 0x00, 0x00, 0x00, 0x00, 0x00, 0x00, 0x00, 0x00, 0x00, 0x00, 0x00, 0x00, 0x00,
  0x00, 0x00, 0x00, 0x00, 0x00, 0x00, 0x00, 0x00, 0x00, 0x00, 0x00, 0x00, 0x00, 0x00, 0xe0, 0x00,
  0x00, 0x00, 0x00, 0x00, 0x00, 0x00, 0x00, 0x00, 0x00, 0x00, 0x00, 0x00, 0x00, 0x00, 0x00, 0x00,
  0x00, 0x00, 0x00, 0x00, 0x00, 0x00, 0x00, 0x00, 0x00, 0x00, 0x00, 0x00, 0x40, 0x00, 0x00, 0x00,
  0x00, 0x00, 0x00, 0x00, 0x00, 0x00, 0x00, 0x00, 0x00, 0x00, 0x00, 0x00, 0x00, 0x00, 0x00, 0x00,
  0x00, 0x00, 0x00, 0x00, 0x00, 0x00, 0x00, 0x00, 0x00, 0x00, 0x00, 0x00, 0x00, 0x00, 0x00, 0x00,
  0x00, 0x00, 0x00, 0x00, 0x00, 0x00, 0x00, 0x00, 0x00, 0x00, 0x00, 0x00, 0x00, 0x00, 0x00, 0x00,
  0x00, 0x00, 0x00, 0x00, 0x00, 0x00, 0x00, 0x00, 0x00, 0x00, 0x00, 0x00, 0x00, 0x00, 0x00, 0x00,
  0x00, 0x00, 0x00, 0x00, 0x00, 0x00, 0x00, 0x00, 0x00, 0x00, 0x00, 0x00, 0x00, 0x00, 0x00, 0x00,
  0x00, 0x00, 0x00, 0x00, 0x00, 0x00, 0x00, 0x00, 0x00, 0x00, 0x00, 0x00, 0x00, 0x00, 0x00, 0x00,
  0x00, 0x00, 0x00, 0x00, 0x00, 0x00, 0x00, 0x00, 0x00, 0x00, 0x00, 0x00, 0x00, 0x00, 0x00, 0x00,
  0x00, 0x00, 0x00, 0x00, 0x00, 0x00, 0x00, 0x00, 0x00, 0x00, 0x00, 0x00, 0x00, 0x00, 0x00, 0x00,
  0x00, 0x00, 0x00, 0x00, 0x00, 0x00, 0x00, 0x00, 0x00, 0x00, 0x00, 0x00, 0x00, 0x00, 0x00, 0x00,
  0x00, 0x00, 0x00, 0x00, 0x00, 0x00, 0x00, 0x00, 0x00, 0x00, 0x00, 0x00, 0x00, 0x00, 0x00, 0x00,
  0x00, 0x00, 0x00, 0x00, 0x00, 0x00, 0x00, 0x00, 0x00, 0x00, 0x00, 0x00, 0x00, 0x00, 0x00, 0x00,
  0x00, 0x00, 0x00, 0x00, 0x00, 0x00, 0x00, 0x00, 0x00, 0x00, 0x00, 0x00, 0x00, 0x00, 0x00, 0x00,
  0x00, 0x00, 0x00, 0x00, 0x00, 0x00, 0x00, 0x00, 0x00, 0x00, 0x00, 0x00, 0x00, 0x00, 0x00, 0x00,
  0x00, 0x00, 0x00, 0x00, 0x00, 0x00, 0x00, 0x00, 0x00, 0x00, 0x00, 0x00, 0x00, 0x00, 0x00, 0x00,
  0x00, 0x00, 0x00, 0x00, 0x00, 0x00, 0x00, 0x00, 0x00, 0x00, 0x00, 0x00, 0x00, 0x00, 0x00, 0x00,
  0x00, 0x00, 0x00, 0x00, 0x00, 0x00, 0x00, 0x00, 0x00, 0x00, 0x00, 0x00, 0x00, 0x00, 0x00, 0x00,
  0x00, 0x00, 0x00, 0x00, 0x00, 0x00, 0x00, 0x00, 0x00, 0x00, 0x00, 0x00, 0x00, 0x00, 0x00, 0x00,
  0x00, 0x00, 0x00, 0x00, 0x00, 0x00, 0x00, 0x00, 0x00, 0x00, 0x00, 0x00, 0x00, 0x00, 0x00, 0x00,
  0x00, 0x00, 0x00, 0x00, 0x00, 0x00, 0x00, 0x00, 0x00, 0x00, 0x00, 0x00, 0x00, 0x00, 0x00, 0x00,
  0x00, 0x00, 0x00, 0x00, 0x00, 0x00, 0x00, 0x00, 0x00, 0x00, 0x00, 0x00, 0x00, 0x00, 0x00, 0x00,
  0x00, 0x00, 0x00, 0x00, 0x00, 0x00, 0x00, 0x00, 0x00, 0x00, 0x00, 0x00, 0x00, 0x00, 0x00, 0x00,
  0x00, 0x00, 0x00, 0x00, 0x00, 0x00, 0x00, 0x00, 0x00, 0x00, 0x00, 0x00, 0x00, 0x00, 0x00, 0x00,
  0x00, 0x00, 0x00, 0x00, 0x00, 0x00, 0x00, 0x00, 0x00, 0x00, 0x00, 0x00, 0x00, 0x00, 0x00, 0x00,
  0x00, 0x00, 0x00, 0x00, 0x00, 0x00, 0x00, 0x00, 0x00, 0x00, 0x00, 0x00, 0x00, 0x00, 0x00, 0x00,
  0x00, 0x00, 0x00, 0x00, 0x00, 0x00, 0x00, 0x00, 0x00, 0x00, 0x00, 0x00, 0x00, 0x00, 0x00, 0x00,
  0x00, 0x00, 0x00, 0x00, 0x00, 0x00, 0x00, 0x00, 0x00, 0x00, 0x00, 0x00, 0x00, 0x00, 0x00, 0x00,
  0x00, 0x00, 0x00, 0x00, 0x00, 0x00, 0x00, 0x00, 0x00, 0x00, 0x00, 0x00, 0x00, 0x00, 0x00, 0x00,
  0x00, 0x00, 0x00, 0x00, 0x00, 0x00, 0x00, 0x00, 0x00, 0x00, 0x00, 0x00, 0x00, 0x00, 0x00, 0x00,
  0x00, 0x00, 0x00, 0x00, 0x00, 0x00, 0x00, 0x00, 0x00, 0x00, 0x00, 0x00, 0x00, 0x00, 0x00, 0x00,
  0x00, 0x00, 0x00, 0x00, 0x00, 0x00, 0x00, 0x00, 0x00, 0x00, 0x00, 0x00, 0x00, 0x00, 0x00, 0x00,
  0x00, 0x00, 0x00, 0x00, 0x00, 0x00, 0x00, 0x00, 0x00, 0x00, 0x00, 0x00, 0x00, 0x00, 0x00, 0x00,
  0x00, 0x00, 0x00, 0x00, 0x00, 0x00, 0x00, 0x00, 0x00, 0x00, 0x00, 0x00, 0x00, 0x00, 0x00, 0x00,
  0x00, 0x00, 0x00, 0x00, 0x00, 0x00, 0x00, 0x00, 0x00, 0x00, 0x00, 0x00, 0x00, 0x00, 0x00, 0x00,
  0x00, 0x00, 0x00, 0x00, 0x00, 0x00, 0x00, 0x00, 0x00, 0x00, 0x00, 0x00, 0x00, 0x00, 0x00, 0x00,
  0x00, 0x00, 0x00, 0x00, 0x00, 0x00, 0x00, 0x00, 0x00, 0x00, 0x00, 0x00, 0x00, 0x00, 0x00, 0x00,
  0x00, 0x00, 0x00, 0x00, 0x00, 0x00, 0x00, 0x00, 0x00, 0x00, 0x00, 0x00, 0x00, 0x00, 0x00, 0x00,
  0x00, 0x00, 0x00, 0x00, 0x00, 0x00, 0x00, 0x00, 0x00, 0x00, 0x00, 0x00, 0x00, 0x00, 0x00, 0x00,
  0x00, 0x00, 0x00, 0x00, 0x00, 0x00, 0x00, 0x00, 0x00, 0x00, 0x00, 0x00, 0x00, 0x00, 0x00, 0x00,
  0x00, 0x00, 0x00, 0x00, 0x00, 0x00, 0x00, 0x00, 0x00, 0x00, 0x00, 0x00, 0x00, 0x00, 0x00, 0x00,
  0x00, 0x00, 0x00, 0x00, 0x00, 0x00, 0x00, 0x00, 0x00, 0x00, 0x00, 0x00, 0x00, 0x00, 0x00, 0x00,
  0x00, 0x00, 0x00, 0x00, 0x00, 0x00, 0x00, 0x00, 0x00, 0x00, 0x00, 0x00, 0x00, 0x00, 0x00, 0x00,
  0x00, 0x00, 0x00, 0x00, 0x00, 0x00, 0x00, 0x00, 0x00, 0x00, 0x00, 0x00, 0x00, 0x00, 0x00, 0x00,
  0x00, 0x00, 0x00, 0x00, 0x00, 0x00, 0x00, 0x00, 0x00, 0x00, 0x00, 0x00, 0x00, 0x00, 0x00, 0x00,
  0x00, 0x00, 0x00, 0x00, 0x00, 0x00, 0x00, 0x00, 0x00, 0x00, 0x00, 0x00, 0x00, 0x00, 0x00, 0x00,
  0x00, 0x00, 0x00, 0x00, 0x00, 0x00, 0x00, 0x00, 0x00, 0x00, 0x00, 0x00, 0x00, 0x00, 0x00, 0x00,
  0x00, 0x00, 0x00, 0x00, 0x00, 0x00, 0x00, 0x00, 0x00, 0x00, 0x00, 0x00, 0x00, 0x00, 0x00, 0x00,
  0x00, 0x00, 0x00, 0x00, 0x00, 0x00, 0x00, 0x00, 0x00, 0x00, 0x00, 0x00, 0x00, 0x00, 0x00, 0x00,
  0x00, 0x00, 0x00, 0x00, 0x00, 0x00, 0x00, 0x00, 0x00, 0x00, 0x00, 0x00, 0x00, 0x00, 0x00, 0x00,
  0x00, 0x00, 0x00, 0x00, 0x00, 0x00, 0x00, 0x00, 0x00, 0x00, 0x00, 0x00, 0x00, 0x00, 0x00, 0x00,
  0x00, 0x00, 0x00, 0x00, 0x00, 0x00, 0x00, 0x00, 0x00, 0x00, 0x00, 0x00, 0x00, 0x00, 0x00, 0x00,
  0x00, 0x00, 0x00, 0x00, 0x00, 0x00, 0x00, 0x00, 0x00, 0x00, 0x00, 0x00, 0x00, 0x00, 0x00, 0x00,
  0x00, 0x00, 0x00, 0x00, 0x00, 0x00, 0x00, 0x00, 0x00, 0x00, 0x00, 0x00, 0x00, 0x00, 0x00, 0x00,
  0x00, 0x00, 0x00, 0x00, 0x00, 0x00, 0x00, 0x00, 0x00, 0x00, 0x00, 0x00, 0x00, 0x00, 0x00, 0x00,
  0x00, 0x00, 0x00, 0x00, 0x00, 0x00, 0x00, 0x00, 0x00, 0x00, 0x00, 0x00, 0x00, 0x00, 0x00, 0x00,
  0x00, 0x00, 0x00, 0x00, 0x00, 0x00, 0x00, 0x00, 0x00, 0x00, 0x00, 0x00, 0x00, 0x00, 0x00, 0x00,
  0x00, 0x00, 0x00, 0x00, 0x00, 0x00, 0x00, 0x00, 0x00, 0x00, 0x00, 0x00, 0x00, 0x00, 0x00, 0x00,
  0x00, 0x00, 0x00, 0x00, 0x00, 0x00, 0x00, 0x00, 0x00, 0x00, 0x00, 0x00, 0x00, 0x00, 0x00, 0x00,
  0x00, 0x00, 0x00, 0x00, 0x00, 0x00, 0x00, 0x00, 0x00, 0x00, 0x00, 0x00, 0x00, 0x00, 0x00, 0x00,
  0x00, 0x00, 0x00, 0x00, 0x00, 0x00, 0x00, 0x00, 0x00, 0x00, 0x00, 0x00, 0x00, 0x00, 0x00, 0x00,
  0x00, 0x00, 0x00, 0x00, 0x00, 0x00, 0x00, 0x00, 0x00, 0x00, 0x00, 0x00, 0x00, 0x00, 0x00, 0x00,
  0x00, 0x00, 0x00, 0x00, 0x00, 0x00, 0x00, 0x00, 0x00, 0x00, 0x00, 0x00, 0x00, 0x00, 0x00, 0x00,
  0x00, 0x00, 0x00, 0x00, 0x00, 0x00, 0x00, 0x00, 0x00, 0x00, 0x00, 0x00, 0x00, 0x00, 0x00, 0x00,
  0x00, 0x00, 0x00, 0x00, 0x00, 0x00, 0x00, 0x00, 0x00, 0x00, 0x00, 0x00, 0x00, 0x00, 0x00, 0x00,
  0x00, 0x00, 0x00, 0x00, 0x00, 0x00, 0x00, 0x00, 0x00, 0x00, 0x00, 0x00, 0x00, 0x00, 0x00, 0x00,
  0x00, 0x00, 0x00, 0x00, 0x00, 0x00, 0x00, 0x00, 0x00, 0x00, 0x00, 0x00, 0x00, 0x00, 0x00, 0x00,
  0x00, 0x00, 0x00, 0x00, 0x00, 0x00, 0x00, 0x00, 0x00, 0x00, 0x00, 0x00, 0x00, 0x00, 0x00, 0x00,
  0x00, 0x00, 0x00, 0x00, 0x00, 0x00, 0x00, 0x00, 0x00, 0x00, 0x00, 0x00, 0x00, 0x00, 0x00, 0x00,
  0x00, 0x00, 0x00, 0x00, 0x00, 0x00, 0x00, 0x00, 0x00, 0x00, 0x00, 0x00, 0x00, 0x00, 0x00, 0x00,
  0x00, 0x00, 0x00, 0x00, 0x00, 0x00, 0x00, 0x00, 0x00, 0x00, 0x00, 0x00, 0x00, 0x00, 0x00, 0x00,
  0x00, 0x00, 0x00, 0x00, 0x00, 0x00, 0x00, 0x00, 0x00, 0x00, 0x00, 0x00, 0x00, 0x00, 0x00, 0x00,
  0x00, 0x00, 0x00, 0x00, 0x00, 0x00, 0x00, 0x00, 0x00, 0x00, 0x00, 0x00, 0x00, 0x00, 0x00, 0x00,
  0x00, 0x00, 0x00, 0x00, 0x00, 0x00, 0x00, 0x00, 0x00, 0x00, 0x00, 0x00, 0x00, 0x00, 0x00, 0x00,
  0x00, 0x00, 0x00, 0x00, 0x00, 0x00, 0x00, 0x00, 0x00, 0x00, 0x00, 0x00, 0x00, 0x00, 0x00, 0x00,
  0x00, 0x00, 0x00, 0x00, 0x00, 0x00, 0x00, 0x00, 0x00, 0x00, 0x00, 0x00, 0x00, 0x00, 0x00, 0x00,
  0x00, 0x00, 0x00, 0x00, 0x00, 0x00, 0x00, 0x00, 0x00, 0x00, 0x00, 0x00, 0x00, 0x00, 0x00, 0x00,
  0x00, 0x00, 0x00, 0x00, 0x00, 0x00, 0x00, 0x00, 0x00, 0x00, 0x00, 0x00, 0x00, 0x00, 0x00, 0x00,
  0x00, 0x00, 0x00, 0x00, 0x00, 0x00, 0x00, 0x00, 0x00, 0x00, 0x00, 0x00, 0x00, 0x00, 0x00, 0x00,
  0x00, 0x00, 0x00, 0x00, 0x00, 0x00, 0x00, 0x00, 0x00, 0x00, 0x00, 0x00, 0x00, 0x00, 0x00, 0x00,
  0x00, 0x00, 0x00, 0x00, 0x00, 0x00, 0x00, 0x00, 0x00, 0x00, 0x00, 0x00, 0x00, 0x00, 0x00, 0x00,
  0x00, 0x00, 0x00, 0x00, 0x00, 0x00, 0x00, 0x00, 0x00, 0x00, 0x00, 0x00, 0x00, 0x00, 0x00, 0x00,
  0x00, 0x00, 0x00, 0x00, 0x00, 0x00, 0x00, 0x00, 0x00, 0x00, 0x00, 0x00, 0x00, 0x00, 0x00, 0x00,
  0x00, 0x00, 0x00, 0x00, 0x00, 0x00, 0x00, 0x00, 0x00, 0x00, 0x3f, 0xc7, 0xfe, 0x3e, 0x04, 0x7f,
  0xff, 0xff, 0xff, 0xff, 0xff, 0xff, 0xff, 0xff, 0xff, 0xff, 0xff, 0xff, 0xff, 0xff, 0xff, 0xff,
  0xff, 0xff, 0xff, 0xe0, 0x00, 0x00, 0x00, 0x00, 0x3f, 0xc7, 0xfe, 0x3e, 0x06, 0x3f, 0xff, 0xff,
  0xff, 0xff, 0xff, 0xff, 0xff, 0xff, 0xff, 0xff, 0xff, 0xff, 0xff, 0xff, 0xff, 0xff, 0xff, 0xff,
  0xff, 0xe0, 0x00, 0x00, 0x00, 0x00, 0x3f, 0xc0, 0x00, 0x00, 0x07, 0x00, 0x00, 0x00, 0x00, 0x00,
  0x00, 0x00, 0x00, 0x00, 0x00, 0x00, 0x00, 0x00, 0x00, 0x00, 0x00, 0x00, 0x00, 0x00, 0x00, 0x00,
  0x00, 0x00, 0x00, 0x00, 0x3f, 0xc0, 0x00, 0x00, 0x07, 0x80, 0x00, 0x00, 0x00, 0x00, 0x00, 0x00,
  0x00, 0x00, 0x00, 0x00, 0x00, 0x00, 0x00, 0x00, 0x00, 0x00, 0x00, 0x00, 0x00, 0x00, 0x00, 0x00,
  0x00, 0x00, 0x3f, 0xc0, 0x00, 0x00, 0x07, 0xc0, 0x00, 0x00, 0x00, 0x00, 0x00, 0x00, 0x00, 0x00,
  0x00, 0x00, 0x00, 0x00, 0x00, 0x00, 0x00, 0x00, 0x00, 0x00, 0x00, 0x00, 0x00, 0x00, 0x00, 0x00,
  0x3f, 0xc1, 0xfe, 0x3f, 0xc7, 0xe0, 0xfe, 0x3f, 0xff, 0xf8, 0x0f, 0xff, 0xfe, 0x7f, 0xff, 0xf0,
  0x00, 0x03, 0xfc, 0x00, 0xff, 0xff, 0xe3, 0xff, 0xff, 0x80, 0x00, 0x00, 0x00, 0x00, 0x3f, 0xc1,
  0xfe, 0x3f, 0xc7, 0xf0, 0xfe, 0x3f, 0xff, 0xfc, 0x1f, 0xff, 0xfe, 0x7f, 0xff, 0xf8, 0x00, 0x03,
  0xfc, 0x01, 0xff, 0xff, 0xe3, 0xff, 0xff, 0xc0, 0x00, 0x00, 0x00, 0x00, 0x3f, 0xc9, 0xfe, 0x3f,
  0xc7, 0xf8, 0xfe, 0x3f, 0xc7, 0xfe, 0x3f, 0xf0, 0x00, 0x00, 0x0f, 0xfc, 0x00, 0x03, 0xfc, 0x03,
  0xff, 0x1f, 0xe3, 0xfc, 0x7f, 0xe0, 0x00, 0x00, 0x00, 0x00, 0x3f, 0xdd, 0xfe, 0x3f, 0xc7, 0xfc,
  0xfe, 0x3f, 0xc1, 0xfe, 0x3f, 0xc0, 0x00, 0x00, 0x03, 0xfc, 0x00, 0x03, 0xfc, 0x03, 0xfc, 0x1f,
  0xe3, 0xfc, 0x1f, 0xe0, 0x00, 0x00, 0x00, 0x00, 0x3f, 0xff, 0xfe, 0x3f, 0xc7, 0xfe, 0xfe, 0x3f,
  0xc1, 0xfe, 0x3f, 0xc0, 0x00, 0x00, 0x03, 0xfc, 0x00, 0x03, 0xfc, 0x03, 0xfc, 0x1f, 0xe3, 0xfc,
  0x1f, 0xe0, 0x00, 0x00, 0x00, 0x00, 0x3f, 0xff, 0xfe, 0x3f, 0xc7, 0xff, 0xfe, 0x3f, 0xc1, 0xfe,
  0x3f, 0xfe, 0x00, 0x7f, 0xff, 0xfc, 0x00, 0x03, 0xfc, 0x03, 0xff, 0xff, 0xe3, 0xff, 0xff, 0xe0,
  0x00, 0x00, 0x00, 0x00, 0x3f, 0xff, 0xfe, 0x3f, 0xc7, 0xff, 0xfe, 0x3f, 0xc1, 0xfe, 0x3f, 0xfe,
  0x00, 0x7f, 0xff, 0xf8, 0x00, 0x03, 0xfc, 0x03, 0xff, 0xff, 0xe3, 0xff, 0xff, 0xe0, 0x00, 0x00,
  0x00, 0x00, 0x3f, 0xff, 0xfe, 0x3f, 0xc7, 0xff, 0xfe, 0x3f, 0xc1, 0xfe, 0x3f, 0xfe, 0x00, 0x7f,
  0xff, 0x80, 0x00, 0x03, 0xfc, 0x03, 0xff, 0xff, 0xe3, 0xff, 0xff, 0xe0, 0x00, 0x00, 0x00, 0x00,
  0x3f, 0xf7, 0xfe, 0x3f, 0xc7, 0xfb, 0xfe, 0x3f, 0xc1, 0xfe, 0x3f, 0xc0, 0x00, 0x7f, 0x83, 0xf0,
  0x00, 0x03, 0xfc, 0x03, 0xfc, 0x1f, 0xe3, 0xfc, 0x1f, 0xe0, 0x00, 0x00, 0x00, 0x00, 0x3f, 0xe3,
  0xfe, 0x3f, 0xc7, 0xf9, 0xfe, 0x3f, 0xc1, 0xfe, 0x3f, 0xc0, 0x00, 0x7f, 0x83, 0xf8, 0x00, 0x03,
  0xfc, 0x03, 0xfc, 0x1f, 0xe3, 0xfc, 0x1f, 0xe0, 0x00, 0x00, 0x00, 0x00, 0x3f, 0xc1, 0xfe, 0x3f,
  0xc7, 0xf8, 0xfe, 0x3f, 0xff, 0xfe, 0x3f, 0xff, 0xfe, 0x7f, 0x83, 0xfc, 0x00, 0x03, 0xff, 0xe3,
  0xfc, 0x1f, 0xe3, 0xff, 0xff, 0xe0, 0x00, 0x00, 0x00, 0x00, 0x3f, 0x80, 0xfe, 0x3f, 0xc7, 0xf8,
  0x7e, 0x3f, 0xff, 0xfe, 0x3f, 0xff, 0xfe, 0x7f, 0x83, 0xfc, 0x00, 0x03, 0xff, 0xe3, 0xfc, 0x1f,
  0xe3, 0xff, 0xff, 0xe0, 0x00, 0x00, 0x00, 0x00, 0x3f, 0x00, 0x7e, 0x3f, 0xc7, 0xf8, 0x3e, 0x3f,
  0xff, 0xfe, 0x3f, 0xff, 0xfe, 0x7f, 0x83, 0xfc, 0x00, 0x03, 0xff, 0xe3, 0xfc, 0x1f, 0xe3, 0xff,
  0xff, 0xe0, 0x00, 0x00, 0x00, 0x00, 0x00, 0x00, 0x3e, 0x00, 0x00, 0x00, 0x1e, 0x00, 0x00, 0x00,
  0x00, 0x00, 0x00, 0x00, 0x00, 0x00, 0x00, 0x00, 0x00, 0x00, 0x00, 0x1f, 0xe0, 0x00, 0x00, 0x00,
  0x00, 0x00, 0x00, 0x00, 0x00, 0x00, 0x1e, 0x00, 0x00, 0x00, 0x0e, 0x00, 0x00, 0x00, 0x00, 0x00,
  0x00, 0x00, 0x00, 0x00, 0x00, 0x00, 0x00, 0x00, 0x00, 0x1f, 0xe0, 0x00, 0x00, 0x00, 0x00, 0x00,
  0x00, 0x00, 0x00, 0x00, 0x0e, 0x00, 0x00, 0x00, 0x06, 0x00, 0x00, 0x00, 0x00, 0x00, 0x00, 0x00,
  0x00, 0x00, 0x00, 0x00, 0x00, 0x00, 0x00, 0x1f, 0xe0, 0x00, 0x00, 0x00, 0x00, 0x00, 0x00, 0x00,
  0x3f, 0xff, 0xe6, 0x3f, 0xff, 0xff, 0xe2, 0x3f, 0xff, 0xff, 0xff, 0xff, 0xff, 0xff, 0xff, 0xff,
  0xff, 0xff, 0xff, 0xff, 0xff, 0x9f, 0xe3, 0xff, 0xff, 0xfc, 0x00, 0x00, 0x00, 0x00, 0x3f, 0xff,
  0xf2, 0x3f, 0xff, 0xff, 0xf0, 0x3f, 0xff, 0xff, 0xff, 0xff, 0xff, 0xff, 0xff, 0xff, 0xff, 0xff,
  0xff, 0xff, 0xff, 0x9f, 0xe3, 0xff, 0xff, 0xfc, 0x00, 0x00, 0x00, 0x00, 0x00, 0x00, 0x00, 0x00,
  0x00, 0x00, 0x00, 0x00, 0x00, 0x00, 0x00, 0x00, 0x00, 0x00, 0x00, 0x00, 0x00, 0x00, 0x00, 0x00,
  0x00, 0x00, 0x00, 0x00, 0x00, 0x00, 0x00, 0x00, 0x00, 0x00, 0x00, 0x00, 0x00, 0x00, 0x00, 0x00,
  0x00, 0x00, 0x00, 0x00, 0x00, 0x00, 0x00, 0x00, 0x00, 0x00, 0x00, 0x00, 0x00, 0x00, 0x00, 0x00,
  0x00, 0x00, 0x00, 0x00, 0x00, 0x00, 0x00, 0x00, 0x00, 0x00, 0x00, 0x00, 0x00, 0x00, 0x00, 0x00,
  0x00, 0x00, 0x00, 0x00, 0x00, 0x00, 0x00, 0x00, 0x00, 0x00, 0x00, 0x00, 0x00, 0x00, 0x00, 0x00,
  0x00, 0x00, 0x00, 0x00, 0x00, 0x00, 0x00, 0x00, 0x00, 0x00, 0x00, 0x00, 0x00, 0x00, 0x00, 0x00,
  0x00, 0x00, 0x00, 0x00, 0x00, 0x00, 0x00, 0x00, 0x00, 0x00, 0x00, 0x00, 0x00, 0x00, 0x00, 0x00,
  0x00, 0x00, 0x00, 0x00, 0x00, 0x00, 0x00, 0x00, 0x00, 0x00, 0x00, 0x00, 0x00, 0x00, 0x00, 0x00,
  0x00, 0x00, 0x00, 0x00, 0x00, 0x00, 0x00, 0x00, 0x00, 0x00, 0x00, 0x00, 0x00, 0x00, 0x00, 0x00,
  0x00, 0x00, 0x00, 0x00, 0x00, 0x00, 0x00, 0x00, 0x00, 0x00, 0x00, 0x00, 0x00, 0x00, 0x00, 0x00,
  0x00, 0x00, 0x00, 0x00, 0x00, 0x00, 0x00, 0x00, 0x00, 0x00, 0x00, 0x00, 0x00, 0x00, 0x00, 0x00,
  0x00, 0x00, 0x00, 0x00, 0x00, 0x00, 0x00, 0x00, 0x00, 0x00, 0x00, 0x00, 0x00, 0x00, 0x00, 0x00,
  0x00, 0x00, 0x00, 0x00, 0x00, 0x00, 0x00, 0x00, 0x00, 0x00, 0x00, 0x00, 0x00, 0x00, 0x00, 0x00,
  0x00, 0x00, 0x00, 0x00, 0x00, 0x00, 0x00, 0x00, 0x00, 0x00, 0x00, 0x00, 0x00, 0x00, 0x00, 0x00,
  0x00, 0x00, 0x00, 0x00, 0x00, 0x00, 0x00, 0x00, 0x00, 0x00, 0x00, 0x00, 0x00, 0x00, 0x00, 0x00,
  0x00, 0x00, 0x00, 0x00, 0x00, 0x00, 0x00, 0x00, 0x00, 0x00, 0x00, 0x00, 0x00, 0x00, 0x00, 0x00,
  0x00, 0x00, 0x00, 0x00, 0x00, 0x00, 0x00, 0x00, 0x00, 0x00, 0x00, 0x00, 0x00, 0x00, 0x00, 0x00,
  0x00, 0x00, 0x00, 0x00, 0x00, 0x00, 0x00, 0x00, 0x00, 0x00, 0x00, 0x00, 0x00, 0x00, 0x00, 0x00,
  0x00, 0x00, 0x00, 0x00, 0x00, 0x00, 0x00, 0x00, 0x00, 0x00, 0x00, 0x00, 0x00, 0x00, 0x00, 0x00,
  0x00, 0x00, 0x00, 0x00, 0x00, 0x00, 0x00, 0x00, 0x00, 0x00, 0x00, 0x00, 0x00, 0x00, 0x00, 0x00,
  0x00, 0x00, 0x00, 0x00, 0x00, 0x00, 0x00, 0x00, 0x00, 0x00, 0x00, 0x00, 0x00, 0x00, 0x00, 0x00,
  0x00, 0x00, 0x00, 0x00, 0x00, 0x00, 0x00, 0x00, 0x00, 0x00, 0x00, 0x00, 0x00, 0x00, 0x00, 0x00,
  0x00, 0x00, 0x00, 0x00, 0x00, 0x00, 0x00, 0x00, 0x00, 0x00, 0x00, 0x00, 0x00, 0x00, 0x00, 0x00,
  0x00, 0x00, 0x00, 0x00, 0x00, 0x00, 0x00, 0x00, 0x00, 0x00, 0x00, 0x00, 0x00, 0x00, 0x00, 0x00,
  0x00, 0x00, 0x00, 0x00, 0x00, 0x00, 0x00, 0x00, 0x00, 0x00, 0x00, 0x00, 0x00, 0x00, 0x00, 0x00,
  0x00, 0x00, 0x00, 0x00, 0x00, 0x00, 0x00, 0x00, 0x00, 0x00, 0x00, 0x00, 0x00, 0x00, 0x00, 0x00,
  0x00, 0x00, 0x00, 0x00, 0x00, 0x00, 0x00, 0x00, 0x00, 0x00, 0x00, 0x00, 0x00, 0x00, 0x00, 0x00,
  0x00, 0x00, 0x00, 0x00, 0x00, 0x00, 0x00, 0x00, 0x00, 0x00, 0x00, 0x00, 0x00, 0x00, 0x00, 0x00,
  0x00, 0x00, 0x00, 0x00, 0x00, 0x00, 0x00, 0x00, 0x00, 0x00, 0x00, 0x00, 0x00, 0x00, 0x00, 0x00,
  0x00, 0x00, 0x00, 0x00, 0x00, 0x00, 0x00, 0x00, 0x00, 0x00, 0x00, 0x00, 0x00, 0x00, 0x00, 0x00,
  0x00, 0x00, 0x00, 0x00, 0x00, 0x00, 0x00, 0x00, 0x00, 0x00, 0x00, 0x00, 0x00, 0x00, 0x00, 0x00,
  0x00, 0x00, 0x00, 0x00, 0x00, 0x00, 0x00, 0x00, 0x00, 0x00, 0x00, 0x00, 0x00, 0x00, 0x00, 0x00,
  0x00, 0x00, 0x00, 0x00, 0x00, 0x00, 0x00, 0x00, 0x00, 0x00, 0x00, 0x00, 0x00, 0x00, 0x00, 0x00,
  0x00, 0x00, 0x00, 0x00, 0x00, 0x00
};
// 'main_logo', 240x60px
const unsigned char  main_logo [] PROGMEM = {
  0x00, 0x00, 0x00, 0x00, 0x00, 0x00, 0x00, 0x00, 0x00, 0x00, 0x00, 0x00, 0x00, 0x00, 0x00, 0x00,
  0x00, 0x00, 0x00, 0x00, 0x00, 0x00, 0x00, 0x00, 0x00, 0x00, 0x00, 0x00, 0x00, 0x00, 0x00, 0x00,
  0x00, 0x00, 0x00, 0x00, 0x00, 0x00, 0x00, 0x00, 0x00, 0x00, 0x00, 0x00, 0x00, 0x00, 0x00, 0x00,
  0x00, 0x00, 0x00, 0x00, 0x00, 0x00, 0x00, 0x00, 0x00, 0x00, 0x00, 0x00, 0x00, 0x00, 0x00, 0x00,
  0x00, 0x00, 0x00, 0x00, 0x00, 0x00, 0x00, 0x00, 0x00, 0x00, 0x00, 0x00, 0x00, 0x00, 0x00, 0x00,
  0x00, 0x00, 0x00, 0x00, 0x00, 0x00, 0x00, 0x00, 0x00, 0x00, 0x00, 0x00, 0x00, 0x00, 0x00, 0x00,
  0x00, 0x00, 0x00, 0x00, 0x00, 0x00, 0x00, 0x00, 0x00, 0x00, 0x00, 0x00, 0x00, 0x00, 0x00, 0x00,
  0x00, 0x00, 0x00, 0x00, 0x00, 0x00, 0x00, 0x00, 0x00, 0x00, 0x00, 0x00, 0x00, 0x00, 0x00, 0x00,
  0x00, 0x00, 0x00, 0x00, 0x00, 0x00, 0x00, 0x00, 0x00, 0x00, 0x00, 0x00, 0x00, 0x00, 0x00, 0x00,
  0x00, 0x00, 0x00, 0x00, 0x00, 0x00, 0x00, 0x00, 0x00, 0x00, 0x00, 0x00, 0x00, 0x00, 0x00, 0x00,
  0x00, 0x00, 0x00, 0x00, 0x00, 0x00, 0x00, 0x00, 0x00, 0x00, 0x00, 0x00, 0x00, 0x00, 0x00, 0x00,
  0x00, 0x00, 0x00, 0x00, 0x00, 0x00, 0x00, 0x00, 0x00, 0x00, 0x00, 0x00, 0x00, 0x00, 0x00, 0x00,
  0x00, 0x00, 0x00, 0x00, 0x00, 0x00, 0x00, 0x00, 0x00, 0x00, 0x00, 0x00, 0x00, 0x00, 0x00, 0x00,
  0x00, 0x00, 0x00, 0x00, 0x00, 0x00, 0x00, 0x00, 0x00, 0x00, 0x00, 0x00, 0x00, 0x00, 0x00, 0x00,
  0x00, 0x00, 0x00, 0x00, 0x00, 0x00, 0x00, 0x00, 0x00, 0x00, 0x00, 0x00, 0x00, 0x00, 0x00, 0x00,
  0x00, 0x00, 0x00, 0x00, 0x00, 0x00, 0x00, 0x7c, 0x00, 0x07, 0xc0, 0x03, 0xff, 0x80, 0x00, 0x00,
  0x1f, 0x00, 0x03, 0xe0, 0x7f, 0xff, 0xc0, 0x00, 0x00, 0x00, 0x00, 0x00, 0x00, 0x00, 0x00, 0x00,
  0x00, 0x00, 0x00, 0x00, 0x00, 0x7c, 0x00, 0x07, 0xc0, 0x1f, 0xff, 0xf0, 0x00, 0x00, 0x1f, 0x00,
  0x03, 0xe0, 0x7f, 0xff, 0xf8, 0x00, 0x00, 0x00, 0x00, 0x00, 0x00, 0x00, 0x00, 0x00, 0x00, 0x00,
  0x00, 0x00, 0x00, 0x7c, 0x00, 0x07, 0xc0, 0x3f, 0xff, 0xf8, 0x00, 0x00, 0x1f, 0x00, 0x03, 0xe0,
  0x7f, 0xff, 0xfc, 0x00, 0x00, 0x00, 0x00, 0x00, 0x00, 0x00, 0x00, 0x00, 0x00, 0x00, 0x00, 0x00,
  0x00, 0x7c, 0x00, 0x07, 0xc0, 0x7f, 0xff, 0xfc, 0x00, 0x00, 0x1f, 0x00, 0x03, 0xe0, 0x7f, 0xff,
  0xfe, 0x00, 0x00, 0x00, 0x00, 0x00, 0x00, 0x00, 0x00, 0x00, 0x00, 0x00, 0x00, 0x00, 0x00, 0x7c,
  0x00, 0x07, 0xc0, 0x7f, 0xff, 0xfc, 0x00, 0x00, 0x1f, 0x00, 0x03, 0xe0, 0x7f, 0xff, 0xfe, 0x00,
  0x00, 0x00, 0x00, 0x00, 0x00, 0x00, 0x00, 0x00, 0x00, 0x00, 0x00, 0x00, 0x00, 0x7c, 0x00, 0x07,
  0xc0, 0xfc, 0x00, 0x7e, 0x00, 0x00, 0x1f, 0x00, 0x03, 0xe0, 0x7c, 0x00, 0x3f, 0x00, 0x00, 0x00,
  0x00, 0x00, 0x00, 0x00, 0x00, 0x00, 0x00, 0x00, 0x00, 0x00, 0x00, 0x7c, 0x00, 0x07, 0xc0, 0xf8,
  0x00, 0x3e, 0x00, 0x00, 0x1f, 0x00, 0x03, 0xe0, 0x7c, 0x00, 0x1f, 0x00, 0x00, 0x00, 0x00, 0x00,
  0x00, 0x00, 0x00, 0x00, 0x00, 0x00, 0x00, 0x00, 0x00, 0x7c, 0x00, 0x07, 0xc0, 0xf8, 0x00, 0x3e,
  0x00, 0x00, 0x1f, 0x00, 0x03, 0xe0, 0x7c, 0x00, 0x1f, 0x00, 0x00, 0x00, 0x00, 0x00, 0x00, 0x00,
  0x00, 0x00, 0x00, 0x00, 0x00, 0x00, 0x00, 0x7c, 0x00, 0x07, 0xc0, 0xf8, 0x00, 0x3e, 0x00, 0x00,
  0x1f, 0x00, 0x03, 0xe0, 0x7c, 0x00, 0x1f, 0x00, 0x00, 0x00, 0x00, 0x00, 0x00, 0x00, 0x00, 0x00,
  0x00, 0x00, 0x00, 0x00, 0x00, 0x7c, 0x00, 0x07, 0xc0, 0xf8, 0x00, 0x3e, 0x00, 0x00, 0x1f, 0x00,
  0x03, 0xe0, 0x7c, 0x00, 0x1f, 0x00, 0x00, 0x00, 0x00, 0x00, 0x00, 0x00, 0x00, 0x00, 0x00, 0x00,
  0x00, 0x00, 0x00, 0x7c, 0x00, 0x07, 0xc0, 0xf8, 0x00, 0x3e, 0x00, 0x00, 0x1f, 0x00, 0x03, 0xe0,
  0x7c, 0x00, 0x1f, 0x00, 0x00, 0x00, 0x00, 0x00, 0x00, 0x00, 0x00, 0x00, 0x00, 0x00, 0x00, 0x00,
  0x00, 0x7c, 0x00, 0x07, 0xc0, 0xf8, 0x00, 0x3e, 0x00, 0x00, 0x1f, 0x00, 0x03, 0xe0, 0x7c, 0x00,
  0x1f, 0x00, 0x00, 0x00, 0x00, 0x00, 0x00, 0x00, 0x00, 0x00, 0x00, 0x00, 0x00, 0x00, 0x00, 0x7c,
  0x00, 0x07, 0xc0, 0xf8, 0x00, 0x3e, 0x00, 0x00, 0x1f, 0x00, 0x03, 0xe0, 0x7c, 0x00, 0x1f, 0x00,
  0x00, 0x00, 0x00, 0x00, 0x00, 0x00, 0x00, 0x00, 0x00, 0x00, 0x00, 0x00, 0x00, 0x7c, 0x00, 0x07,
  0xc0, 0xf8, 0x00, 0x3e, 0x00, 0x00, 0x1f, 0x00, 0x03, 0xe0, 0x7c, 0x00, 0x1f, 0x00, 0x00, 0x00,
  0x00, 0x00, 0x00, 0x00, 0x00, 0x00, 0x00, 0x00, 0x00, 0x00, 0x00, 0x7c, 0x00, 0x07, 0xc0, 0xf8,
  0x00, 0x3e, 0x00, 0x00, 0x1f, 0x00, 0x03, 0xe0, 0x7c, 0x00, 0x1f, 0x00, 0x00, 0x00, 0x00, 0x00,
  0x00, 0x00, 0x00, 0x00, 0x00, 0x00, 0x00, 0x00, 0x00, 0x7c, 0x00, 0x07, 0xc0, 0xf8, 0x00, 0x3e,
  0x00, 0x00, 0x1f, 0x00, 0x03, 0xe0, 0x7c, 0x00, 0x1f, 0x00, 0x00, 0x00, 0x00, 0x00, 0x00, 0x00,
  0x00, 0x00, 0x00, 0x00, 0x00, 0x00, 0x00, 0x7c, 0x00, 0x07, 0xc0, 0xf8, 0x00, 0x3e, 0x00, 0x00,
  0x1f, 0x00, 0x03, 0xe0, 0x7c, 0x00, 0x1f, 0x00, 0x00, 0x00, 0x00, 0x00, 0x00, 0x00, 0x00, 0x00,
  0x00, 0x00, 0x00, 0x00, 0x00, 0x7c, 0x00, 0x07, 0xc0, 0xf8, 0x00, 0x3e, 0x00, 0x00, 0x1f, 0x00,
  0x03, 0xe0, 0x7c, 0x00, 0x1f, 0x00, 0x00, 0x00, 0x00, 0x00, 0x00, 0x00, 0x00, 0x00, 0x00, 0x00,
  0x00, 0x00, 0x00, 0x7c, 0x00, 0x07, 0xc0, 0xf8, 0x00, 0x3e, 0x00, 0x00, 0x1f, 0x00, 0x03, 0xe0,
  0x7c, 0x00, 0x1f, 0x00, 0x00, 0x00, 0x00, 0x00, 0x00, 0x00, 0x00, 0x00, 0x00, 0x00, 0x00, 0x00,
  0x00, 0x7c, 0x00, 0x07, 0xc0, 0xf8, 0x00, 0x3e, 0x00, 0x00, 0x1f, 0x00, 0x03, 0xe0, 0x7c, 0x00,
  0x1f, 0x00, 0x00, 0x00, 0x00, 0x00, 0x00, 0x00, 0x00, 0x00, 0x00, 0x00, 0x00, 0x00, 0x00, 0x7c,
  0x00, 0x07, 0xc0, 0xf8, 0x00, 0x3e, 0x00, 0x00, 0x1f, 0x00, 0x03, 0xe0, 0x7c, 0x00, 0x1f, 0x00,
  0x00, 0x00, 0x00, 0x00, 0x00, 0x00, 0x00, 0x00, 0x00, 0x00, 0x00, 0x00, 0x00, 0x7c, 0x00, 0x07,
  0xc0, 0xf8, 0x00, 0x3e, 0x00, 0x00, 0x1f, 0x00, 0x03, 0xe0, 0x7c, 0x00, 0x1f, 0x00, 0x00, 0x00,
  0x00, 0x00, 0x00, 0x00, 0x00, 0x00, 0x00, 0x00, 0x00, 0x00, 0x00, 0x7c, 0x00, 0x07, 0xc0, 0xf8,
  0x00, 0x3e, 0x00, 0x00, 0x1f, 0x00, 0x03, 0xe0, 0x7c, 0x00, 0x1f, 0x00, 0x00, 0x00, 0x00, 0x00,
  0x00, 0x00, 0x00, 0x00, 0x00, 0x00, 0x00, 0x00, 0x00, 0x7c, 0x00, 0x07, 0xc0, 0xf8, 0x00, 0x3e,
  0x00, 0x00, 0xff, 0xff, 0xff, 0xe0, 0x7c, 0x00, 0x1f, 0x00, 0x00, 0x00, 0x00, 0x00, 0x00, 0x00,
  0x00, 0x00, 0x00, 0x00, 0x00, 0x00, 0x00, 0x7c, 0x00, 0x07, 0xc0, 0xf8, 0x00, 0x3e, 0x00, 0x00,
  0xff, 0xff, 0xff, 0xe0, 0x7c, 0x00, 0x1f, 0x00, 0x00, 0x00, 0x00, 0x00, 0x00, 0x00, 0x00, 0x00,
  0x00, 0x00, 0x00, 0x00, 0x00, 0x7c, 0x00, 0x07, 0xc0, 0xf8, 0x00, 0x3e, 0x00, 0x00, 0xff, 0xff,
  0xff, 0xe0, 0x7c, 0x00, 0x1f, 0x00, 0x00, 0x00, 0x00, 0x00, 0x00, 0x00, 0x00, 0x00, 0x00, 0x00,
  0x00, 0x00, 0x00, 0x7c, 0x00, 0x07, 0xc0, 0xf8, 0x00, 0x3e, 0x00, 0x00, 0xff, 0xff, 0xff, 0xe0,
  0x7c, 0x00, 0x1f, 0x00, 0x00, 0x00, 0x00, 0x00, 0x00, 0x00, 0x00, 0x00, 0x00, 0x00, 0x00, 0x00,
  0x00, 0x7c, 0x00, 0x07, 0xc0, 0xf8, 0x00, 0x3e, 0x00, 0x00, 0xff, 0xff, 0xff, 0xe0, 0x7c, 0x00,
  0x1f, 0x00, 0x00, 0x00, 0x00, 0x00, 0x00, 0x00, 0x00, 0x00, 0x00, 0x00, 0x00, 0x00, 0x00, 0x7c,
  0x00, 0x07, 0xc0, 0xf8, 0x00, 0x3e, 0x00, 0x00, 0x1f, 0x00, 0x03, 0xe0, 0x7c, 0x00, 0x1f, 0x00,
  0x00, 0x00, 0x00, 0x00, 0x00, 0x00, 0x00, 0x00, 0x00, 0x00, 0x00, 0x00, 0x00, 0x7c, 0x00, 0x07,
  0xc0, 0xf8, 0x00, 0x3e, 0x00, 0x00, 0x1f, 0x00, 0x03, 0xe0, 0x7c, 0x00, 0x1f, 0x00, 0x00, 0x00,
  0x00, 0x00, 0x00, 0x00, 0x00, 0x00, 0x00, 0x00, 0x00, 0x00, 0x00, 0x7c, 0x00, 0x07, 0xc0, 0xf8,
  0x00, 0x3e, 0x00, 0x00, 0x1f, 0x00, 0x03, 0xe0, 0x7c, 0x00, 0x1f, 0x00, 0x00, 0x00, 0x00, 0x00,
  0x00, 0x00, 0x00, 0x00, 0x00, 0x00, 0x00, 0x00, 0x00, 0x7c, 0x00, 0x07, 0xc0, 0xf8, 0x00, 0x3e,
  0x00, 0x00, 0x1f, 0x00, 0x03, 0xe0, 0x7c, 0x00, 0x1f, 0x00, 0x00, 0x00, 0x00, 0x00, 0x00, 0x00,
  0x00, 0x00, 0x00, 0x00, 0x00, 0x00, 0x00, 0x7c, 0x00, 0x07, 0xc0, 0xf8, 0x00, 0x3e, 0x00, 0x00,
  0x1f, 0x00, 0x03, 0xe0, 0x7c, 0x00, 0x1f, 0x00, 0x00, 0x00, 0x00, 0x00, 0x00, 0x00, 0x00, 0x00,
  0x00, 0x00, 0x00, 0x00, 0x00, 0x7c, 0x00, 0x07, 0xc0, 0xfc, 0x00, 0x7e, 0x00, 0x00, 0x1f, 0x00,
  0x03, 0xe0, 0x7c, 0x00, 0x3f, 0x00, 0x00, 0x00, 0x00, 0x00, 0x00, 0x00, 0x00, 0x00, 0x00, 0x00,
  0x00, 0x00, 0x00, 0x7f, 0xff, 0x07, 0xc0, 0x7f, 0xff, 0xfc, 0x00, 0x00, 0x1f, 0x00, 0x03, 0xe0,
  0x7f, 0xff, 0xfe, 0x00, 0x00, 0x00, 0x00, 0x00, 0x00, 0x00, 0x00, 0x00, 0x00, 0x00, 0x00, 0x00,
  0x00, 0x7f, 0xff, 0x07, 0xc0, 0x7f, 0xff, 0xfc, 0x00, 0x00, 0x1f, 0x00, 0x03, 0xe0, 0x7f, 0xff,
  0xfe, 0x00, 0x00, 0x00, 0x00, 0x00, 0x00, 0x00, 0x00, 0x00, 0x00, 0x00, 0x00, 0x00, 0x00, 0x7f,
  0xff, 0x07, 0xc0, 0x3f, 0xff, 0xf8, 0x00, 0x00, 0x1f, 0x00, 0x03, 0xe0, 0x7f, 0xff, 0xfc, 0x00,
  0x00, 0x00, 0x00, 0x00, 0x00, 0x00, 0x00, 0x00, 0x00, 0x00, 0x00, 0x00, 0x00, 0x7f, 0xff, 0x07,
  0xc0, 0x1f, 0xff, 0xf0, 0x00, 0x00, 0x1f, 0x00, 0x03, 0xe0, 0x7f, 0xff, 0xf8, 0x00, 0x00, 0x00,
  0x00, 0x00, 0x00, 0x00, 0x00, 0x00, 0x00, 0x00, 0x00, 0x00, 0x00, 0x7f, 0xff, 0x07, 0xc0, 0x03,
  0xff, 0x80, 0x00, 0x00, 0x1f, 0x00, 0x03, 0xe0, 0x7f, 0xff, 0xc0, 0x00, 0x00, 0x00, 0x00, 0x00,
  0x00, 0x00, 0x00, 0x00, 0x00, 0x00, 0x00, 0x00, 0x00, 0x00, 0x00, 0x00, 0x00, 0x00, 0x3f, 0x80,
  0x00, 0x00, 0x00, 0x00, 0x00, 0x00, 0x00, 0x00, 0x00, 0x00, 0x00, 0x00, 0x00, 0x00, 0x00, 0x00,
  0x00, 0x00, 0x00, 0x00, 0x00, 0x00, 0x00, 0x00, 0x00, 0x00, 0x00, 0x00, 0x1f, 0xc0, 0x00, 0x00,
  0x00, 0x00, 0x00, 0x00, 0x00, 0x00, 0x00, 0x00, 0x00, 0x00, 0x00, 0x00, 0x00, 0x00, 0x00, 0x00,
  0x00, 0x00, 0x00, 0x00, 0x00, 0x00, 0x00, 0x00, 0x00, 0x00, 0x0f, 0xe0, 0x00, 0x00, 0x00, 0x00,
  0x00, 0x00, 0x00, 0x00, 0x00, 0x00, 0x00, 0x00, 0x00, 0x00, 0x00, 0x00, 0x00, 0x00, 0x00, 0x00,
  0x00, 0x00, 0x00, 0x00, 0x00, 0x00, 0x00, 0x00, 0x07, 0x80, 0x00, 0x00, 0x00, 0x00, 0x00, 0x00,
  0x00, 0x00, 0x00, 0x00, 0x00, 0x00, 0x00, 0x00, 0x00, 0x00, 0x00, 0x00, 0x00, 0x00, 0x00, 0x00,
  0x00, 0x00, 0x00, 0x00, 0x00, 0x00, 0x03, 0x80, 0x00, 0x00, 0x00, 0x00, 0x00, 0x00, 0x00, 0x00,
  0x00, 0x00, 0x00, 0x00, 0x00, 0x00, 0x00, 0x00, 0x00, 0x00, 0x00, 0x00, 0x00, 0x00, 0x00, 0x00,
  0x00, 0x00, 0x00, 0x00, 0x01, 0x00, 0x00, 0x00, 0x00, 0x00, 0x00, 0x00, 0x00, 0x00, 0x00, 0x00,
  0x00, 0x00, 0x00, 0x00, 0x00, 0x00, 0x00, 0x00, 0x00, 0x00, 0x00, 0x00, 0x00, 0x00, 0x00, 0x00,
  0x00, 0x00, 0x00, 0x00, 0x00, 0x00, 0x00, 0x00, 0x00, 0x00, 0x00, 0x00, 0x00, 0x00, 0x00, 0x00,
  0x00, 0x00, 0x00, 0x00, 0x00, 0x00, 0x00, 0x00, 0x00, 0x00, 0x00, 0x00, 0x00, 0x00, 0x00, 0x00,
  0x00, 0x00, 0x00, 0x00, 0x00, 0x00, 0x00, 0x00, 0x00, 0x00, 0x00, 0x00, 0x00, 0x00, 0x00, 0x00,
  0x00, 0x00, 0x00, 0x00, 0x00, 0x00, 0x00, 0x00, 0x00, 0x00, 0x00, 0x00, 0x00, 0x00, 0x00, 0x00,
  0x00, 0x00, 0x00, 0x00, 0x00, 0x00, 0x00, 0x00, 0x00, 0x00, 0x00, 0x00, 0x00, 0x00, 0x00, 0x00,
  0x00, 0x00, 0x00, 0x00, 0x00, 0x00, 0x00, 0x00, 0x00, 0x00, 0x00, 0x00, 0x00, 0x00, 0x00, 0x00,
  0x00, 0x00, 0x00, 0x00, 0x00, 0x00, 0x00, 0x00, 0x00, 0x00, 0x00, 0x00, 0x00, 0x00, 0x00, 0x00,
  0x00, 0x00, 0x00, 0x00, 0x00, 0x00, 0x00, 0x00, 0x00, 0x00, 0x00, 0x00, 0x00, 0x00, 0x00, 0x00,
  0x00, 0x00, 0x00, 0x00, 0x00, 0x00, 0x00, 0x00, 0x00, 0x00, 0x00, 0x00, 0x00, 0x00, 0x00, 0x00,
  0x00, 0x00, 0x00, 0x00, 0x00, 0x00, 0x00, 0x00, 0x00, 0x00, 0x00, 0x00, 0x00, 0x00, 0x00, 0x00,
  0x00, 0x00, 0x00, 0x00, 0x00, 0x00, 0x00, 0x00, 0x00, 0x00, 0x00, 0x00, 0x00, 0x00, 0x00, 0x00,
  0x00, 0x00, 0x00, 0x00, 0x00, 0x00, 0x00, 0x00, 0x00, 0x00, 0x00, 0x00, 0x00, 0x00, 0x00, 0x00,
  0x00, 0x00, 0x00, 0x00, 0x00, 0x00, 0x00, 0x00
};


const unsigned char settings_icon [] PROGMEM = {
  // 'Very-Basic-Settings-Filled-icon-2, 32x32px
  0x00, 0x03, 0xc0, 0x00, 0x00, 0x07, 0xe0, 0x00, 0x00, 0x07, 0xe0, 0x00, 0x02, 0x07, 0xe0, 0x40,
  0x07, 0x07, 0xe0, 0xe0, 0x0f, 0xdf, 0xfb, 0xf0, 0x1f, 0xff, 0xff, 0xf8, 0x0f, 0xff, 0xff, 0xf0,
  0x07, 0xff, 0xff, 0xe0, 0x07, 0xff, 0xff, 0xe0, 0x03, 0xff, 0xff, 0xc0, 0x07, 0xfe, 0x7f, 0xe0,
  0x07, 0xf8, 0x1f, 0xe0, 0x7f, 0xf0, 0x0f, 0xfe, 0xff, 0xf0, 0x0f, 0xff, 0xff, 0xe0, 0x07, 0xff,
  0xff, 0xe0, 0x07, 0xff, 0xff, 0xf0, 0x0f, 0xff, 0x7f, 0xf0, 0x0f, 0xfe, 0x07, 0xf8, 0x1f, 0xe0,
  0x07, 0xfe, 0x7f, 0xe0, 0x03, 0xff, 0xff, 0xc0, 0x07, 0xff, 0xff, 0xe0, 0x07, 0xff, 0xff, 0xe0,
  0x0f, 0xff, 0xff, 0xf0, 0x1f, 0xff, 0xff, 0xf8, 0x0f, 0xdf, 0xfb, 0xf0, 0x07, 0x07, 0xe0, 0xe0,
  0x02, 0x07, 0xe0, 0x40, 0x00, 0x07, 0xe0, 0x00, 0x00, 0x07, 0xe0, 0x00, 0x00, 0x03, 0xc0, 0x00
};

const unsigned char settings_icon2 [] PROGMEM = {
  // 'Settings-icon-3, 32x32px
  0x00, 0x00, 0x00, 0x00, 0x00, 0x7f, 0xfe, 0x00, 0x01, 0xff, 0xff, 0x80, 0x07, 0xff, 0xff, 0xe0,
  0x0f, 0xff, 0xff, 0xf0, 0x1f, 0xff, 0xff, 0xf8, 0x1f, 0xc7, 0xff, 0xf8, 0x3f, 0xc3, 0xff, 0xfc,
  0x3f, 0xe3, 0xff, 0xfc, 0x7f, 0xe3, 0xff, 0xfe, 0x7c, 0xc3, 0xff, 0xfe, 0x7c, 0x03, 0xff, 0xfe,
  0x7e, 0x01, 0xff, 0xfe, 0x7f, 0x00, 0xff, 0xfe, 0x7f, 0xf8, 0x7f, 0xfe, 0x7f, 0xfc, 0x3f, 0xfe,
  0x7f, 0xfe, 0x1f, 0xfe, 0x7f, 0xff, 0x0f, 0xfe, 0x7f, 0xff, 0x80, 0x7e, 0x7f, 0xff, 0xc0, 0x3e,
  0x7f, 0xff, 0xc0, 0x3e, 0x7f, 0xff, 0xc3, 0x3e, 0x7f, 0xff, 0xc7, 0xfe, 0x3f, 0xff, 0xc7, 0xfc,
  0x3f, 0xff, 0xe3, 0xfc, 0x1f, 0xff, 0xf3, 0xf8, 0x1f, 0xff, 0xff, 0xf8, 0x0f, 0xff, 0xff, 0xf0,
  0x07, 0xff, 0xff, 0xe0, 0x01, 0xff, 0xff, 0x80, 0x00, 0x7f, 0xfe, 0x00, 0x00, 0x00, 0x00, 0x00
};
```
