## Extended Date - LIQ HD Arduino Source Code for "LIQ HD (Lick Instance Quantifier Home cage Device): An open-source tool for recording undisturbed two-bottle drinking behavior in a home cage environment": Pages.ino

```
//240x320 pixel screen

void main_menu_page()
{
  while (display_page == "main") { //show main menu screen

    set_brightness(); // set screen brightness depending on light cycle
    display_time(); //display date and time in corner, refresh every minute

    if (refresh_page) { //do this only once every time main menu is opened
      refresh_page = !refresh_page;
      reset_total_LN();
      timeouts = 0;
      tft.fillRect(0, 20, 240, 320, ILI9341_RED);
      drawBitmap(0, 40, main_logo, 240, 51, ILI9341_WHITE);
      drawBitmap(8, 28, settings_icon, 32, 32, ILI9341_WHITE);

      tft.setFont(&FreeSansBold9pt7b);
      NickText_center("Experimental Side", 120, 110, 1);
      tft.setFont(&FreeSans9pt7b);
      Button_center(60, 150, 80, 40, ILI9341_BLACK, ILI9341_BLACK, ILI9341_WHITE, "LEFT", 1);
      Button_center(180, 150, 80, 40, ILI9341_BLACK, ILI9341_BLACK, ILI9341_WHITE, "RIGHT", 1);
      tft.setFont(&FreeSansBold12pt7b);
      Button_center(120, 220, 140, 60, ILI9341_BLACK, ILI9341_BLACK, ILI9341_WHITE, "START!", 1); //"Start recording" button --> go to recording_page (display_page = 2)

      tft.setFont(&FreeSans9pt7b); //Experimental Side --> left or right?
      if (E_side == "LEFT") {
        Button_center(60, 150, 80, 40, ILI9341_BLACK, ILI9341_GREEN, ILI9341_WHITE, "LEFT", 1);
      }

      if (E_side == "RIGHT") {
        Button_center(180, 150, 80, 40, ILI9341_BLACK, ILI9341_GREEN, ILI9341_WHITE, "RIGHT", 1);
      }

      if (!SD.begin(chipSelect)) { //Info if SD card is instered, if not, --> display button to mount SD card to insert and retest
        NickText(F("SD Card failed..."), 10, 295, 1);
        DEBUG_PRINTLN("SD Card failed to initialize");
        eject_button = false;
        Button_center(190, 290, 90, 30, ILI9341_BLACK, ILI9341_BLACK, ILI9341_WHITE, "Mount SD", 1);
      }
      //error(F("Card failed, or not present"));
      else {
        NickText(F("SD card ready!"), 15, 295, 1);
        eject_button = true;
        //start_SD = false;
        Button_center(190, 290, 90, 30, ILI9341_BLACK, ILI9341_BLACK, ILI9341_WHITE, "Eject SD", 1);
      }
    }

    if (ts.touched()) {
      TS_Point p = ts.getPoint();

      p.x = map(p.x, 0, 240, 0, 240);
      p.y = map(p.y, 0, 320, 0, 320);

      DEBUG_PRINT("(X = "); DEBUG_PRINT(p.x);
      DEBUG_PRINT(", Y = "); DEBUG_PRINT(p.y);
      DEBUG_PRINT(", Pressure = "); DEBUG_PRINT(p.z);
      DEBUG_PRINTLN(")");

      tft.setFont(&FreeSans9pt7b);

      if (p.x > 50 && p.x < 190 && p.y > 190 && p.y < 250) { //Start button coordiantes
        p = {}; //debounce touchscreen
        tft.drawRoundRect(50, 190, 140, 60, 4, ILI9341_GREEN);
        reset_variables();
        if (!SD.begin(chipSelect)) {  //test SD card
          tft.fillRect(0, 20, 240, 320, ILI9341_RED);
          NickText(F("SD Card failed or not inserted"), 2, 100, 1);
          wait(500);
          refresh_page = true;
        }
        else {
          wait(500);
          make_file(); // generate filename to save spreadsheet
          display_page = "recording"; // go to recording_page
          refresh_page = true;
        }
      }

      if (p.x > 20 && p.x < 100 && p.y > 130 && p.y < 170) { //Experimental LEFT button coordiantes
        p = {}; //debounce touchscreen
        E_side = "LEFT"; // Change Experimental side for spreadsheet
        tft.drawRoundRect(20, 130, 80, 40, 4, ILI9341_GREEN);// highlight with green border
        tft.drawRoundRect(140, 130, 80, 40, 4, ILI9341_BLACK);// unhighlight other button
        wait(100);
      }

      if (p.x > 140 && p.x < 220 && p.y > 130 && p.y < 170) { //Experimental RIGHT button coordiantes
        p = {}; //debounce touchscreen
        E_side = "RIGHT"; // Change Experimental side for spreadsheet
        tft.drawRoundRect(20, 130, 80, 40, 4, ILI9341_BLACK);// highlight with green border
        tft.drawRoundRect(140, 130, 80, 40, 4, ILI9341_GREEN);// unhighlight other button
        wait(100);
      }

      if (p.x > 145 && p.x < 235 && p.y > 275 && p.y < 305 && eject_button) { //Eject Button button coordiantes
        p = {}; //debounce touchscreen
        tft.drawRoundRect(145, 275, 90, 30, 4, ILI9341_GREEN);
        SD.end(); // Eject SD card
        tft.fillRect(0, 270, 140, 30, ILI9341_RED);
        eject_button = false;
        wait(1000);
        NickText(F("Safe to remove"), 15, 295, 1);
        Button_center(190, 290, 90, 30, ILI9341_BLACK, ILI9341_BLACK, ILI9341_WHITE, "Mount SD", 1);
      }

      if (p.x > 145 && p.x < 235 && p.y > 275 && p.y < 305 && !eject_button) { //Mount Button button coordiantes
        p = {}; //debounce touchscreen
        tft.drawRoundRect(145, 275, 90, 30, 4, ILI9341_GREEN);
        tft.fillRect(0, 270, 140, 30, ILI9341_RED);
        wait(1000);
        if (!SD.begin(chipSelect)) { //Info if SD card is instered, if not, --> display button to mount SD card to insert and retest
          NickText(F("SD Card failed..."), 10, 295, 1);
          eject_button = false;
          Button_center(190, 290, 90, 30, ILI9341_BLACK, ILI9341_BLACK, ILI9341_WHITE, "Mount SD", 1);
        }
        else {
          NickText(F("SD card ready!"), 15, 295, 1);
          eject_button = true;
          Button_center(190, 290, 90, 30, ILI9341_BLACK, ILI9341_BLACK, ILI9341_WHITE, "Eject SD", 1);
        }
      }

      if (p.x > 0 && p.x < 42 && p.y > 20 && p.y < 70) { //Settings button button coordiantes
        p = {}; //debounce touchscreen
        drawBitmap(8, 28, settings_icon, 32, 32, ILI9341_GREEN);
        wait(500);
        display_page = "settings";
        refresh_page = true;
      }
    }
  }
}

//======================================================================================

void recording_page()
{
  while (display_page == "recording") {

    if (refresh_page) { //do this only once every time coming to recording page
      refresh_page = !refresh_page;
      if (!cap.begin(0x5A)) {
        error("MPR121 #1 not found, check wiring?");
        while (!cap.begin(0x5A)) {
          resetWire();
          if (ts.touched()) {
            TS_Point p = ts.getPoint(); wait(100);
            p.x = map(p.x, 0, 240, 0, 240); p.y = map(p.y, 0, 320, 0, 320);
            if (p.x > 10 && p.x < 230 && p.y > 220 && p.y < 280) {
              p = {}; display_page = "main"; refresh_page = true; break;
            }
          }
        }
      }
      if (!cap2.begin(0x5B)) {
        error("MPR121 #2 not found, check wiring?");
        while (!cap2.begin(0x5B)) {
          resetWire();
          if (ts.touched()) {
            TS_Point p = ts.getPoint(); wait(100);
            p.x = map(p.x, 0, 240, 0, 240); p.y = map(p.y, 0, 320, 0, 320);
            if (p.x > 10 && p.x < 230 && p.y > 220 && p.y < 280) {
              p = {}; display_page = "main"; refresh_page = true; break;
            }
          }
        }
      }
      if (!cap3.begin(0x5C)) {
        error("MPR121 #3 not found, check wiring?");
        while (!cap3.begin(0x5C)) {
          resetWire();
          if (ts.touched()) {
            TS_Point p = ts.getPoint(); wait(100);
            p.x = map(p.x, 0, 240, 0, 240); p.y = map(p.y, 0, 320, 0, 320);
            if (p.x > 10 && p.x < 230 && p.y > 220 && p.y < 280) {
              p = {}; display_page = "main"; refresh_page = true; break;
            }
          }
        }
      }

      if (cap.begin(0x5A) && cap2.begin(0x5B) && cap3.begin(0x5C)) {
        display_licks(); //display lick information
        tft.setFont();
        NickText(filename, 165, 306, 1); //show what file is being recorded to (filename)

        tft.setFont(&FreeSans9pt7b);
        NickText("Exp. on ", 2, 315, 1);
        tft.print(E_side); //Show what side experimental liquid is on

        Button_center(57, 280, 90, 40, ILI9341_BLACK, ILI9341_BLACK, ILI9341_WHITE, "PAUSE", 1);
        Button_center(170, 280, 120, 40, ILI9341_BLACK, ILI9341_BLACK, ILI9341_WHITE, "Save & Quit", 1);

        cap.setThresholds(touch_threshold, release_threshold); //set sensitivity of touch and release of capacitive sensors
        cap2.setThresholds(touch_threshold, release_threshold);
        cap3.setThresholds(touch_threshold, release_threshold);
      }
      else {
        show_time = true;
        break;
      }
    }

    now = rtc.now();
    curr_min = now.minute();
    LOG_COUNTER = now.minute();
    prev_min = now.minute();
    currentMillis = millis();
    previousMillis = currentMillis;
    reset_variables(); //Reset lick variables and timer

    //==========================================================================================
    while (LOG_COUNTER < (prev_min + LOG_INTERVAL)) {
      set_brightness(); //set screen brightness depending on light cycle
      display_time();
      auto_calibration();
      currentMillis = millis();

      if (Wire.getWireTimeoutFlag()) { //if the I2C timesout, rescue the sensors and rtc to continue
        refresh_page = true;
        Wire.clearWireTimeoutFlag();
        timeouts += 1;
        logfile.print("Timeouts,");
        logfile.println(timeouts);
        DEBUG_PRINT("Timeouts, ");
        DEBUG_PRINTLN(timeouts);
        resetWire();
        if (!rtc.begin()) {
          logfile.println(F("rtc failed"));
          DEBUG_PRINTLN(F("rtc failed"));
        }
        break;
      }

      if (!SD.begin(chipSelect)) { //test if SD is working, bring to error page if not
        display_page = "SDerror";
        break;
      }

      Record_Licks(); //record licks from MPR121 sensor #1, 2, & 3

      if (ts.touched()) {
        TS_Point p = ts.getPoint();

        p.x = map(p.x, 0, 240, 0, 240);
        p.y = map(p.y, 0, 320, 0, 320);

        DEBUG_PRINT("(X = "); DEBUG_PRINT(p.x);
        DEBUG_PRINT(", Y = "); DEBUG_PRINT(p.y);
        DEBUG_PRINT(", Pressure = "); DEBUG_PRINT(p.z);
        DEBUG_PRINTLN(")");

        tft.setFont(&FreeSans9pt7b);
        if (p.x > 12 && p.x < 102 && p.y > 260 && p.y < 300) { //PAUSE button coordiantes --> write leftover data to card and eject so you can safely remove SD card --> go to pause_page (display_page = "pause")
          p = {}; //debounce touchscreen
          Button_center(57, 280, 90, 40, ILI9341_BLACK, ILI9341_GREEN, ILI9341_WHITE, "PAUSE", 1);
          logfile.flush();
          DEBUG_PRINT("Writing data to SD card!");
          logfile.close();
          SD.end();
          display_page = "pause"; // go to pause_page
          refresh_page = true;
          wait(100);
          break;
        }

        if (p.x > 110 && p.x < 230 && p.y > 260 && p.y < 300) { //Save & Quit button coordiantes --> write leftover data to card --> return to main_menu_page (display_page = "main")
          p = {}; //debounce touchscreen
          Button_center(170, 280, 120, 40, ILI9341_BLACK, ILI9341_GREEN, ILI9341_WHITE, "Save & Quit", 1);// highlight with green border
          logfile.flush();
          DEBUG_PRINT("Writing data to SD card!");
          logfile.close();
          SD.end();
          display_page = "main"; //go to main_menu_page
          refresh_page = true;
          wait(100);
          break;
        }

        if (p.x > 193 && p.x < 238 && p.y > 46 && p.y < 66) { //refresh button coordiantes
          p = {}; //debounce touchscreen
          display_licks();
        }
      }

      now = rtc.now();
      curr_min = now.minute();
      if (curr_min != prev_min || currentMillis > previousMillis + emergency_counter) {
        LOG_COUNTER += 1;
        SYNC_COUNTER += 1;
        curr_min = now.minute();
        currentMillis = millis();
        previousMillis = currentMillis;
      }
    }
    //==========================================================================================

    if (display_page != "recording") {
      break; //break the loop if page should be changed
    }

    update_sippers(); //update duration of sippers that are in use during data logging

    reset_time_now(); //reset time_now variables to millis()

    write_to_file(); //log data to file

    calc_total_LN(); //calculate total lick duration for each sipper to be diplayed on screen

    write_data(); // Now we write data to disk! Don't sync too often - requires 2048 ints of I/O to SD card, which uses a bunch of power and takes time
  }
}

//======================================================================================

void pause_page()
{
  while (display_page == "pause") {

    display_time(); //show what file was being recorded to (filename), date and time paused, and actual time (refresh every minute)
    set_brightness(); //set screen brightness depending on light cycle

    if (refresh_page) {
      refresh_page = !refresh_page;
      tft.fillRect(0, 20, 240, 320, ILI9341_RED);
      tft.setFont(&FreeSans18pt7b);
      tft.setTextColor(ILI9341_WHITE);
      NickText_center("PAUSED", 120, 80, 1);
      tft.setFont(&FreeSans12pt7b);
      wait(1000);
      NickText_center("Safe to remove", 120, 120, 1);
      NickText_center("SD card!", 120, 150, 1);
      tft.setFont(&FreeSans9pt7b);
      NickText_center("Reinsert SD before resuming", 120, 180, 1);

      tft.setFont(&FreeSans12pt7b);
      Button_center(120, 250, 110, 40, ILI9341_BLACK, ILI9341_BLACK, ILI9341_WHITE, "Resume", 1);
    }

    if (ts.touched()) {
      TS_Point p = ts.getPoint();

      // serial monitor will show when and where touched
      // Scale from ~0->4000 to tft.width using the calibration #'s
      p.x = map(p.x, 0, 240, 0, 240);
      p.y = map(p.y, 0, 320, 0, 320);

      DEBUG_PRINT("(X = "); DEBUG_PRINT(p.x);
      DEBUG_PRINT(", Y = "); DEBUG_PRINT(p.y);
      DEBUG_PRINT(", Pressure = "); DEBUG_PRINT(p.z);
      DEBUG_PRINTLN(")");

      tft.setFont(&FreeSans9pt7b);

      if (p.x > 65 && p.x < 175 && p.y > 230 && p.y < 270) { //Resume button coordiantes, Unpause --> mount SD card and continue recording to the same file --> go to recording_page (display_page = "recording")
        p = {}; //debounce touchscreen
        tft.drawRoundRect(65, 230, 110, 40, 4, ILI9341_GREEN);
        wait(100);
        if (!SD.begin(chipSelect)) {
          display_page = "SDerror";
          break;
        }
        logfile = SD.open(filename, FILE_WRITE);
        logfile.seek(EOF);
        logfile.println("PAUSED");
        DEBUG_PRINTLN("PAUSED");
        logging_same_file();
        display_page = "recording"; // go to recording_page
        refresh_page = true;
      }
    }
  }
}

//======================================================================================

void SD_error_page()
{
  while (display_page == "SDerror") {

    display_time();
    set_brightness();

    if (refresh_page) {
      refresh_page = !refresh_page;
      now = rtc.now();
      tft.fillRect(0, 20, 240, 320, ILI9341_RED);
      tft.setFont(&FreeSans9pt7b);
      NickText_center(F("SD Card failed or not inserted"), 120, 100, 1);
      NickText(F("Card failed at:"), 0, 150, 1); //dispaly time that SD failed
      tft.println();
      tft.print(now.month());
      tft.print("/");
      tft.print(now.day());
      tft.print("/");
      tft.print(now.year());
      tft.print(" ");
      tft.print(now.hour());
      tft.print(":");
      if (now.minute() < 10) {
        tft.print("0");
      }
      tft.print(now.minute());
      Button_center(120, 250, 220, 60, ILI9341_BLACK, ILI9341_BLACK, ILI9341_WHITE, "Return to Main Menu", 1);
    }

    if (ts.touched()) {
      TS_Point p = ts.getPoint();

      p.x = map(p.x, 0, 240, 0, 240);
      p.y = map(p.y, 0, 320, 0, 320);

      DEBUG_PRINT("(X = "); DEBUG_PRINT(p.x);
      DEBUG_PRINT(", Y = "); DEBUG_PRINT(p.y);
      DEBUG_PRINT(", Pressure = "); DEBUG_PRINT(p.z);
      DEBUG_PRINTLN(")");

      if (p.x > 10 && p.x < 230 && p.y > 220 && p.y < 280) { //Return to Main Menu button coordiantes // fix this
        p = {}; //debounce touchscreen
        Button_center(120, 250, 220, 60, ILI9341_BLACK, ILI9341_GREEN, ILI9341_WHITE, "Return to Main Menu", 1);
        wait(100);
        display_page = "main"; // go to main_menu_page
        refresh_page = true;
      }
    }
  }
}

//======================================================================================

void settings_page()
{
  while (display_page == "settings") {

    display_time();
    set_brightness();

    if (refresh_page) {
      refresh_page = !refresh_page;

      tft.fillRect(0, 20, 240, 320, ILI9341_RED);
      tft.setFont(&FreeSansBold12pt7b);
      NickText_center("Settings", 120, 40, 1);
      tft.setFont(&FreeSans9pt7b);
      NickText("Date/Time", 10, 85, 1); Button_center(130, 80, 60, 26, ILI9341_BLACK, ILI9341_BLACK, ILI9341_WHITE, "Edit", 1);
      NickText("Lights ON Time", 10, 115, 1); Button_center(160, 110, 35, 26, ILI9341_BLACK, ILI9341_BLACK, ILI9341_WHITE, lights_on, 1);
      NickText("Lights OFF Time", 10, 145, 1); Button_center(168, 140, 35, 26, ILI9341_BLACK, ILI9341_BLACK, ILI9341_WHITE, lights_off, 1);
      NickText("Sensor Settings", 10, 175, 1); Button_center(175, 170, 60, 26, ILI9341_BLACK, ILI9341_BLACK, ILI9341_WHITE, "Edit", 1);
      NickText("Parameters to Log", 10, 205, 1); Button_center(195, 200, 60, 26, ILI9341_BLACK, ILI9341_BLACK, ILI9341_WHITE, "Edit", 1);
      NickText("Time Bin Size", 10, 235, 1); Button_center(145, 230, 35, 26, ILI9341_BLACK, ILI9341_BLACK, ILI9341_WHITE, LOG_INTERVAL, 1);
      NickText("Sync Interval", 10, 265, 1); Button_center(137, 260, 35, 26, ILI9341_BLACK, ILI9341_BLACK, ILI9341_WHITE, SYNC_INTERVAL, 1);
      Button_center(180, 295, 60, 40, ILI9341_BLACK, ILI9341_BLACK, ILI9341_WHITE, "Save", 1);
      Button_center(60, 295, 60, 40, ILI9341_BLACK, ILI9341_BLACK, ILI9341_WHITE, "Reset", 1);
    }

    if (ts.touched()) {
      TS_Point p = ts.getPoint();

      p.x = map(p.x, 0, 240, 0, 240);
      p.y = map(p.y, 0, 320, 0, 320);

      DEBUG_PRINT("(X = "); DEBUG_PRINT(p.x);
      DEBUG_PRINT(", Y = "); DEBUG_PRINT(p.y);
      DEBUG_PRINT(", Pressure = "); DEBUG_PRINT(p.z);
      DEBUG_PRINTLN(")");

      if (p.x > 100 && p.x < 160 && p.y > 67 && p.y < 93) { //Date/Time Edit
        p = {}; //debounce touchscreen
        Button_center(130, 80, 60, 26, ILI9341_BLACK, ILI9341_GREEN, ILI9341_WHITE, "Edit", 1);
        wait(100);
        display_page = "settings_datetime";
        refresh_page = true;
        show_time = true;

      }
      if (p.x > 143 && p.x < 178 && p.y > 97 && p.y < 123) { //Lights ON
        p = {}; //debounce touchscreen
        Button_center(160, 110, 35, 26, ILI9341_BLACK, ILI9341_GREEN, ILI9341_WHITE, lights_on, 1);
        wait(100);
        GetNum("Lights ON Hour", lights_on);
        if (lights_on >= 24) {
          lights_on = default_lights_on;
        }
        ok = false;
        refresh_page = true;
        show_time = true;
      }
      if (p.x > 151 && p.x < 186 && p.y > 127 && p.y < 153) { //Lights OFF
        p = {}; //debounce touchscreen
        Button_center(168, 140, 35, 26, ILI9341_BLACK, ILI9341_GREEN, ILI9341_WHITE, lights_off, 1);
        wait(100);
        GetNum("Lights OFF Hour", lights_off);
        if (lights_off >= 24) {
          lights_off = default_lights_off;
        }
        ok = false;
        refresh_page = true;
        show_time = true;
      }
      if (p.x > 145 && p.x < 205 && p.y > 157 && p.y < 183) { //Sensor Settings Edit
        p = {}; //debounce touchscreen
        Button_center(175, 170, 60, 26, ILI9341_BLACK, ILI9341_GREEN, ILI9341_WHITE, "Edit", 1);
        wait(100);
        display_page = "settings_sensor";
        refresh_page = true;
        show_time = true;
      }
      if (p.x > 165 && p.x < 225 && p.y > 187 && p.y < 213) { //Parameters to log Edit
        p = {}; //debounce touchscreen
        Button_center(195, 200, 60, 26, ILI9341_BLACK, ILI9341_GREEN, ILI9341_WHITE, "Edit", 1);
        wait(100);
        display_page = "settings_parameters";
        refresh_page = true;
        show_time = true;
      }
      if (p.x > 128 && p.x < 163 && p.y > 217 && p.y < 243) { //Time bin size
        p = {}; //debounce touchscreen
        Button_center(145, 230, 35, 26, ILI9341_BLACK, ILI9341_GREEN, ILI9341_WHITE, LOG_INTERVAL, 1);
        wait(100);
        GetNum("Time Bin Size (min)", LOG_INTERVAL);
        if (SYNC_INTERVAL < LOG_INTERVAL) {
          SYNC_INTERVAL = LOG_INTERVAL;
        }
        ok = false;
        refresh_page = true;
        show_time = true;
      }
      if (p.x > 120 && p.x < 155 && p.y > 247 && p.y < 273) { //Sync interval
        p = {}; //debounce touchscreen
        Button_center(137, 260, 35, 26, ILI9341_BLACK, ILI9341_GREEN, ILI9341_WHITE, SYNC_INTERVAL, 1);
        wait(100);
        GetNum("Sync Interval (min)", SYNC_INTERVAL);
        if (SYNC_INTERVAL < LOG_INTERVAL) {
          SYNC_INTERVAL = LOG_INTERVAL;
        }
        ok = false;
        refresh_page = true;
        show_time = true;
      }
      if (p.x > 150 && p.x < 210 && p.y > 275 && p.y < 315) { //Save
        p = {}; //debounce touchscreen
        Button_center(180, 295, 60, 40, ILI9341_BLACK, ILI9341_GREEN, ILI9341_WHITE, "Save", 1);
        wait(100);
        display_page = "main"; // go to main_menu_page
        refresh_page = true;
        show_time = true;
      }
      if (p.x > 30 && p.x < 90 && p.y > 275 && p.y < 315) { //Reset
        p = {}; //debounce touchscreen
        Button_center(60, 295, 60, 40, ILI9341_BLACK, ILI9341_GREEN, ILI9341_WHITE, "Reset", 1);
        wait(100);
        lights_on = default_lights_on;
        lights_off = default_lights_off;
        LOG_INTERVAL = default_LOG_INTERVAL;
        SYNC_INTERVAL = dafault_SYNC_INTERVAL;
        refresh_page = true;
        show_time = true;
      }
    }
  }

  while (display_page == "settings_datetime") {
    display_time();
    set_brightness();

    if (refresh_page) {
      refresh_page = !refresh_page;
      DateTime now = rtc.now();
      tft.fillRect(0, 20, 240, 320, ILI9341_RED);
      tft.setFont(&FreeSansBold12pt7b);
      NickText_center("Date/Time Settings", 120, 40, 1);
      tft.setFont(&FreeSans9pt7b);
      NickText("Month", 10, 85, 1); Button_center(90, 80, 35, 26, ILI9341_BLACK, ILI9341_BLACK, ILI9341_WHITE, now.month(), 1);
      NickText("Day", 10, 115, 1); Button_center(90, 110, 35, 26, ILI9341_BLACK, ILI9341_BLACK, ILI9341_WHITE, now.day(), 1);
      NickText("Year", 10, 145, 1); Button_center(90, 140, 60, 26, ILI9341_BLACK, ILI9341_BLACK, ILI9341_WHITE, now.year(), 1);
      NickText("Hour", 10, 205, 1); Button_center(90, 200, 35, 26, ILI9341_BLACK, ILI9341_BLACK, ILI9341_WHITE, now.hour(), 1);
      NickText("Minute", 10, 235, 1); Button_center(90, 230, 35, 26, ILI9341_BLACK, ILI9341_BLACK, ILI9341_WHITE, now.minute(), 1);
      Button_center(180, 295, 60, 40, ILI9341_BLACK, ILI9341_BLACK, ILI9341_WHITE, "Save", 1);
    }

    if (ts.touched()) {
      TS_Point p = ts.getPoint();

      p.x = map(p.x, 0, 240, 0, 240);
      p.y = map(p.y, 0, 320, 0, 320);

      DEBUG_PRINT("(X = "); DEBUG_PRINT(p.x);
      DEBUG_PRINT(", Y = "); DEBUG_PRINT(p.y);
      DEBUG_PRINT(", Pressure = "); DEBUG_PRINT(p.z);
      DEBUG_PRINTLN(")");

      if (p.x > 73 && p.x < 108 && p.y > 67 && p.y < 93) { //month
        p = {}; //debounce touchscreen
        Button_center(90, 80, 35, 26, ILI9341_BLACK, ILI9341_GREEN, ILI9341_WHITE, now.month(), 1);
        wait(100);
        int new_month = now.month();
        GetNum("Month", new_month);
        if (new_month > 12) {
          new_month = 12;
        }
        rtc.adjust(DateTime(now.year(), new_month, now.day(), now.hour(), now.minute(), 0));
        ok = false;
        refresh_page = true;
        show_time = true;
      }
      if (p.x > 73 && p.x < 108 && p.y > 97 && p.y < 123) { //day
        p = {}; //debounce touchscreen
        Button_center(90, 110, 35, 26, ILI9341_BLACK, ILI9341_GREEN, ILI9341_WHITE, now.day(), 1);
        wait(100);
        int new_day = now.day();
        GetNum("Day", new_day);
        if (new_day > 31) {
          new_day = 31;
        }
        rtc.adjust(DateTime(now.year(), now.month(), new_day, now.hour(), now.minute(), 0));
        ok = false;
        refresh_page = true;
        show_time = true;
      }
      if (p.x > 60 && p.x < 120 && p.y > 127 && p.y < 153) { //year
        p = {}; //debounce touchscreen
        Button_center(90, 140, 60, 26, ILI9341_BLACK, ILI9341_GREEN, ILI9341_WHITE, now.year(), 1);
        wait(100);
        int new_year = now.year();
        GetNum("Year", new_year);
        if (new_year > 2099) {
          new_year = 2099;
        }
        rtc.adjust(DateTime(new_year, now.month(), now.day(), now.hour(), now.minute(), 0));
        ok = false;
        refresh_page = true;
        show_time = true;
      }
      if (p.x > 73 && p.x < 108 && p.y > 187 && p.y < 213) { //hour
        p = {}; //debounce touchscreen
        Button_center(90, 200, 35, 26, ILI9341_BLACK, ILI9341_GREEN, ILI9341_WHITE, now.hour(), 1);
        wait(100);
        int new_hour = now.hour();
        GetNum("Hour", new_hour);
        if (new_hour > 23) {
          new_hour = 0;
        }
        rtc.adjust(DateTime(now.year(), now.month(), now.day(), new_hour, now.minute(), 0));
        ok = false;
        refresh_page = true;
        show_time = true;
      }
      if (p.x > 73 && p.x < 108 && p.y > 217 && p.y < 243) { //minute
        p = {}; //debounce touchscreen
        Button_center(90, 230, 35, 26, ILI9341_BLACK, ILI9341_GREEN, ILI9341_WHITE, now.minute(), 1);
        wait(100);
        int new_minute = now.minute();
        GetNum("Minute", new_minute);
        if (new_minute > 59) {
          new_minute = 0;
        }
        rtc.adjust(DateTime(now.year(), now.month(), now.day(), now.hour(), new_minute, 0));
        ok = false;
        refresh_page = true;
        show_time = true;
      }
      if (p.x > 150 && p.x < 210 && p.y > 275 && p.y < 315) { //Save
        p = {}; //debounce touchscreen
        Button_center(180, 295, 60, 40, ILI9341_BLACK, ILI9341_GREEN, ILI9341_WHITE, "Save", 1);
        wait(100);
        display_page = "settings";
        refresh_page = true;
        show_time = true;
        break;
      }
    }
  }

  while (display_page == "settings_parameters") {
    display_time();
    set_brightness();

    if (refresh_page) {
      refresh_page = !refresh_page;
      tft.fillRect(0, 20, 240, 320, ILI9341_RED);
      tft.setFont(&FreeSansBold12pt7b);
      NickText_center("Parameters to Log", 120, 40, 1);
      tft.setFont(&FreeSans9pt7b);
      if (log_LN) {
        logging[0] = "X";
      } else {
        logging[0] = "";
      }
      if (log_LD) {
        logging[1] = "X";
      } else {
        logging[1] = "";
      }
      if (log_BN) {
        logging[2] = "X";
      } else {
        logging[2] = "";
      }
      if (log_BD) {
        logging[3] = "X";
      } else {
        logging[3] = "";
      }
      if (log_BLN) {
        logging[4] = "X";
      } else {
        logging[4] = "";
      }
      if (log_BLD) {
        logging[5] = "X";
      } else {
        logging[5] = "";
      }
      NickText("Lick Number", 44, 85, 1); tft.setFont(&FreeSansBold12pt7b); Button_center(22, 80, 24, 24, ILI9341_BLACK, ILI9341_BLACK, ILI9341_WHITE, logging[0], 1); tft.setFont(&FreeSans9pt7b);
      NickText("Lick Duration", 44, 115, 1); tft.setFont(&FreeSansBold12pt7b); Button_center(22, 110, 24, 24, ILI9341_BLACK, ILI9341_BLACK, ILI9341_WHITE, logging[1], 1); tft.setFont(&FreeSans9pt7b);
      NickText("Bout Number", 44, 145, 1); tft.setFont(&FreeSansBold12pt7b); Button_center(22, 140, 24, 24, ILI9341_BLACK, ILI9341_BLACK, ILI9341_WHITE, logging[2], 1); tft.setFont(&FreeSans9pt7b);
      NickText("Bout Duration", 44, 175, 1); tft.setFont(&FreeSansBold12pt7b); Button_center(22, 170, 24, 24, ILI9341_BLACK, ILI9341_BLACK, ILI9341_WHITE, logging[3], 1); tft.setFont(&FreeSans9pt7b);
      NickText("Bout Lick Number", 44, 205, 1); tft.setFont(&FreeSansBold12pt7b); Button_center(22, 200, 24, 24, ILI9341_BLACK, ILI9341_BLACK, ILI9341_WHITE, logging[4], 1); tft.setFont(&FreeSans9pt7b);
      NickText("Bout Lick Duration", 44, 235, 1); tft.setFont(&FreeSansBold12pt7b); Button_center(22, 230, 24, 24, ILI9341_BLACK, ILI9341_BLACK, ILI9341_WHITE, logging[5], 1); tft.setFont(&FreeSans9pt7b);
      Button_center(180, 295, 60, 40, ILI9341_BLACK, ILI9341_BLACK, ILI9341_WHITE, "Save", 1);
      Button_center(60, 295, 60, 40, ILI9341_BLACK, ILI9341_BLACK, ILI9341_WHITE, "Reset", 1);
    }

    if (ts.touched()) {
      TS_Point p = ts.getPoint();

      p.x = map(p.x, 0, 240, 0, 240);
      p.y = map(p.y, 0, 320, 0, 320);

      DEBUG_PRINT("(X = "); DEBUG_PRINT(p.x);
      DEBUG_PRINT(", Y = "); DEBUG_PRINT(p.y);
      DEBUG_PRINT(", Pressure = "); DEBUG_PRINT(p.z);
      DEBUG_PRINTLN(")");

      if (p.x < 190 && p.y > 68 && p.y < 92) { //LN
        p = {}; //debounce touchscreen
        log_LN = !log_LN;
        if (log_LN) {
          logging[0] = "X";
        } else {
          logging[0] = "";
        }
        tft.setFont(&FreeSansBold12pt7b);
        Button_center(22, 80, 24, 24, ILI9341_BLACK, ILI9341_BLACK, ILI9341_WHITE, logging[0], 1);
        wait(100);
      }
      if (p.x < 190 && p.y > 98 && p.y < 122) { //LD
        p = {}; //debounce touchscreen
        log_LD = !log_LD;
        if (log_LD) {
          logging[1] = "X";
        } else {
          logging[1] = "";
        }
        tft.setFont(&FreeSansBold12pt7b);
        Button_center(22, 110, 24, 24, ILI9341_BLACK, ILI9341_BLACK, ILI9341_WHITE, logging[1], 1);
        wait(100);
      }
      if (p.x < 190 && p.y > 128 && p.y < 152) { //BN
        p = {}; //debounce touchscreen
        log_BN = !log_BN;
        if (log_BN) {
          logging[2] = "X";
        } else {
          logging[2] = "";
        }
        tft.setFont(&FreeSansBold12pt7b);
        Button_center(22, 140, 24, 24, ILI9341_BLACK, ILI9341_BLACK, ILI9341_WHITE, logging[2], 1);
        wait(100);
      }
      if (p.x < 190 && p.y > 158 && p.y < 182) { //BD
        p = {}; //debounce touchscreen
        log_BD = !log_BD;
        if (log_BD) {
          logging[3] = "X";
        } else {
          logging[3] = "";
        }
        tft.setFont(&FreeSansBold12pt7b);
        Button_center(22, 170, 24, 24, ILI9341_BLACK, ILI9341_BLACK, ILI9341_WHITE, logging[3], 1);
        wait(100);
      }
      if (p.x < 190 && p.y > 188 && p.y < 212) { //BLN
        p = {}; //debounce touchscreen
        log_BLN = !log_BLN;
        if (log_BLN) {
          logging[4] = "X";
        } else {
          logging[4] = "";
        }
        tft.setFont(&FreeSansBold12pt7b);
        Button_center(22, 200, 24, 24, ILI9341_BLACK, ILI9341_BLACK, ILI9341_WHITE, logging[4], 1);
        wait(100);
      }
      if (p.x < 190 && p.y > 218 && p.y < 242) { //BLD
        p = {}; //debounce touchscreen
        log_BLD = !log_BLD;
        if (log_BLD) {
          logging[5] = "X";
        } else {
          logging[5] = "";
        }
        tft.setFont(&FreeSansBold12pt7b);
        Button_center(22, 230, 24, 24, ILI9341_BLACK, ILI9341_BLACK, ILI9341_WHITE, logging[5], 1);
        wait(100);
      }
      if (p.x > 150 && p.x < 210 && p.y > 275 && p.y < 315) { //Save
        p = {}; //debounce touchscreen
        tft.setFont(&FreeSans9pt7b);
        Button_center(180, 295, 60, 40, ILI9341_BLACK, ILI9341_GREEN, ILI9341_WHITE, "Save", 1);
        wait(100);
        display_page = "settings";
        refresh_page = true;
      }
      if (p.x > 30 && p.x < 90 && p.y > 275 && p.y < 315) { //Reset
        p = {}; //debounce touchscreen
        tft.setFont(&FreeSans9pt7b);
        Button_center(60, 295, 60, 40, ILI9341_BLACK, ILI9341_GREEN, ILI9341_WHITE, "Reset", 1);
        wait(100);
        log_LN = true;
        log_LD = true;
        log_BN = true;
        log_BD = true;
        log_BLN = true;
        log_BLD = true;
        refresh_page = true;
      }
    }
  }

  while (display_page == "settings_sensor") {
    display_time();
    set_brightness();
    String X;
    if (refresh_page) {
      refresh_page = !refresh_page;
      DateTime now = rtc.now();
      tft.fillRect(0, 20, 240, 320, ILI9341_RED);
      tft.setFont(&FreeSansBold12pt7b);
      NickText_center("Sensor Settings", 120, 40, 1);
      if (auto_cal) {
        X = "X";
      } else {
        X = "";
      }
      tft.setFont(&FreeSans9pt7b);
      NickText("Touch Threshold", 10, 85, 1); Button_center(170, 80, 35, 26, ILI9341_BLACK, ILI9341_BLACK, ILI9341_WHITE, touch_threshold, 1);
      NickText("Release Threshold", 10, 115, 1); Button_center(185, 110, 35, 26, ILI9341_BLACK, ILI9341_BLACK, ILI9341_WHITE, release_threshold, 1);
      NickText("Auto Calibrate?", 10, 145, 1); tft.setFont(&FreeSansBold12pt7b); Button_center(150, 140, 24, 24, ILI9341_BLACK, ILI9341_BLACK, ILI9341_WHITE, X, 1);  tft.setFont(&FreeSans9pt7b);
      if (auto_cal) {
        NickText("Time (24hr)", 30, 175, 1); Button_center(150, 170, 35, 26, ILI9341_BLACK, ILI9341_BLACK, ILI9341_WHITE, auto_cal_time, 1);
        NickText("Time since last lick (sec) ", 30, 205, 1); Button_center(120, 225, 40, 26, ILI9341_BLACK, ILI9341_BLACK, ILI9341_WHITE, auto_cal_sec_since_last_lick, 1);
        tft.setFont();
        NickText_center("Sensors will only auto calibrate once", 120, 240, 1);
        NickText_center("at the specified hour and if no licks", 120, 250, 1);
        NickText_center("have occured within the specified time.", 120, 260, 1);
      }
      tft.setFont(&FreeSans9pt7b);
      Button_center(180, 295, 60, 40, ILI9341_BLACK, ILI9341_BLACK, ILI9341_WHITE, "Save", 1);
      Button_center(60, 295, 60, 40, ILI9341_BLACK, ILI9341_BLACK, ILI9341_WHITE, "Reset", 1);
    }

    if (ts.touched()) {
      TS_Point p = ts.getPoint();

      p.x = map(p.x, 0, 240, 0, 240);
      p.y = map(p.y, 0, 320, 0, 320);

      DEBUG_PRINT("(X = "); DEBUG_PRINT(p.x);
      DEBUG_PRINT(", Y = "); DEBUG_PRINT(p.y);
      DEBUG_PRINT(", Pressure = "); DEBUG_PRINT(p.z);
      DEBUG_PRINTLN(")");

      if (p.x > 153 && p.x < 188 && p.y > 67 && p.y < 93) { //touch threshold
        p = {}; //debounce touchscreen
        tft.setFont(&FreeSans9pt7b);
        Button_center(170, 80, 35, 26, ILI9341_BLACK, ILI9341_GREEN, ILI9341_WHITE, touch_threshold, 1);
        wait(100);
        GetNum("Touch Threshold", touch_threshold);
        if (touch_threshold > 255) {
          touch_threshold = 255;
        }
        ok = false;
        refresh_page = true;
        show_time = true;
      }

      if (p.x > 168 && p.x < 203 && p.y > 97 && p.y < 123) { //release threshold
        p = {}; //debounce touchscreen
        tft.setFont(&FreeSans9pt7b);
        Button_center(185, 110, 35, 26, ILI9341_BLACK, ILI9341_GREEN, ILI9341_WHITE, release_threshold, 1);
        wait(100);
        GetNum("Release Threshold", release_threshold);
        if (release_threshold > 255) {
          release_threshold = 255;
        }
        ok = false;
        refresh_page = true;
        show_time = true;
      }
      if (p.x > 138 && p.x < 162 && p.y > 128 && p.y < 152) { //auto calibrate?
        p = {}; //debounce touchscreen
        auto_cal = !auto_cal;
        wait(100);
        refresh_page = true;
        show_time = true;
      }
      if (p.x > 133 && p.x < 168 && p.y > 157 && p.y < 183 && auto_cal) { //Time to cal
        p = {}; //debounce touchscreen
        tft.setFont(&FreeSans9pt7b);
        Button_center(150, 170, 35, 26, ILI9341_BLACK, ILI9341_GREEN, ILI9341_WHITE, auto_cal_time, 1);
        wait(100);
        GetNum("Auto Cal Time", auto_cal_time);
        if (auto_cal_time > 23) {
          auto_cal_time = 0;
        }
        ok = false;
        refresh_page = true;
        show_time = true;
      }
      if (p.x > 85 && p.x < 155 && p.y > 217 && p.y < 243 && auto_cal) { //time since last cal
        p = {}; //debounce touchscreen
        tft.setFont(&FreeSans9pt7b);
        Button_center(120, 225, 40, 26, ILI9341_BLACK, ILI9341_GREEN, ILI9341_WHITE, auto_cal_sec_since_last_lick, 1);
        wait(100);
        GetNum("Sec since last lick", auto_cal_sec_since_last_lick);
        ok = false;
        refresh_page = true;
        show_time = true;
      }
      if (p.x > 150 && p.x < 210 && p.y > 275 && p.y < 315) { //Save
        p = {}; //debounce touchscreen
        tft.setFont(&FreeSans9pt7b);
        Button_center(180, 295, 60, 40, ILI9341_BLACK, ILI9341_GREEN, ILI9341_WHITE, "Save", 1);
        wait(100);
        display_page = "settings";
        refresh_page = true;
        show_time = true;
        break;
      }
      if (p.x > 30 && p.x < 90 && p.y > 275 && p.y < 315) { //Reset
        p = {}; //debounce touchscreen
        tft.setFont(&FreeSans9pt7b);
        Button_center(60, 295, 60, 40, ILI9341_BLACK, ILI9341_GREEN, ILI9341_WHITE, "Reset", 1);
        wait(100);
        default_touch_threshold = touch_threshold;
        default_release_threshold = release_threshold;
        auto_cal = default_auto_cal;
        auto_cal_time = default_auto_cal_time;
        auto_cal_sec_since_last_lick = default_auto_cal_sec_since_last_lick;
        refresh_page = true;
        show_time = true;
      }
    }
  }
}
```
