## Extended Date - LIQ HD Arduino Source Code for "LIQ HD (Lick Instance Quantifier Home cage Device): An open-source tool for recording undisturbed two-bottle drinking behavior in a home cage environment": Screen_functions.ino

```
//==========================================================================================

void start_display()
{
  tft.begin();
  if (!ts.begin()) {
    DEBUG_PRINTLN("Couldn't start touchscreen controller");
    while (1);
  }
  DEBUG_PRINTLN("Touchscreen started");
}

//==========================================================================================

void WelcomeScreen() //work on this with new name and logo design
{
  tft.setRotation(2);
  tft.fillScreen(ILI9341_BLACK);
  drawBitmap(0, 60, welcome_logo, 240, 173, ILI9341_WHITE);
  wait(2000);
}

//==========================================================================================

void set_brightness()
{
  // fetch the time
  hour_now = now.hour();

  if (lights_on < lights_off) {
    if (hour_now >= lights_on && hour_now < lights_off) {
      analogWrite(5, 255);
    }
    if (hour_now < lights_on || hour_now >= lights_off) {
      analogWrite(5, 10);
    }
  }

  if (lights_on > lights_off) {
    if (hour_now >= lights_on || hour_now < lights_off) {
      analogWrite(5, 255);
    }
    if (hour_now < lights_on && hour_now >= lights_off) {
      analogWrite(5, 10);
    }
  }
}

//======================================================================================

void NickText(String text, int x, int y, int size)
{
  tft.setTextSize(size);
  tft.setCursor(x, y);
  tft.print(text);
}

//======================================================================================

void NickText(int w, int x, int y, int size)
{
  tft.setTextSize(size);
  tft.setCursor(x, y);
  tft.print(w);
}

//======================================================================================

void NickText_center(String text, int center_x, int center_y, int size)
{
  tft.setTextSize(size);
  int16_t x1; int16_t y1; uint16_t w; uint16_t h;
  tft.getTextBounds(text, 0, 320, &x1, &y1, &w, &h);
  int xx = (center_x - 1) - (w / 2);
  int yy = (center_y - 1) + (h / 2);
  tft.setCursor(xx, yy);
  tft.print(text);
}


//======================================================================================

void NickText_center(int variable, int center_x, int center_y, int size)
{
  tft.setTextSize(size);
  int16_t x1; int16_t y1; uint16_t w; uint16_t h;
  String text = String(variable);
  tft.getTextBounds(text, 0, 320, &x1, &y1, &w, &h);
  int xx = (center_x - 1) - (w / 2);
  int yy = (center_y - 1) + (h / 2);
  tft.setCursor(xx, yy);
  tft.print(variable);
}

//==========================================================================================

void Button_center(int center_x, int center_y, int sizeX, int sizeY, uint16_t button_color, uint16_t border_color, uint16_t text_color, String text, int textsize)
{
  int16_t x = center_x - (sizeX / 2);
  int16_t y = center_y - (sizeY / 2);
  tft.fillRoundRect(x, y, sizeX, sizeY, 4, button_color);
  tft.drawRoundRect(x, y, sizeX, sizeY, 4, border_color);
  int16_t x1; int16_t y1; uint16_t w; uint16_t h;
  tft.getTextBounds(text, 0, 320, &x1, &y1, &w, &h);
  int xx = (center_x - 1) - (w / 2);
  int yy = (center_y - 1) + (h / 2);
  tft.setTextColor(text_color);
  tft.setTextSize(textsize);
  tft.setCursor(xx, yy);
  tft.print(text);

  int x_min; int x_max; int y_min; int y_max;
  x_min = x;
  x_max = x + sizeX;
  y_min = y;
  y_max = y + sizeY;
  DEBUG_PRINTLN();
  DEBUG_PRINT(text);
  DEBUG_PRINT(" button borders: p.x>");
  DEBUG_PRINT(x_min);
  DEBUG_PRINT(" && p.x<");
  DEBUG_PRINT(x_max);
  DEBUG_PRINT(" && p.y>");
  DEBUG_PRINT(y_min);
  DEBUG_PRINT(" && p.y<");
  DEBUG_PRINT(y_max);
  DEBUG_PRINT("");
}

//==========================================================================================

void Button_center(int center_x, int center_y, int sizeX, int sizeY, uint16_t button_color, uint16_t border_color, uint16_t text_color, int text, int textsize)
{
  int16_t x = center_x - (sizeX / 2);
  int16_t y = center_y - (sizeY / 2);
  tft.fillRoundRect(x, y, sizeX, sizeY, 4, button_color);
  tft.drawRoundRect(x, y, sizeX, sizeY, 4, border_color);
  int16_t x1; int16_t y1; uint16_t w; uint16_t h;
  tft.getTextBounds(String(text), 0, 320, &x1, &y1, &w, &h);
  int xx = (center_x - 1) - (w / 2);
  int yy = (center_y - 1) + (h / 2);
  tft.setTextColor(text_color);
  tft.setTextSize(textsize);
  tft.setCursor(xx, yy);
  tft.print(text);

  int x_min; int x_max; int y_min; int y_max;
  x_min = x;
  x_max = x + sizeX;
  y_min = y;
  y_max = y + sizeY;
  DEBUG_PRINTLN();
  DEBUG_PRINT(text);
  DEBUG_PRINT(" button borders: p.x>");
  DEBUG_PRINT(x_min);
  DEBUG_PRINT(" && p.x<");
  DEBUG_PRINT(x_max);
  DEBUG_PRINT(" && p.y>");
  DEBUG_PRINT(y_min);
  DEBUG_PRINT(" && p.y<");
  DEBUG_PRINT(y_max);
  DEBUG_PRINT("");
}

//==========================================================================================

void Button(int x, int y, int sizeX, int sizeY, int rad, uint16_t color)
{
  tft.fillRoundRect(x, y, sizeX, sizeY, rad, color);
  tft.drawRoundRect(x, y, sizeX, sizeY, rad, ILI9341_BLACK);
}

//======================================================================================

void error(String str)
{
  show_time = true;
  display_time();
  tft.fillRect(0, 20, 240, 320, ILI9341_RED);
  DEBUG_PRINT("error: ");
  DEBUG_PRINTLN(str);
  tft.setFont();
  tft.setTextWrap(true);
  tft.setFont(&FreeSans9pt7b);
  NickText(str, 5, 50, 1);
  Button_center(120, 250, 220, 60, ILI9341_BLACK, ILI9341_BLACK, ILI9341_WHITE, "Return to Main Menu", 1);
}

//======================================================================================

void display_licks()
{
  tft.setFont(&FreeSans9pt7b);
  tft.fillRect(0, 20, 240, 222, ILI9341_RED);
  NickText_center("Total lick number", 100, 32, 1);
  NickText_center("for each cage (L-R)", 105, 51, 1);
  tft.setFont();
  Button_center(215, 56, 45, 20, ILI9341_BLACK, ILI9341_BLACK, ILI9341_WHITE, "", 1); //refresh button
  NickText_center("Refresh", 217, 49, 1);

  int spacing = 20;
  int x1 = 4;
  int x2 = 120;
  int y1 = 80;
  int y2 = y1 + spacing;
  int y3 = y2 + spacing;
  int y4 = y3 + spacing;
  int y5 = y4 + spacing;
  int y6 = y5 + spacing;
  int y7 = y6 + spacing;
  int y8 = y7 + spacing;
  int y9 = y8 + spacing;

  tft.setFont(&FreeSans9pt7b);
  NickText(F("1"), x1, y1, 0);
  tft.print(F("-"));
  tft.print(total_LN[0]);
  tft.print(F(":"));
  tft.print(total_LN[1]);

  NickText(F("2"), x1, y2, 0);
  tft.print(F("-"));
  tft.print(total_LN[2]);
  tft.print(F(":"));
  tft.print(total_LN[3]);

  NickText(F("3"), x1, y3, 0);
  tft.print(F("-"));
  tft.print(total_LN[4]);
  tft.print(F(":"));
  tft.print(total_LN[5]);

  NickText(F("4"), x1, y4, 0);
  tft.print(F("-"));
  tft.print(total_LN[6]);
  tft.print(F(":"));
  tft.print(total_LN[7]);

  NickText(F("5"), x1, y5, 0);
  tft.print(F("-"));
  tft.print(total_LN[8]);
  tft.print(F(":"));
  tft.print(total_LN[9]);

  NickText(F("6"), x1, y6, 0);
  tft.print(F("-"));
  tft.print(total_LN[10]);
  tft.print(F(":"));
  tft.print(total_LN[11]);

  NickText(F("7"), x1, y7, 0);
  tft.print(F("-"));
  tft.print(total_LN[12]);
  tft.print(F(":"));
  tft.print(total_LN[13]);

  NickText(F("8"), x1, y8, 0);
  tft.print(F("-"));
  tft.print(total_LN[14]);
  tft.print(F(":"));
  tft.print(total_LN[15]);

  NickText(F("9"), x1, y9, 0);
  tft.print(F("-"));
  tft.print(total_LN[16]);
  tft.print(F(":"));
  tft.print(total_LN[17]);

  NickText(F("10"), x2, y1, 0);
  tft.print(F("-"));
  tft.print(total_LN[18]);
  tft.print(F(":"));
  tft.print(total_LN[19]);

  NickText(F("11"), x2, y2, 0);
  tft.print(F("-"));
  tft.print(total_LN[20]);
  tft.print(F(":"));
  tft.print(total_LN[21]);

  NickText(F("12"), x2, y3, 0);
  tft.print(F("-"));
  tft.print(total_LN[22]);
  tft.print(F(":"));
  tft.print(total_LN[23]);

  NickText(F("13"), x2, y4, 0);
  tft.print(F("-"));
  tft.print(total_LN[24]);
  tft.print(F(":"));
  tft.print(total_LN[25]);

  NickText(F("14"), x2, y5, 0);
  tft.print(F("-"));
  tft.print(total_LN[26]);
  tft.print(F(":"));
  tft.print(total_LN[27]);

  NickText(F("15"), x2, y6, 0);
  tft.print(F("-"));
  tft.print(total_LN[28]);
  tft.print(F(":"));
  tft.print(total_LN[29]);

  NickText(F("16"), x2, y7, 0);
  tft.print(F("-"));
  tft.print(total_LN[30]);
  tft.print(F(":"));
  tft.print(total_LN[31]);

  NickText(F("17"), x2, y8, 0);
  tft.print(F("-"));
  tft.print(total_LN[32]);
  tft.print(F(":"));
  tft.print(total_LN[33]);

  NickText(F("18"), x2, y9, 0);
  tft.print(F("-"));
  tft.print(total_LN[34]);
  tft.print(F(":"));
  tft.print(total_LN[35]);

  NickText_center("TO#:", 120, 251, 0);
  tft.print(timeouts);
}


//==========================================================================================

void display_time()
{
  if (show_time) {
    show_time = false;

    now = rtc.now();
    min_now = now.minute();
    tft.setFont(&FreeSansBold9pt7b);
    tft.setTextSize(1);
    tft.fillRect(0, 0, 240, 20, ILI9341_BLACK);
    tft.setCursor(0, 14);
    tft.setTextColor(ILI9341_WHITE, ILI9341_BLACK);
    tft.print(now.month());
    tft.print("/");
    tft.print(now.day());
    tft.print("/");
    tft.print(now.year());
    tft.setCursor(190, 14);
    tft.print(now.hour());
    tft.print(":");
    if (now.minute() < 10) {
      tft.print("0");
    }
    tft.print(now.minute());
  }

  now = rtc.now();
  if (min_now != now.minute()) {
    show_time = true;  //refresh page every minute
  }
}


//==========================================================================================

void drawBitmap(int16_t x, int16_t y, const uint8_t *bitmap, int16_t w, int16_t h, uint16_t color) {
  int16_t i, j, byteWidth = (w + 7) / 8;
  uint8_t byte;
  for (j = 0; j < h; j++) {
    for (i = 0; i < w; i++) {
      if (i & 7) byte <<= 1;
      else      byte   = pgm_read_byte(bitmap + j * byteWidth + i / 8);
      if (byte & 0x80) tft.drawPixel(x + i, y + j, color);
    }
  }
}

//======================================================================================

void GetNum(String prompt, int &Number) {
  // print a numeric keyboard and a prompt message
  // and get a numeric integer
  tft.fillScreen(ILI9341_RED);
  draw_BoxNButtons();// draw the virtual keyboard
  tft.setCursor(10, 35);
  tft.setTextSize(1);
  tft.setTextColor(ILI9341_WHITE);
  tft.setFont(&FreeSansBold12pt7b);
  tft.println(prompt); //print prompt
  tft.setFont(&FreeSansBold24pt7b);
  tft.setCursor(10, 110);
  tft.setTextSize(1);
  tft.setTextColor(ILI9341_WHITE);
  tft.println(Number); //update new value

  while (!ok) { // repeat until ok is presse
    if (ts.touched()) {
      TS_Point p = ts.getPoint(); wait(100);
      p.x = map(p.x, 0, 240, 0, 240);
      p.y = map(p.y, 0, 320, 0, 320);

      if (p.x < 60 && p.x > 0 && p.y > 260 && p.y < 320) {
        Number = 0;  //If Cancel button is pressed
        p = {};
      }
      if (p.x < 60 && p.x > 0 && p.y > 200 && p.y < 260) {
        Number = (Number * 10) + 5;  // button 1 is pressed
        p = {};
      }
      if (p.x < 60 && p.x > 0 && p.y > 140 && p.y < 200) {
        Number = (Number * 10) + 1;  // button 6 is pressed
        p = {};
      }

      if (p.x < 120 && p.x > 60 && p.y > 260 && p.y < 320) {
        Number = (Number * 10) + 9;  // button 0 is pressed
        p = {};
      }
      if (p.x < 120 && p.x > 60 && p.y > 200 && p.y < 260) {
        Number = (Number * 10) + 6;  // button 3 is pressed
        p = {};
      }
      if (p.x < 120 && p.x > 60 && p.y > 140 && p.y < 200) {
        Number = (Number * 10) + 2;  // button 7 is pressed
        p = {};
      }

      if (p.x < 180 && p.x > 120 && p.y > 260 && p.y < 320) {
        Number = (Number * 10) + 0;  // button 1 is pressed
        p = {};
      }
      if (p.x < 180 && p.x > 120 && p.y > 200 && p.y < 260) {
        Number = (Number * 10) + 7;  // button 4 is pressed
        p = {};
      }
      if (p.x < 180 && p.x > 120 && p.y > 140 && p.y < 200) {
        Number = (Number * 10) + 3;  // button 8 is pressed
        p = {};
      }

      if (p.x < 240 && p.x > 180 && p.y > 260 && p.y < 320) {
        ok = true;  // button ok is pressed
        p = {};
      }
      if (p.x < 240 && p.x > 180 && p.y > 200 && p.y < 260) {
        Number = (Number * 10) + 8;  // button 5 is pressed
        p = {};
      }
      if (p.x < 240 && p.x > 180 && p.y > 140 && p.y < 200) {
        Number = (Number * 10) + 4;  // button 9 is pressed
        p = {};
      }

      tft.fillRect(0, 60, 240, 80, ILI9341_RED);  //clear result box
      tft.setFont(&FreeSansBold24pt7b);
      tft.setCursor(10, 110);
      tft.setTextSize(1);
      tft.setTextColor(ILI9341_WHITE);
      tft.println(Number); //update new value
    }
  }
}

//======================================================================================

void draw_BoxNButtons() {
  String symbol[3][4] = {
    { "1", "2", "3", "4" },
    { "5", "6", "7", "8" },
    { "C", "9", "0", "OK" }
  };
  //Draw the Result Box
  tft.fillRect(0, 60, 240, 80, ILI9341_RED);

  //Draw keys
  tft.fillRect(0, 260, 60, 60, ILI9341_RED);
  tft.fillRect(0, 140, 240, 120, ILI9341_BLACK);
  tft.fillRect(60, 260, 120, 60, ILI9341_BLACK);
  tft.fillRect(180, 260, 60, 60, ILI9341_GREEN);

  //Draw Horizontal Lines
  for (int h = 140; h <= 320; h += 60)
    tft.drawFastHLine(0, h, 240, ILI9341_WHITE);

  //Draw Vertical Lines
  for (int v = 0; v <= 240; v += 60)
    tft.drawFastVLine(v, 140, 1800, ILI9341_WHITE);

  tft.setFont(&FreeSansBold18pt7b);

  //Display keypad lables
  for (int j = 0; j < 3; j++) {
    for (int i = 0; i < 4; i++) {
      if (j == 2 && i == 3) {
        tft.setTextColor(ILI9341_BLACK);// ok is more visible
        tft.setCursor(5 + (60 * i), 183 + (60 * j));
        tft.println(symbol[j][i]);
        return;
      }
      tft.setCursor(20 + (60 * i), 183 + (60 * j));
      tft.setTextSize(1);
      tft.setTextColor(ILI9341_WHITE);
      tft.println(symbol[j][i]);
    }
  }
}
```
