## Extended Date - LIQ HD Arduino Source Code for "LIQ HD (Lick Instance Quantifier Home cage Device): An open-source tool for recording undisturbed two-bottle drinking behavior in a home cage environment": SD_RTC_functions.ino

```
//==========================================================================================

void start_rtc()
{

  // connect to rtc
  Wire.begin();
  Wire.setClock(3400000);
  Wire.setWireTimeout();

  if (!rtc.begin()) {
    logfile.println(F("rtc failed"));
    DEBUG_PRINTLN(F("rtc failed"));
  }

  if (!rtc.initialized() || rtc.lostPower()) {
    DEBUG_PRINT("RTC syncing to computer clock.");
    // following line sets the RTC to the date & time this sketch was compiled
    rtc.adjust(DateTime(__DATE__, __TIME__));
  }
}


//======================================================================================

void resetWire()
{
  TWCR = 0;
  Wire.end();
  Wire.begin();
  Wire.setClock(3400000);
  Wire.setWireTimeout();
}

//==========================================================================================

void make_file()
{
  // fetch the time
  DateTime now;
  now = rtc.now();

  SdFile::dateTimeCallback(dateTime_);

  filename[0] = (now.year() / 10) % 10 + '0'; //To get 3rd digit from year()
  filename[1] = now.year() % 10 + '0'; //To get 4th digit from year()
  filename[2] = now.month() / 10 + '0'; //To get 1st digit from month()
  filename[3] = now.month() % 10 + '0'; //To get 2nd digit from month()
  filename[4] = now.day() / 10 + '0'; //To get 1st digit from day()
  filename[5] = now.day() % 10 + '0'; //To get 2nd digit from day()

  for (uint8_t i = 0; i < 100; i++) {
    filename[6] = i / 10 + '0';
    filename[7] = i % 10 + '0';
    if (! SD.exists(filename)) {
      // only open a new file if it doesn't exist
      logfile = SD.open(filename, FILE_WRITE);
      break;  // leave the loop!
    }
  }
  if (!logfile) {
    error(F("Could not create file"));
  }

  DEBUG_PRINT(F("Logging to: "));
  DEBUG_PRINTLN(filename);
  nowloggingto();
  logfile.flush();

  if (!logfile.println()) {
    error(F("Could not write file"));
  }

  // Write headers to file
  logfile.print(F("Experimental_side,millis,datetime,"));
  if (log_LN) {
    for (int i = 1; i <= 36; i++) {
      logfile.print("LickNumber");
      logfile.print(i);
      logfile.print(",");
    }
  }
  if (log_LD) {
    for (int i = 1; i <= 36; i++) {
      logfile.print("LickDuration");
      logfile.print(i);
      logfile.print(",");
    }
  }
  if (log_BN) {
    for (int i = 1; i <= 36; i++) {
      logfile.print("BoutNumber");
      logfile.print(i);
      logfile.print(",");
    }
  }
  if (log_BD) {
    for (int i = 1; i <= 36; i++) {
      logfile.print("BoutDuration");
      logfile.print(i);
      logfile.print(",");
    }
  }
  if (log_BLN) {
    for (int i = 1; i <= 36; i++) {
      logfile.print("BoutLickNumber");
      logfile.print(i);
      logfile.print(",");
    }
  }
  if (log_BLD) {
    for (int i = 1; i <= 36; i++) {
      logfile.print("BoutLickDuration");
      logfile.print(i);
      logfile.print(",");
    }
  }
  logfile.println();

  //echo to serial monitor
  DEBUG_PRINT(F("Experimental_side,millis,datetime,"));
  if (log_LN) {
    for (int i = 1; i <= 36; i++) {
      DEBUG_PRINT("LickNumber");
      DEBUG_PRINT(i);
      DEBUG_PRINT(",");
    }
  }
  if (log_LD) {
    for (int i = 1; i <= 36; i++) {
      DEBUG_PRINT("LickDuration");
      DEBUG_PRINT(i);
      DEBUG_PRINT(",");
    }
  }
  if (log_BN) {
    for (int i = 1; i <= 36; i++) {
      DEBUG_PRINT("BoutNumber");
      DEBUG_PRINT(i);
      DEBUG_PRINT(",");
    }
  }
  if (log_BD) {
    for (int i = 1; i <= 36; i++) {
      DEBUG_PRINT("BoutDuration");
      DEBUG_PRINT(i);
      DEBUG_PRINT(",");
    }
  }
  if (log_BD) {
    for (int i = 1; i <= 36; i++) {
      DEBUG_PRINT("BoutDuration");
      DEBUG_PRINT(i);
      DEBUG_PRINT(",");
    }
  }
  if (log_BD) {
    for (int i = 1; i <= 36; i++) {
      DEBUG_PRINT("BoutDuration");
      DEBUG_PRINT(i);
      DEBUG_PRINT(",");
    }
  }
  DEBUG_PRINTLN();

  syncTime = now.minute();
  SYNC_COUNTER = 0;
}

//==========================================================================================

void nowloggingto()
{
  tft.fillRect(0, 20, 240, 320, ILI9341_RED);
  tft.setFont(&FreeSansBold12pt7b);
  tft.setTextColor(ILI9341_WHITE);
  NickText_center("Logging to: ", 120, 120, 1);
  NickText_center(filename, 120, 150, 1);//show what file is being recorded to (filename)
  tft.setFont(&FreeSans12pt7b);
  NickText_center("Calibrating Sensors", 120, 200, 1);
  NickText_center("Stand Back!", 120, 230, 1);
  wait(2000);//wait 2000ms
}

//==========================================================================================
void logging_same_file()
{
  tft.fillRect(0, 20, 240, 320, ILI9341_RED);
  tft.setFont(&FreeSans12pt7b);
  tft.setTextColor(ILI9341_WHITE);
  NickText_center("Logging to same file: ", 120, 120, 1);
  NickText_center(filename, 120, 150, 1);//show what file is being recorded to (filename)
  NickText_center("Calibrating Sensors", 120, 200, 1);
  NickText_center("Stand Back!", 120, 230, 1);
  wait(2000);//wait 2000ms
}

//======================================================================================

void write_to_file()
{
  // fetch the time
  now = rtc.now();
  uint32_t m = millis();

  logfile.print(E_side); //log Experimental Side
  logfile.print(F(", "));
  logfile.print(m); // log milliseconds since starting
  logfile.print(F(", "));
  logfile.print('"'); // log time
  logfile.print(now.year(), DEC);
  logfile.print(F("/"));
  logfile.print(now.month(), DEC);
  logfile.print(F("/"));
  logfile.print(now.day(), DEC);
  logfile.print(F(" "));
  logfile.print(now.hour(), DEC);
  logfile.print(F(":"));
  if (now.minute() < 10) {
    logfile.print("0");
  }
  logfile.print(now.minute(), DEC);
  logfile.print(F(":"));
  if (now.second() < 10) {
    logfile.print("0");
  }
  logfile.print(now.second(), DEC);
  logfile.print('"');
  if (log_LN) {
    for (int k = 0; k < 36; k++) { //write data to file for each sipper
      logfile.print(F(", "));
      logfile.print(LickNumber[k]);
    }
  }
  if (log_LD) {
    for (int k = 0; k < 36; k++) { //write data to file for each sipper
      logfile.print(F(", "));
      logfile.print(LickDuration[k]);
    }
  }
  if (log_BN) {
    for (int k = 0; k < 36; k++) { //write data to file for each sipper
      logfile.print(F(", "));
      logfile.print(BoutNumber[k]);
    }
  }
  if (log_BD) {
    for (int k = 0; k < 36; k++) { //write data to file for each sipper
      logfile.print(F(", "));
      logfile.print(BoutDuration[k]);
    }
  }
  if (log_BLN) {
    for (int k = 0; k < 36; k++) { //write data to file for each sipper
      logfile.print(F(", "));
      logfile.print(BoutLickNumber[k]);
    }
  }
  if (log_BLD) {
    for (int k = 0; k < 36; k++) { //write data to file for each sipper
      logfile.print(F(", "));
      logfile.print(BoutLickDuration[k]);
    }
  }

  logfile.println();  // this is going to next line

  DEBUG_PRINT(E_side);
  DEBUG_PRINT(F(", "));
  DEBUG_PRINT(m);
  DEBUG_PRINT(F(", "));
  DEBUG_PRINT('"');
  DEBUG_PRINTDEC(now.year());
  DEBUG_PRINT(F("/"));
  DEBUG_PRINTDEC(now.month());
  DEBUG_PRINT(F("/"));
  DEBUG_PRINTDEC(now.day());
  DEBUG_PRINT(F(" "));
  DEBUG_PRINTDEC(now.hour());
  DEBUG_PRINT(F(":"));
  if (now.minute() < 10) {
    DEBUG_PRINT("0");
  }
  DEBUG_PRINTDEC(now.minute());
  DEBUG_PRINT(F(":"));
  if (now.second() < 10) {
    DEBUG_PRINT("0");
  }
  DEBUG_PRINTDEC(now.second());
  DEBUG_PRINT('"');
  if (log_LN) {
    for (int k = 0; k < 36; k++) {
      DEBUG_PRINT(F(", "));
      DEBUG_PRINT(LickNumber[k]);
    }
  }
  if (log_LD) {
    for (int k = 0; k < 36; k++) {
      DEBUG_PRINT(F(", "));
      DEBUG_PRINT(LickDuration[k]);
    }
  }
  if (log_BN) {
    for (int k = 0; k < 36; k++) {
      DEBUG_PRINT(F(", "));
      DEBUG_PRINT(BoutNumber[k]);
    }
  }
  if (log_BD) {
    for (int k = 0; k < 36; k++) {
      DEBUG_PRINT(F(", "));
      DEBUG_PRINT(BoutDuration[k]);
    }
  }
  if (log_BLN) {
    for (int k = 0; k < 36; k++) {
      DEBUG_PRINT(F(", "));
      DEBUG_PRINT(BoutLickNumber[k]);
    }
  }
  if (log_BLD) {
    for (int k = 0; k < 36; k++) {
      DEBUG_PRINT(F(", "));
      DEBUG_PRINT(BoutLickDuration[k]);
    }
  }
  DEBUG_PRINTLN();
  Serial.print("Time it took to write to file: ");
  Serial.print(millis() - m);
  Serial.println("ms");
}

//======================================================================================

void write_data()
{
  if (SYNC_COUNTER > SYNC_INTERVAL) {
    uint32_t m = millis();
    syncTime = now.minute();
    SYNC_COUNTER = 0;
    DEBUG_PRINT("Writing data to SD card!");
    digitalWrite(redLEDpin, HIGH); // blink LED to show we are syncing data to the card & updating FAT!
    logfile.flush();
    digitalWrite(redLEDpin, LOW);
    Serial.print("Time it took to sync to SD card: ");
    Serial.print(millis() - m);
    Serial.println("ms");
  }
}

//==========================================================================================

void dateTime_(uint16_t* date, uint16_t* Time)
{
  *date = FAT_DATE(now.year(), now.month(), now.day());
  *Time = FAT_TIME(now.hour(), now.minute(), now.second());
}
```
